# Breathing and tilting: mesoscale simulations illuminate influenza glycoprotein vulnerabilities

**DOI:** 10.1101/2022.08.02.502576

**Authors:** Lorenzo Casalino, Christian Seitz, Julia Lederhofer, Yaroslav Tsybovsky, Ian A. Wilson, Masaru Kanekiyo, Rommie E. Amaro

## Abstract

Influenza virus has resurfaced recently from inactivity during the early stages of the COVID-19 pandemic, raising serious concerns about the nature and magnitude of future epidemics. The main antigenic targets of influenza virus are two surface glycoproteins, hemagglutinin (HA) and neuraminidase (NA). Whereas the structural and dynamical properties of both glycoproteins have been studied previously, the understanding of their plasticity in the whole-virion context is fragmented. Here, we investigate the dynamics of influenza glycoproteins in a crowded protein environment through mesoscale all-atom molecular dynamics simulations of two evolutionary-linked glycosylated influenza A whole-virion models. Our simulations reveal and kinetically characterize three main molecular motions of influenza glycoproteins: NA head tilting, HA ectodomain tilting, and HA head breathing. The flexibility of HA and NA highlights antigenically relevant conformational states, as well as facilitates the characterization of a novel monoclonal antibody, derived from human convalescent plasma, that binds to the underside of the NA head. Our work provides previously unappreciated views on the dynamics of HA and NA, advancing the understanding of their interplay and suggesting possible strategies for the design of future vaccines and antivirals against influenza.

**One-Sentence Summary:** In situ dynamics of influenza glycoproteins expose antigenically relevant states and a new site of vulnerability in neuraminidase.

## Main Text

Influenza is a single-stranded, negative-sense RNA virus that contains two membrane-embedded glycoproteins: hemagglutinin (HA) and neuraminidase (NA) (*1–4*). The viral envelope is also dotted with transmembrane proton channels named M2 (*5, 6*). HA and NA are involved in a complex interplay whose combined functions include viral entry, progeny release, and evading host immune pressure (*7–9*). These processes are, in part, modulated by glycans, whose number and position can vary from year to year owing to the relentless evolution of the influenza virus (*10–13*). As a part of influenza seasonal antigenic drift, the introduced missense mutations may directly alter the antigenicity of HA (*10*) and NA (*13*) epitopes or sometimes result in the gain or loss of a glycosylation site (*10*), thus leading to shielding or unmasking of nearby epitopes. Antigenic drift means that vaccines targeting influenza need to be regularly updated to match the currently predominant strains of the virus (*14, 15*). Despite HA being the immunodominant protein (*16, 17*), NA also represents a potential vaccine target (*18*); knowledge of accessible and currently utilized epitopes is crucial in retaining influenza vaccine efficacy and in designing new vaccines. This is particularly relevant as the scientific community marches toward a universal influenza vaccine that is not affected by antigenic drift (*16, 17, 19*).

Antigenic drift also affects the HA/NA interplay, which in turn is relevant in the context of influenza endocytosis (*20*) and exocytosis (or viral budding) (*21–23*). The understanding of these processes is far from complete. As just one example, it was previously thought that NA played no role in endocytosis or exocytosis (*24*). Next, it was assumed that HA was involved in receptor binding and was not essential for viral budding (*25*), whereas NA was only involved in exocytosis by cleaving terminal host cell sialic acid moieties hydrogen bound to HA. More recently, it has been recognized that the role these two proteins play in cell entry and exit is more nuanced. NA can play a role in receptor binding (*26–28*) while HA can initiate viral budding (*21*). This egress process appears to be the rate-limiting step in viral replication, with only ∼10% of budded viral particles being fully released (*22*) and is the major target of current antivirals. Considering the weak binding capabilities of both HA and NA, multivalent interactions are needed for both cell entry and exit (*26, 29–32*). Although the exact valency for influenza cell entry and exit is still not known (*30*), sialic acid has been determined to be the cell receptor recruitment group involved in these multivalent interactions (*33–36*).

One potential reason for our limited understanding of the specifics of HA and NA function may be their poorly understood native dynamics *in vivo*. Whole virion protein dynamics are extremely challenging to study experimentally; current techniques such as surface affixation or aggregated kinetics assays are difficult to interpret due to their non-native environment or virion pleiomorphy, respectively. Cryo-electron tomography (cryoET) images can capture native virions, but protein flexibility diminishes the resolution of the resulting images. Instead, cryo-electron microscopy (cryoEM) has been successful in characterizing the plastic behavior of the individual glycoprotein ectodomains, showing, for example, the existence of tilted HA conformations (*37*), or “breathing” HA heads (*38–43*) and open NA heads (*44*). Some of the above-mentioned difficulties can be allayed through computational studies, which can create specific, pre-defined mesoscale models and examine protein dynamics in the mesoscale (*45–49*). Whole-virion scale simulations, whether coarse-grained (*50–52*) or all-atom (*45, 53–62*) can provide a detailed look at protein dynamics in a more realistic, crowded protein environment than simulations of discrete proteins. Additionally, such large simulations naturally afford statistical sampling for proteins of interest, i.e., an influenza virion will contain a few hundred HA proteins and a few dozen NA proteins, enabling the application of statistical analysis techniques, such as Markov state models (MSM), to quantify conformational transitions (*45, 63–65*).

Building on our previous work (*45*), we created two glycosylated whole-virion models accounting for two evolutionary-linked influenza A viruses, namely, A/swine/Shandong/N1/2009 (H1N1) (Fig. 1), hereafter referred to as H1N1-Shan2009, and A/45/Michigan/2015 (H1N1), hereafter referred to as H1N1-Mich2015 (fig. S1). We then performed all-atom MD simulations of these massive systems—tallying 161 million atoms including explicit water and ions—to examine the conformational plasticity of influenza glycoproteins in a crowded environment and its impact on their antigenic properties and dynamic interplay (Movie S1 and Movie S2).

**Fig. 1.**
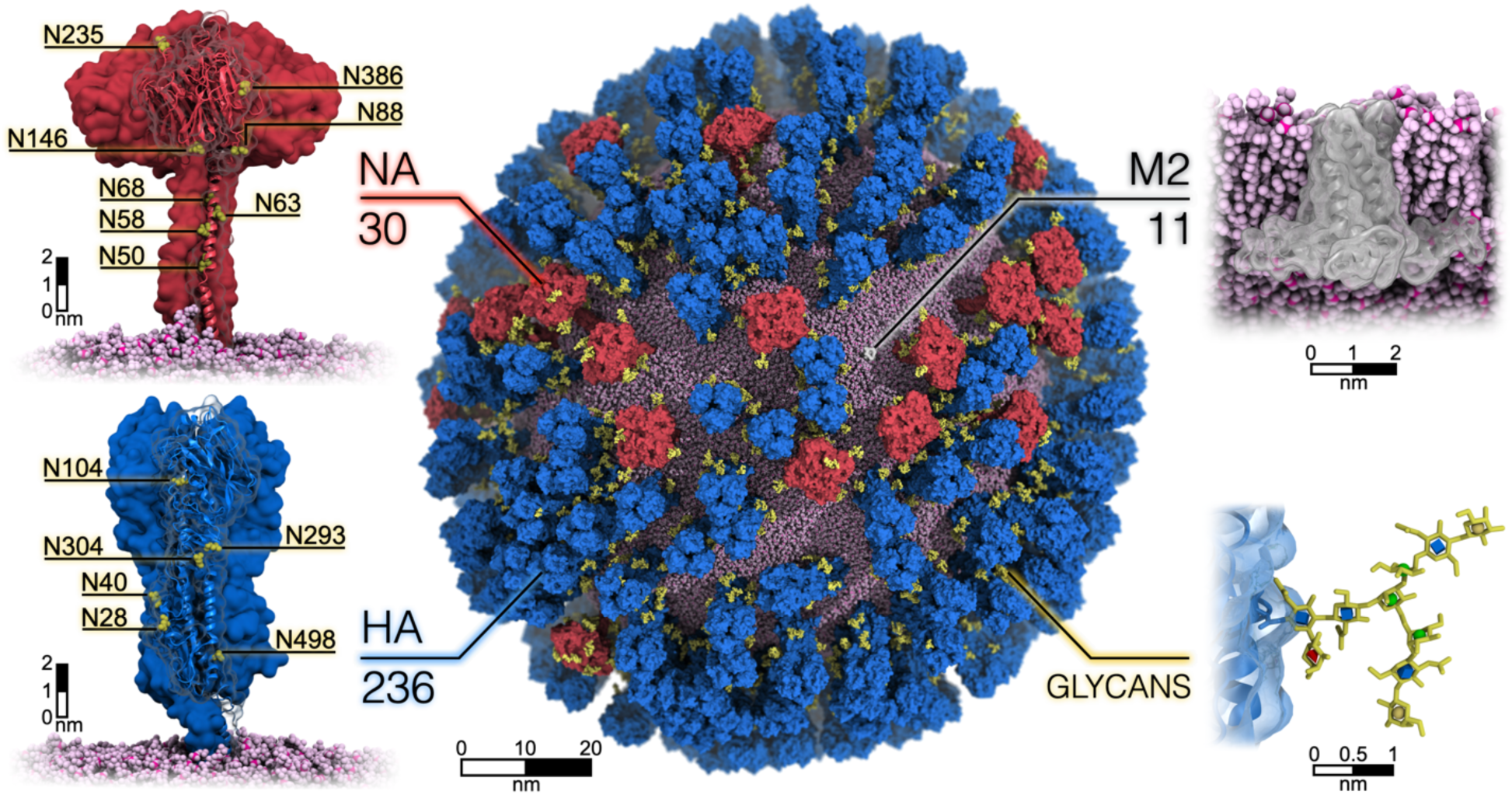
All-atom glycosylated model of influenza A virion. The center panel shows the all-atom model of the influenza H1N1-Shan2009 virion, where the neuraminidase (NA) and hemagglutinin (HA) glycoproteins, and the M2 ion channels, are depicted with red, blue, and white surfaces, respectively. N-linked glycans are shown in yellow. The spherical lipid envelope is represented with pink van der Waals (vdW) spheres. Magnified views of NA (top left) and HA (bottom left) are provided, where asparagine residues within each N-linked glycosylation sequon are shown with yellow vdW spheres. A cross-section, magnified view of an M2 ion channel is represented in the top-right corner. In the bottom right corner, a magnified view of a glycan is displayed, where the glycan is shown with yellow sticks, and the different monosaccharides are highlighted using 3D-SNFG representation (blue cube for GlcNAc, yellow sphere for galactose, red cone for fucose, and green spheres for mannose). Hydrogen atoms have been hidden for clarity.

As a result, our simulations indicate three main molecular motions underlying the exceeding flexibility of the influenza glycoproteins: NA head tilting, HA ectodomain tilting, and HA breathing. Using a Markov state model analysis framework, we characterize the kinetics of these motions. Concomitant with these three motions, the vulnerabilities of the glycoproteins are uncovered, including a novel epitope seen by a human monoclonal antibody and located on the underside of the NA globular head, which becomes directly accessible when NA head tilting occurs. In addition, we show how cryptic or usually occluded epitopes become transiently accessible during HA breathing and HA ectodomain tilting, respectively. Finally, we characterize the interplay between glycoproteins, breaking down the number and type of inter-glycoprotein contacts, and measuring the extent of glycoprotein “clumping” occurring on the virion surface. Although not many differences are observed between H1N1-Shan2009 and H1N1-Mich2015, the loss of glycan N386 within the NA head domain in the H1N1-Mich2015 remarkably reduces NA propensity to engage with neighboring glycoproteins.

Overall, our findings advance the understanding of HA/NA functional balance as affected by the antigenic drift, shedding light on the viral egress process. We propose that the synergy of the three identified motions is critical for the exocytosis cycle. Moreover, our work provides previously unappreciated views into the antigenicity of the influenza glycoproteins by accounting for their dynamics in the crowded environment of the virion surface. This information suggests that locking HA and NA in transiently sampled vulnerable states can be pursued as a viable strategy for the development of the next flu vaccines and antiviral drugs.

## Results

All-atom MD simulations of H1N1-Shan2009 and H1N1-Mich2015 glycosylated whole-virion models were conducted for ∼442 and ∼425 ns, respectively. Considering that our models comprise 236 HA trimers and 30 NA tetramers, an aggregate monomeric sampling of ∼313 and ∼53 μs were attained for H1N1-Shan2009, respectively, whereas ∼310 and ∼51 μs for H1N1-Mich2015, respectively. A detailed description of system setup, mutation of H1N1-Shan2009 into H1N1-Mich2015, glycosylation process, and MD simulations are provided in the Supplementary Information (SI) Material and Methods section and shown in Figs. S2-S8. Analysis of the average Root Mean Square Deviation (RMSD) values calculated for HA and NA evinces that the conformation of both glycoproteins keeps changing during the simulations with respect to the initial geometry (Fig. S9). More stability is attained only towards the end, where 4-7 Å and 4-11 Å RMSD ranges are observed for HA and NA, respectively. This behavior underlies the presence of dynamic interplay among the glycoproteins during the simulations facilitated by the remarkable flexibility of the glycoprotein ectodomains.

### Extensive tilt exposes neuraminidase head underside to immune recognition

The NA globular head sits on top of the stalk and is formed by six β-sheets arranged into a propeller-like structure (*66*). The first β-sheet is connected to the stalk helix by an unstructured, 14-residue-long linkage (residues 82–95) that acts as a hinge point and shows to be highly flexible during the simulations, as evinced by the Root Mean Square Fluctuations (RMSF) analysis (Fig. S10). In the tetrameric arrangement, the impact of this structural feature is two-fold: uncoupling the motions of the head and the stalk and augmenting the head’s conformational freedom. Our whole-virion simulations show that the NA head undergoes an extensive tilt motion, exhibiting extraordinary flexibility (Fig. 2A-B, Movie S3, Movie S4).

**Fig. 2.**
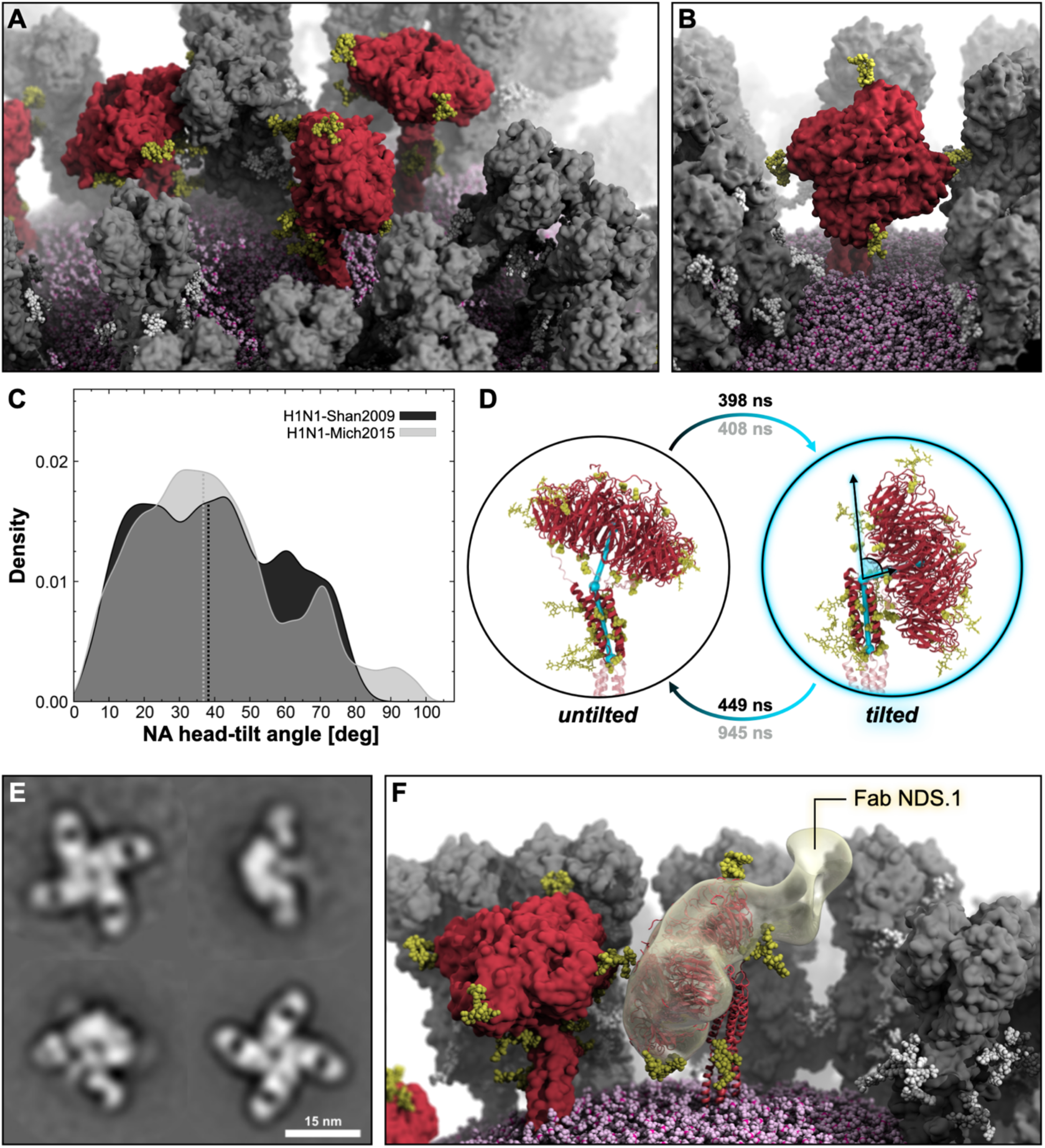
NA head-tilt motion. **A**) A snapshot from the MD simulation of the influenza H1N1-Shan2009 virion capturing different NAs (red surface) with varying degrees of head tilt. HAs are represented with a gray surface, whereas N-glycans linked to NA or HA are shown with yellow or white vdW spheres, respectively. The lipid envelope is depicted using pink vdW spheres. **B**) Close-up view of a NA tetramer exhibiting a high degree of head tilt. **C**) Distribution of NA head-tilt angle obtained from the MD simulation of the H1N1-Shan2009 (black) and H1N1-Mich2015 (gray) virions. The distribution is shown as a kernel density. **D**) Two-state MSM of NA head-tilt with representative structures from the MD simulation of the H1N1-Shan2009 virion. The mean first-passage times between the states are shown for H1N1-Shan2009 (black) and H1N1-Mich2015 (gray). NA head-tilt angle is highlighted with cyan cylinders, spheres, and lines drawn within the molecular representation of the NA. NA residues used for angle calculation are shown with opaque red cartoons. N-glycans are represented with yellow sticks, whereas glycosylation sequons are illustrated with yellow vdW spheres. **E)** NS-EM 2D class averages of recombinant A/Darwin/9/2021 N2 in complex with Fab NDS.1 (scale bar: 15 nm) **F**) The density of a single NDS.1 Fab bound to A/Darwin/9/2021 N2 is fitted onto a tilted NA head (red cartoons) as captured in a snapshot from the H1N1-Shan2009 whole-virion simulation. The density is shown with a transparent yellow surface. The surrounding HA and NA are represented in gray and red surfaces, respectively, whereas N-glycans are depicted with yellow vdW spheres.

Remarkably, the NA head can tilt >90° relative to the stalk axis, as shown in Fig. 2B. We calculated the tilt angle formed by the stalk’s principal axis and the vector joining the top of the stalk with the center of mass (COM) of the tetrameric head for all 30 NA tetramers of the two virion models, at each frame of their respective simulations. Next, we plotted the two sets of angle values as different distributions (Fig. 2C). The tilt angle is displayed in Fig. 2D. Interestingly, despite the differences in amino acid sequences, NA tetramers in H1N1-Shan2009 and H1N1-Mich2015 exhibit a similar extent of head tilt, with median values at 38.6° and 37.2°, respectively. A one-to-one correspondence between the two strains is also observed, with a few exceptions, for the maximum head tilt achieved by each NA tetramer during the simulations (Fig. S11). Values of head tilt slightly larger than 90° are allowed by the flexibility of the hinge region between the head and the stalk. As the NA tetramer head tilts, at least one of the four monomer heads will move closer to the stalk. Incorporating this geometric criterion into a set of two features (see Material and Methods in the SI, and Fig. S12-S13) and exploiting the large cumulative sampling generated by the many NA tetramers present on the virion surface, we built a two-state MSM to characterize the kinetics of the NA head-tilt motion (Fig. 2D). As a result, for H1N1-Shan2009, the stationary distributions for the untilted and tilted states, which provide an understanding of the weight of each state in the MSM, are 0.51 and 0.49, respectively. The average times required for the transitions between the two states to first occur, referred to as mean first-passage times (MFPTs), are comparable: 398 ns for the untilted-to-tilted motion and 449 ns for the opposite, tilted-to-untilted transition. Instead, for H1N1-Mich2015, the stationary distributions are 0.44 (untilted) and 0.56 (tilted), with MFPTs of 408 ns (untilted-to-tilted) and 945 ns (tilted-to-untilted). These findings evince a similar propensity to tilt for the NA heads of both strains, showing that they can tilt >60° within hundreds of nanoseconds, whereas it takes slightly longer for them to return to their upright state (see Table S1 for the tilt angles calculated for the macrostates extracted from the MSM).

Aiming to investigate the immunogenic implications of the NA head-tilt motion, we studied an H3N2 influenza convalescent donor and discovered a human monoclonal antibody, termed NDS.1, which recognizes the underside of the NA head. NA-specific B cells were isolated from peripheral blood mononuclear cells from the donor using recombinant N2 NA head by flow cytometry (Fig. S14), while the recombinant antibody NDS.1 was synthesized by using immunoglobulin sequences recovered from the corresponding B cell. Biolayer interferometry (BLI) assays show that Fab of NDS.1 antibody possesses relatively high affinity (10^-8^ M) to two distinct N2 NA tetramers tested, namely A/Wisconsin/672005 and A/Darwin/9/2021 (Fig. S14). Negative-stain electron microscopy (NS-EM) analysis and its 3D reconstruction reveal that NDS.1 Fab recognizes an epitope residing on the underside of the NA head (Fig. 2E, Fig. S15). This was further confirmed by analyzing ternary complex of N2 NA of NDS.1 Fab and 1G01 Fab by NS-EM (Fig. S16), a recently discovered broadly cross-reactive antibody that binds the catalytic site of NA (*67*). We then fitted the conformation of one tilted NA tetramer head obtained from the H1N1-Shan2009 whole-virion simulation into the density map of the A/Darwin/9/2021 N2 NA tetramer head bound to a single NDS.1 Fab. As shown in Fig. 2F, the NA head underside region targeted by NDS.1 becomes directly available for immune recognition as the head tilts. Although the NA head underside is solvent accessible in the untilted state, it is not a facile target since the antibody and/or B cell receptors on B cells must reach this region from underneath with a slightly upward approach angle. Instead, the tilt exposes the head underside of at least one monomer and changes the antibody approach angle to downward, thus allowing NDS.1 Fab to gain direct access to the underside epitope. As evinced from the fitted density depicted in Fig. 2F, the binding of NDS.1 appears sterically compatible with the N-glycans linked to the (N1) NA head, i.e., N88, N146, N235, and N386 (for H1N1-Shan2009).

### Hemagglutinin ectodomain tilt facilitates the approach of anchor epitope-directed antibodies

The HA ectodomain protrudes ∼15 nm from the viral membrane, to which it is anchored via a coiled-coil transmembrane domain (TMD) (*37*). As also indicated by the average RMSF for HA (Fig. S10), a flexible hinge region encompassing residues 504–528 connects the HA ectodomain to the TMD, increasing the plasticity of the ectodomain itself (*37*). Our simulations show considerable flexibility of the HA ectodomain, which exhibits a profuse tilting motion relative to the TMD axis (Fig. 3A and Fig. 3B, Movie S5 and Movie S6). We measured the tilt angle formed by the TMD principal axis and the vector joining the residues at the top of the TMD and the center-of-mass (COM) of the ectodomain’s long α-helices (LαHs) for all 236 HA trimers from the two virion models, at each frame of their respective simulations. The distributions for the ectodomain-tilt angles are presented in Fig. 3C, whereas the calculated tilt angle is depicted in Fig. 3D. The distribution profiles obtained for H1N1-Shan2009 and H1N1-Mich2015 are very similar in shape, with median values of 22.6° and 22.3°, respectively. We note that the calculated tilt angle does not always coincide with the same extent of tilting relative to the membrane normal since the TMD also moves and tilts within the lipid bilayer. Remarkably, a few HA trimers display ectodomain tilt angle values larger than 50° over the course of the simulations (Fig. 3C). These values are in striking agreement with cryoEM experiments examining full-length HA in detergent micelles, where an ectodomain tilt of 52° was reported (*37*).

**Fig. 3.**
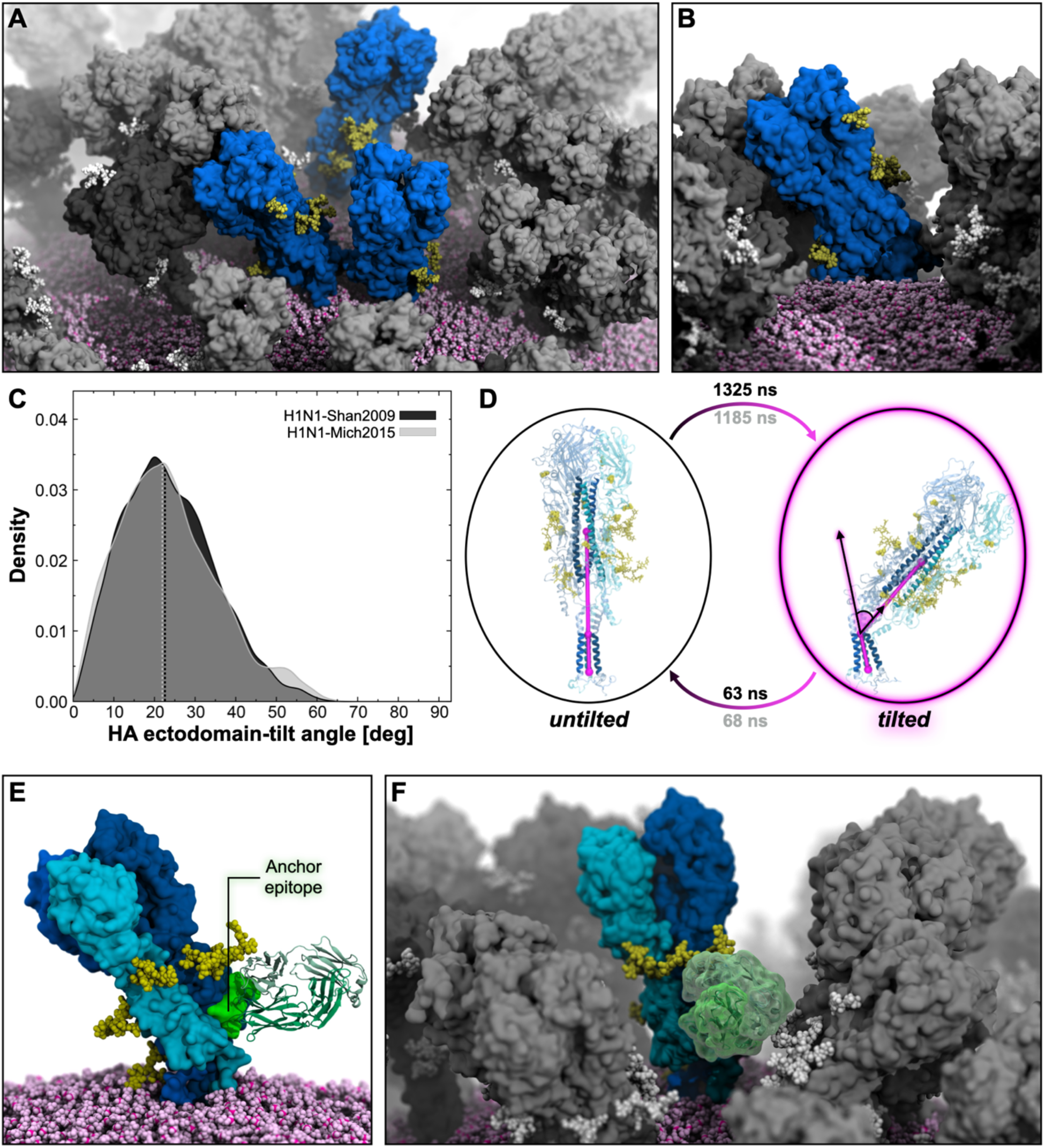
HA ectodomain-tilt motion. **A**) A snapshot from the MD simulation of the influenza H1N1-Shan2009 virion capturing different HAs with varying degrees of ectodomain tilt (blue surface). NAs are represented with a dark gray surface, while other HAs are represented with a gray surface. N-glycans linked to HAs or NAs are shown with yellow or white vdW spheres, respectively. The lipid envelope is depicted using pink vdW spheres. **B**) Close-up view of an HA trimer exhibiting a high degree of ectodomain tilt. **C**) Distribution of HA ectodomain-tilt angle obtained from the MD simulation of the H1N1-Shan2009 (black) and H1N1-Mich2015 (gray) virions. The distribution is shown as a kernel density. **D**) Two-state MSM of HA ectodomain tilt with representative structures from the MD simulation of the H1N1-Shan2009) virion. The mean first-passage times between the states are shown for H1N1-Shan2009 (black) and H1N1-Mich2015 (gray). HA ectodomain-tilt angle is highlighted with magenta cylinders, spheres, and lines drawn within the molecular representation of the HA. HA residues used for angle calculation are shown with opaque blue cartoons. N-glycans are represented with yellow sticks, whereas glycosylation sequons are illustrated with yellow vdW spheres. **E**) Molecular representation (side view) of a tilted HA trimer exposing one “anchor” epitope, highlighted with a shiny green surface, to FISW84 Fab (green cartoons) for direct binding (*37*). HA protomers are represented with surfaces colored with different shades of blue, whereas N-linked glycans are shown with yellow vdW spheres. The viral membrane is depicted with pink vdW spheres. **F**) Molecular representation (frontal view) of FISW84 Fab (green transparent surface) bound to the same HA trimer shown in panel E but in the context of the crowded environment of the H1N1-Shan2009 virion, where other HAs are represented with a gray surface, NAs with a dark gray surface, and N-linked glycans with white vdW spheres.

To gauge the transition time scales between the untilted and the tilted states, we built a two-state MSM of the HA ectodomain tilt using a combination of head–stalk distance criteria as features (Fig. 3D) (see Extended Material and Methods in the SI and Fig. S17-S18). The resulting stationary distributions are strongly shifted toward the untilted state, with values of 0.94 (untilted) and 0.06 (tilted) for both H1N1-Shan2009 and H1N1-Mich2009 (see Table S2 for the tilt angles calculated for the macrostates extracted with MSM). The MFPTs are also comparable between the two strains: 1325 ns (H1N1-Shan2009) and 1185 ns (H1N1-Mich2015) for the untilted-to-tilted transition; 63 ns (H1N1-Shan2009) and 68 ns (H1N1-Mich2015) for the tilted-to-untilted transition. This analysis reveals that, on average, longer time scales are needed by the ectodomain to considerably tilt (> 50°) relative to the TMD, whereas the untilted position is restored much faster.

The gradual tilting of the ectodomain has a profound impact on the accessibility of the so-called “anchor” epitope located on the lower portion of the HA stalk in the proximity of the viral membrane. As this epitope is known as a target for broadly neutralizing antibodies (*37, 68*), we docked the anchor-binding FISW84 Fab (*37*) onto both an untilted and a tilted conformation assumed by an HA trimer throughout the virion simulations, examining the respective poses. When the HA is untilted and upright, the antibody must approach the anchor epitope with an upward angle (Fig. S19). Due to the trimer symmetry, accessibility to all three epitopes is hampered by the presence of the lipid bilayer underneath (Fig. S18). On the contrary, when HA flexes onto the membrane, the angle of approach to at least one of the three anchor epitopes on the HA trimer gradually shifts from upward to lateral or slightly downward (Fig. 3E, Fig S19), facilitating antibody binding. Next, we docked the FISW84 Fab fragment (*37*) onto a tilted HA in the context of the crowded virion environment. As shown in Fig. 3F, the antibody can directly target the anchor epitope, even in the presence of neighboring glycoproteins.

### Breathing of hemagglutinin head reveals a cryptic epitope

The immunodominant domain of HA is the globular head (*39, 69*), spanning residues 66 to 286 and comprising the receptor-binding site (RBS) and the vestigial esterase domain (*70*). In the trimer assembly, the HA1 subunits and the respective head domains are tightly packed in a “closed” state around and atop the HA2 LαHs. However, upon binding to sialylated receptors during the infection process, HA heads, followed by the HA1 subunits, swing away from HA2, making room for the necessary conformational changes leading to membrane fusion (*71*). We note that in this work HA was modeled in its uncleaved form (HA0). In our whole-virion constructs, all of the HA trimers are initially in the closed state, with the heads interacting with each other atop the LαHs. However, over the course of the virion simulations, some of the HA trimers start “breathing,” i.e., reversibly transitioning from the closed state to a partially “open” state where the heads transiently move away from each other (Fig. 4A–B, Movie S7 and Movie S8).

**Fig. 4.**
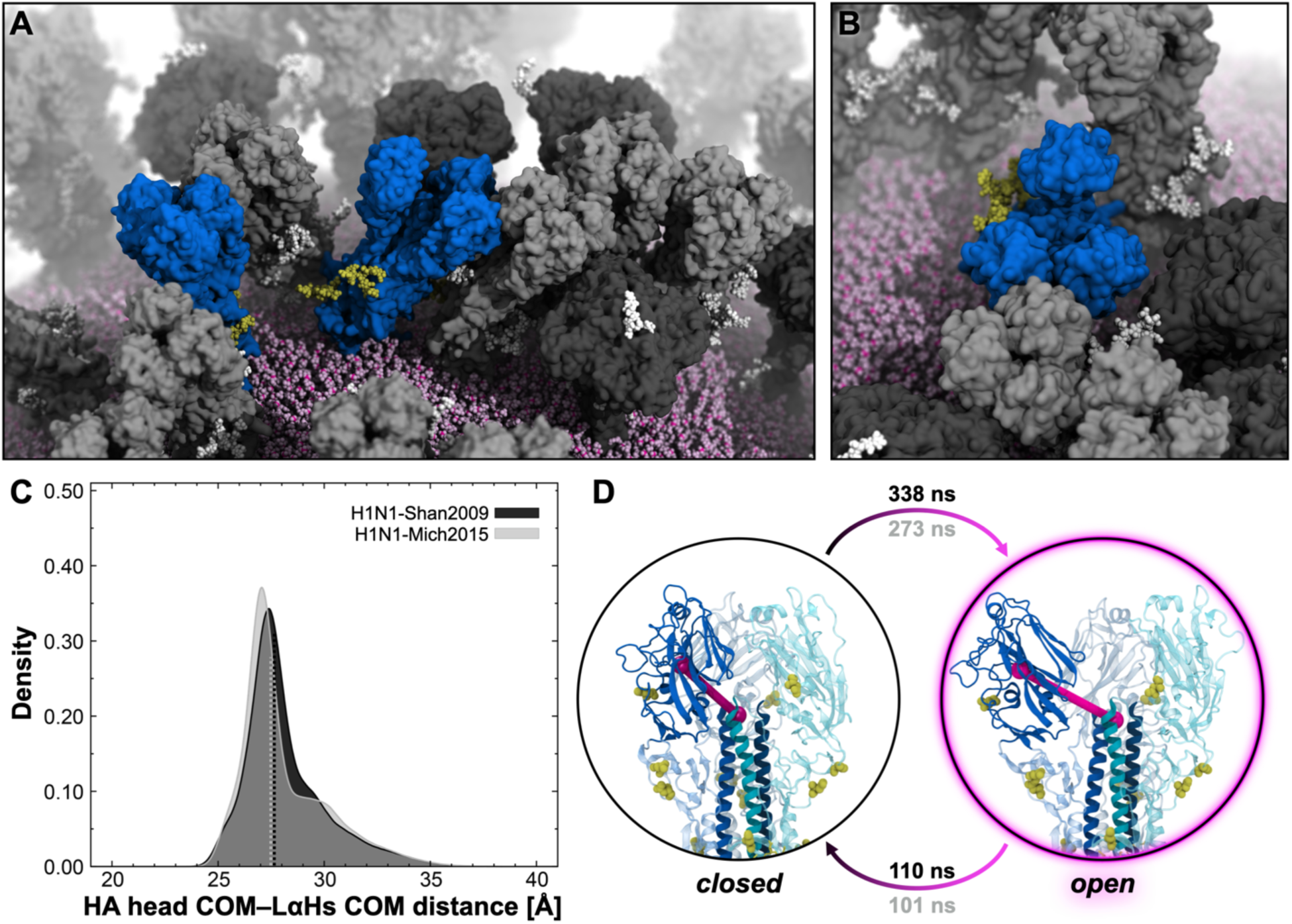
Breathing motion of the HA head domain. **A**) A snapshot from the MD simulation of the H1N1-Shan2009 virion capturing different HAs with varying degrees of head breathing motions (blue surface). NAs are represented with a dark gray surface, whereas other HAs are represented with a gray surface. N-glycans linked to HAs or NAs are shown with yellow or white vdW spheres, respectively. The lipid envelope is depicted using pink vdW spheres. **B**) Close-up, top view of an open HA trimer where the head domains undergo extensive breathing motion. **C**) Value distribution of the distance between the center-of-mass (COM) of the HA head and the COM of the long α-helices’ apical residues (LαHs) obtained from the MD simulation of the H1N1-Shan2009 (black) and H1N1-Mich2015 (gray) virions. The distribution is shown as a kernel density. Dashed lines indicate the median values of the respective distribution. **D**) Two-state MSM of HA head-breathing motion with representative structures from the MD simulation of the H1N1-Shan2009 virion. The mean first-passage times between the states are shown for H1N1-Shan2009 (black) and H1N1-Mich2015 (gray). Monitored HA head–LαHs distance is depicted with magenta cylinders drawn within the molecular representation of the HA. HA residues of the head domain and LαHs are shown with opaque blue cartoons. N-glycans are represented with yellow sticks, whereas glycosylation sequons are illustrated with yellow vdW spheres.

We examined the extent of the HA head breathing motion for all the 236 HA trimers of the two virion models by measuring the distance between the COM of each monomeric HA head and the COM of the apical residues of the three LαHs. The distributions of the calculated values are presented in Fig. 4C, whereas the monitored distance is illustrated in Fig. 4D. Analogously to the NA head-tilt and HA ectodomain-tilt motions, H1N1-Shan2009 and H1N1-Mich2015 display a similar extent of HA breathing, exhibiting almost overlapping right-skewed distributions with median values of 27.5 Å and 27.7 Å, respectively (Fig. 4C). These values coincide with a closed state. When the HA head starts breathing, its distance from the LαHs becomes larger than 30 Å, reaching maximum values of 41.0 Å and 42.2 Å for H1N1-Shan2009 and H1N1-Mich2015, respectively.

The two-state MSM constructed for the HA head breathing motion allows us to quantify the kinetics of the closed-to-open and open-to-closed transitions. In this case, we used one feature, defined as the smallest mean distance of the HA heads to each other (see Material and Methods in the SI, and Fig. S20-S21). For H1N1-Shan2009, a stationary distribution of 0.75 was estimated for the closed state, whereas 0.25 for the open state. For H1N1-Mich205, a stationary distribution of 0.73 was calculated for the closed state, while 0.27 for the open state. The MFPTs for the closed-to-open transition are 338 ns (H1N1-Shan2009) and 273 ns (H1N1-Mich2015), whereas MFPTs of 110 ns (H1N1-Shan2009) and 101 ns (H1N1-Mich2015) were computed for the open-to-closed transition. The opening motion is slower on average than the return to a closed conformation, which is almost three times faster, pinpointing the open state as a short-lived state. The values of the distance between the HA head and the LαHs for the macrostates extracted with MSM are reported in Table S3.

Recently, a novel, extremely conserved epitope within the HA head was discovered (*39*), providing further opportunities for the development of a universal vaccine against influenza (*72*). Promisingly, FluA-20, a broadly protective human antibody, was found to target this cryptic epitope in the head interface region with high breadth and potency against many influenza A virus subtypes (*39*). Similar to a previously discovered epitope buried in the HA head trimer interface (*73*), the FluA-20 epitope is completely occluded when the HA is in the closed state (Fig. S22), necessitating an opening, or breathing, of the HA head to become accessible (*39*). Our simulations show that the FluA-20 cryptic epitope becomes transiently accessible during HA head breathing (Fig. 5A–B). To characterize whether the HA head opening is compatible with FluA-20 binding to the cryptic epitope, we extracted the most open conformation of one HA trimer from the H1N1-Shan2009 simulation, and we docked FluA-20 onto it. Strikingly, FluA-20 could be fitted onto the cryptic epitope with no apparent clashes with neighboring monomers (Fig. 5A and Fig. S22). This pose is similar to the one disclosed in a recent structural study (*43*). Interestingly, the FluA-20 binding to the open HA head is also compatible in the crowded virion environment (Fig. 5B). We next estimated the accessibility of the cryptic epitope to FluA-20. FluA-20 variable domains can be fitted into a sphere of radius 18.6 Å (Fig. 5C), which we used as a probe to calculate the antibody accessible surface area (AbASA) of the cryptic epitope during the simulation (Fig. 5D). As a test case, we considered an HA trimer that showed extensive breathing. Together with the AbASA profile, we also present the time evolution profile of the distance between the three HA head COMs and the LαHs COM (Fig. 5E).

**Fig. 5.**
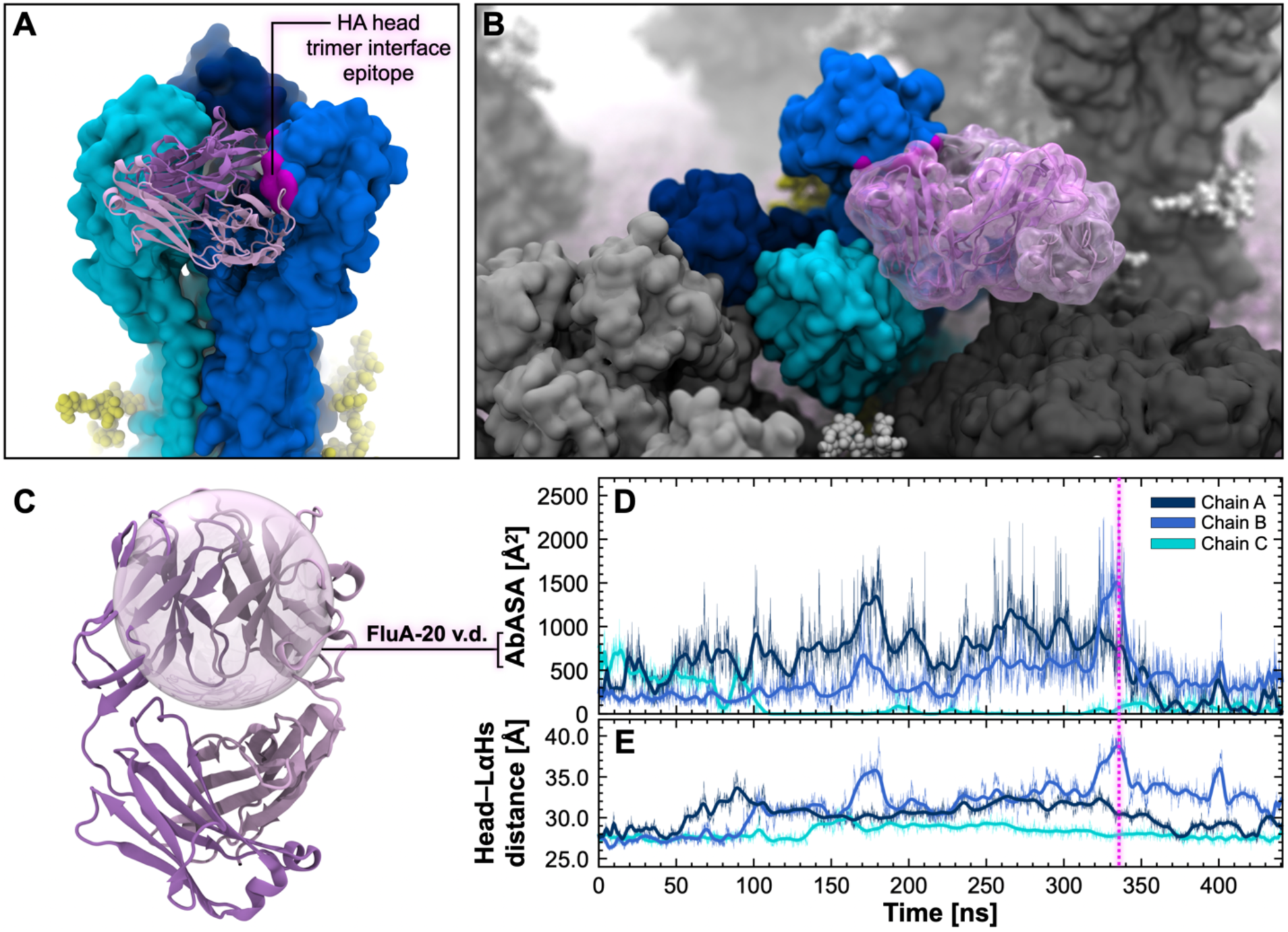
HA breathing reveals the FluA-20 cryptic epitope. **A**) Molecular representation of an HA trimer in the open state extracted from the H1N1-Shan2009 simulations (side view). The three HA monomers are depicted with surfaces colored with different shades of blue. FluA-20 heavy and light chains are represented with pink and purple cartoons, respectively. The cryptic epitope targeted by FluA-20 is highlighted with a magenta surface. N-linked glycans are shown with yellow vdW spheres. **B**) Molecular representation of the same open HA trimer shown in panel A, in the context of the crowded virion environment. NAs are represented with a dark gray surface, whereas other HAs with a gray surface. N-glycans linked to HAs or NAs are shown with yellow or white vdW spheres, respectively. The lipid envelope is depicted using pink vdW spheres. **B**) Close-up view of an open HA trimer where the head domains undergo extensive breathing motion. **C**) Molecular representation of the FluA-20 human antibody, where the heavy and light chains are depicted with pink and purple cartoons, respectively. The 18.6 Å radius probe sphere comprising FluA-20 variable domains is overlaid as a transparent surface. **D**) The time evolution profile of the AbASA of the FluA-20 epitope is shown for the three HA chains illustrated in panels A and B. A magenta dashed line indicates the frame shown in panel A and B. **E**) The time evolution profile of the distance between the COM of the HA head and the COM of the LαHs’ apical residues is shown for the three HA chains illustrated in panels A and B.

The respective epitopes in chains A, B, and C in the trimeric HA head show different accessibility according to the extent of breathing. Chains A and B in the head display the most extensive breathing, and the maximum epitope accessibility occurs in correspondence to the full opening of chain B in the head (Fig. 5D–E). The snapshot corresponding to the maximum head opening of chain B was used to dock FluA-20 onto the respective epitope (Fig. 5A–B). Another interesting consideration is that the heads appear to breathe asymmetrically, opening or closing irrespective of each other (Fig. 5E). They can fleetingly open and suddenly rotate back to a close position. Interestingly, possibly due to the presence of nearby glycoproteins, the head of chain C does not appear to breathe, keeping the respective FluA-20 cryptic epitope fully occluded.

### The dynamic interplay between glycoproteins

Throughout the simulation of both virus strains, HA and NA are characterized by exceptional flexibility of their extra-virion functional domains. Among others, we have highlighted the NA head-tilt motion, the HA ectodomain-tilt motion, and the breathing of HA heads. Although these motions relate to individual glycoproteins, they occur in a crowded subcellular environment that is realistically reproduced in our MD simulations with an atomic level of detail. This is one of the main advantages of performing simulations at the “whole-virion” regime rather than at the “single-protein” regime. Embedded in the viral membrane are, in fact, 236 HA trimers and 30 NA tetramers that do more than wiggling and jiggling and whose interplay and balance are at the basis of viral fitness (*74*). We, therefore, examined the intricate yet fundamental interplay between HA and NA glycoproteins in our mesoscale simulations. The glycoproteins can move across the lipid bilayer and can twist, bend, tilt and interact with each other using their extra-virion functional domains. Even though transmembrane proteins in model membranes, such as ours, usually partition into unrealistic liquid disordered domains (*75*), we see protein clustering, which is also seen biologically (*76, 77*). As proteins usually do not form these sorts of clusters, or quinary interactions, without a purpose (*78, 79*), these patches are then likely to have functional significance in viral entry and/or egress (*76, 80, 81*). To investigate the motion of the glycoproteins’ TMD within the viral membrane, we measured the RMSF of the TMD COM of every HA and NA over the course of the simulations with respect to the starting frame. Similarly, we also computed the RMSF of the ectodomain COM of every HA and NA. We then compared the resulting distributions of RMSF values (Fig. S22). For both strains, this analysis evinces that HA and NA do not vastly translate through the membrane in the microsecond timescales, whereas their ectodomains show higher mobility than the TMDs since they can tilt or breathe (Fig. S23). However, we cannot rule out the possibility that HA and NA could translate more over longer timescales. With this regard, we note that the interior of the virion models simulated here is filled with water molecules and that the matrix (M1) proteins lining the internal side of the POPC lipid bilayer were not modeled.

Next, we analyzed the number of one-to-one connections established by each glycoprotein with the surrounding glycoproteins. A connection is formed when two glycoproteins move within 5 Å of each other considering only their protein/glycan heavy atoms (see Material and Methods in the SI for additional details). The schematic and the analysis are shown in Fig. 6A. We found that each glycoprotein can connect with up to five other glycoproteins at once. Interestingly, the fraction of isolated glycoproteins (i.e., glycoproteins without a connection to any other glycoprotein) decreased over the course of the simulations from 34.2% to 12.4% in H1N1-Shan2009 and from 36.8% to 16.5% in H1N1-Mich2009. The glycoproteins tend to interact with each other forming new connections rapidly (Movie S9). The most considerable growth is in the fraction of glycoproteins forming two connections, 17.3% to 34.2 % (H1N1-Shan2009) and 16.2% to 34.2 (H1N1-Mich2015), followed by the fraction of glycoproteins forming three connections, 5.6% to 13.5% (H1N1-Shan2009) and 4.1% to 11.3% (H1N1-Mich2015). More rarely, HA and NA interconnect with four or five glycoproteins simultaneously. Considering that few quinary interactions last into the µs timescale (*82*) our results suggest dynamic fluctuations in the degree of protein clumping: within just one microsecond, large protein clumps can form and dissipate. Despite the point mutations and the glycan addition/deletion within HA/NA, the overall glycoprotein interplay has not dramatically changed from H1N1-Shan2009 to H1N1-Mich2015. However, appreciable differences are observed in the case of NA in H1N1-Mich2015, where the loss of glycan N386 located at the edge of the head appears to have slowed down its propensity to form new connections (Fig. S24). The dynamic evolution of the pattern of glycoprotein inter-connections is shown in Movie S9 for H1N1-Shan2009 and H1N1-Mich2015.

**Figure 6.**
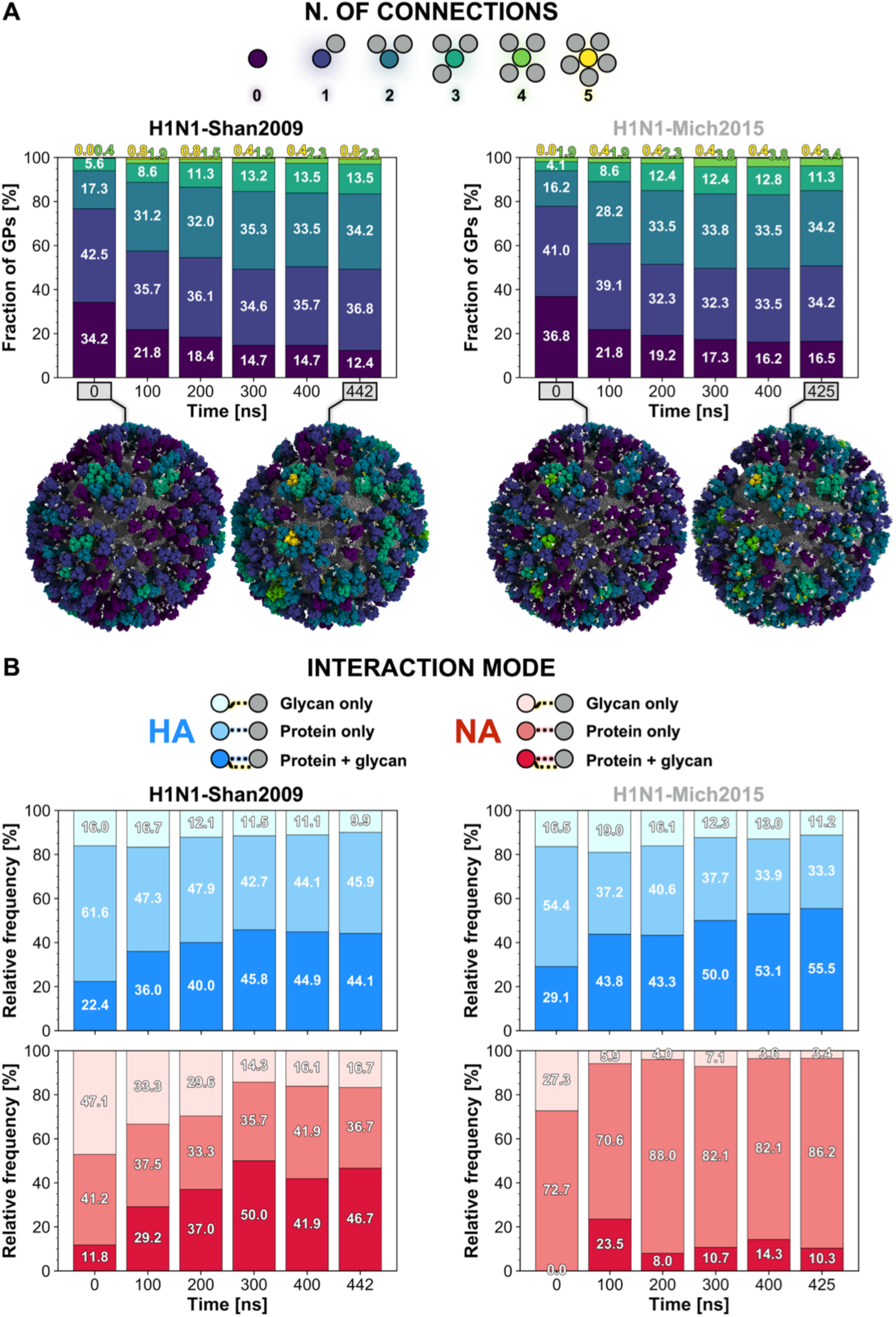
HA/NA interplay. **A)** The pattern of connections (0 to 5) made by either HA or NA (circles colored in viridis palette) with surrounding glycoproteins (gray circles) is represented in the schematic shown on top of the panel. Below are the stacked bar plots representing the fraction (%) of glycoproteins establishing a certain number of connections with surrounding glycoproteins along the simulation of H1N1-Shan2009 (left) and H1N1-Mich2015 (right). Percentages rounded to the first decimal digit are reported. Molecular representations of the virions at the initial and last frame of the simulations are shown under the plots. The glycoproteins are depicted with a surface colored according to the number of connections made by the respective glycoprotein at that frame. Interacting glycans are highlighted with yellow vdW spheres, non-interacting glycans with white vdW spheres. **B)** The legend for the interaction mode between glycoproteins (either HA or NA) is schematically represented on top of the panel. Below are the stacked bar plots representing the relative frequency (%) of connections made by the glycoproteins and involving glycan only (lighter shade), protein only or protein and glycan (darker shade) simultaneously along the simulation of H1N1-Shan2009 (left) and H1N1-Mich2015 (right). A blue gradient for the three categories is adopted for HA, whereas a red gradient is used for NA. Percentages rounded to the first decimal digit are reported.

A consequence of influenza glycoproteins clumping together is the formation of large macro-clusters of mixed HA/NA when multiple small clusters come into close contact, i.e., less than 5 Å from each other (Fig. S25). Congregations of HAs and local clusters of NAs have been previously reported through cryoET experiments (*83*). Here, we observed the formation of macro-clusters comprising as many as 42 (H1N1-Shan2009) and 34 (H1N1-Mich2015) mixed HA/NA glycoproteins at the same time (Fig. S25). Larger macro-clusters are more frequent and more stable in H1N1-Shan2009 than in H1N1-Mich2015. This is again primarily due to the missing N386 glycan in H1N1-Mich2015 NAs, which leads to disruption of the continuous pattern of connections between glycoproteins. The dynamic evolution of the glycoprotein macro-clusters is shown in Movie S10 for H1N1-Shan2009 and H1N1-Mich2015.

Subsequently, we sought to dissect the interaction modes between glycoproteins, i.e., how inter-connections are formed. There are three ways a glycoprotein can interact with another one: using i) protein residues only; ii) glycan residues only; iii) protein and glycan residues at the same time. The schematic and the analysis are shown in Fig. 6B. N-glycans play an important role in modulating the HA/NA interplay as they are largely involved in the glycoprotein interactions. The number of connections formed by HA involving glycans progressively increases during the simulations (Fig. 6B). At the end of the H1N1-Shan2009 simulation, 54.0% of connections made by HAs entail glycans, either alone (9.1%) or concomitant with protein residues (44.1%). Intriguingly, the introduction of glycan N179 within the HA head in H1N1-Mich2015 augments the participation of glycans in connections formed by HA to 66.7% (glycan only 11.2%, concomitant with protein 55.5%) (Fig. 6B). A similar trend is also observed for NA in H1N1-Shan2009. Instead, the opposite scenario takes place in the case of NA in H1N1-Mich2015, owing to the loss of glycan N386 within the head. While at the end of the H1N1-Shan2009 simulation, glycans are involved in 63.4% of connections formed by NA (16.7% alone, 46.7% with protein), in H1N1-Mich2015, only 13.7% of NA connections include glycans (3.4% alone, 10.3% with protein) (Fig. 6B). This finding highlights the critical role of NA glycan N386 in H1N1-Shan2009 as a molecular sensor to boost the glycoprotein-glycoprotein interplay. The relative frequency of connections involving glycans alone progressively decreases during the simulations in favor of interactions involving glycans and protein simultaneously and occurring only after the glycoproteins are brought in proximity to each other (Fig. 6B). Finally, an in-depth analysis of which glycans are mostly used by HA and NA to form connections reveals that NA preferably uses head glycans while HA mostly utilizes stalk glycans (Fig. S26). The relative frequency of interacting NA head glycans considerably drops in H1N1-Mich205 due to the loss of N386. Instead, the relative frequency of interacting HA head glycans considerably increases in H1N1-Mich205 due to the addition of N179 glycan (Fig. S26).

## Discussion

A critical determinant of influenza A infection, replication, and transmissibility is the functional balance between HA and NA (*8, 74, 84–86*). One fascinating aspect of the influenza viral cycle is that the two glycoproteins compete, with opposite actions, for the same substrate, i.e., the terminal sialic acid (SA) residue presented by host cell-linked glycans. While HA mediates viral entry by binding to SA, NA modulates the release of the viral progeny by cleaving SA moieties from both the sialylated receptors and HA (*74*). In a scenario where the virus fitness depends on the delicate, antigenic drift-dependent equilibrium between HA binding avidity and NA enzymatic activity (*8*), substantial knowledge gaps persist in the understanding of the HA/NA dynamic interplay taking place at the level of the virus surface. To shed light on this crucial aspect and its antigenic implications, we performed massive, all-atom whole-virion MD simulations of two evolutionarily linked influenza A strains, H1N1-Shan2009 (Fig. 1) and H1N1-Mich2015 (Fig. S1). Maintaining the accuracy of the atomic detail, we implemented a multiscale computational protocol that exploits different modeling tools, analysis methods, and algorithms for crossing spatial scales (from molecular to subcellular) and exploring events occurring in different temporal scales. Our simulations provided previously unseen dynamic views of the plastic conformational landscape of HA and NA ectodomains, revealing three main extensive motions: NA head tilt (Fig. 2), HA ectodomain tilt (Fig. 3), and HA head breathing (Fig. 4). While the last two motions find evidence in previous experimental work (*37, 43, 72*), to the best of our knowledge, no work has yet detailed the flexibility of the NA head relative to the stalk. In a recent study by Ellis et al. (*44*), the authors investigated the plasticity of a recombinant NA tetramer, revealing an open tetrameric assembly where the four heads move away from each other, giving the NA a clover-like appearance. As shown in Fig. S27, in our simulations, we did not observe the breathing of the monomeric head domains, most likely due to insufficient sampling and/or inconsistencies between our model and the experimental construct. Instead, the NA tetrameric head tilts as a single body as much as >90° relative to the stalk (Fig. 2C and Fig. S11). Although not discussed in Ellis et al. (*44*), cryoEM micrographs included therein hint at tilted head conformations potentially compatible with poses extracted from our simulations, thus indirectly supporting our findings. The tilting of prefusion HA trimer has been previously mentioned in a theoretical study (*87*) and more recently demonstrated by cryoEM experiments probing it in detergent micelles (*37*). Additionally, the predisposition to assume tilted orientations is a common property shared by class I fusion viral glycoproteins such as HIV Env (*88*) and SARS-CoV-2 spike (*53, 89*). The HA head breathing was initially supported by smFRET experiments (*90*) and then indirectly reported concomitant with the discovery of a cryptic epitope located at the interface between monomers in the head (*38, 39, 41, 43, 91*). Breathing has been reported for other viral glycoproteins in flavivirus (*92*) and HIV (*93*). Not only are our simulations in agreement with the experiments (*43*), but they are the first of their kind to show the HA breathing or flexing on the viral membrane in the context of the whole virion. The probability densities of NA head-tilt angle, HA ectodomain-tilt angle, and HA head–LαHs distance (Figs. 2C, 3C, 4C, respectively) indicate that open HA head (HA head–LαHs distance > 30 Å) or tilted HA (> 50°) and NA (>75°) conformations are rarely sampled during the simulations, whereas closed, untilted, or slightly tilted conformations are most likely. We also characterized the kinetics of all three motions, building a two-state MSM for each. MFPTs of HA head-breathing and HA ectodomain-tilt motions (Figs. 3D, 4D) indicate that transitioning to an open HA trimer or a tilted HA ectodomain is slower than recovering a closed or an upright conformation. Instead, MFPTs of the NA head-tilt motion pinpoint that the transition to a tilted state is faster than the reverse one.

The high-dimensional mobility of the HA and NA ectodomains proffers wide-ranging implications. From an immunogenic point of view, the glycoproteins’ accentuated conformational plasticity observed in our simulations has unveiled their vulnerabilities, exposing epitopes that otherwise would not be accessible or would be sterically occluded. As discussed above, tilted, open or partially open conformations are transient, and, most importantly, reversible. Stabilizing these fleeting states can be used as a strategy to lock the glycoproteins in vulnerable orientations that can be harnessed for vaccine development and the design of antiviral drugs. This could be achieved through amino acid substitutions imparting a specific tertiary/quaternary structure to the glycoprotein (*94*). The list of immunogenically relevant states revealed by our simulations includes a previously uncharacterized NA with a tilted head. We identified a novel epitope located on the underside of an N2-subtype NA (Fig. 2E, F and Figs. S15, S16) head that becomes directly accessible for immune recognition during head tilting. Importantly, the underside epitope is conserved in the NA of contemporary H3N2 viruses unlike the more accessible epitopes surrounding the catalytic site where the virus acquires an additional glycosylation site (*95*) resulting in resistance to a subset of antibodies targeting the catalytic site. This reaffirms the importance of NA, and potentially of its head underside, as a key target for vaccine and drug discovery.

Next, the flexibility of the HA ectodomain affects the approachability of a conserved epitope located at the juxta-membrane region of the HA stalk (Fig. 3E-F), also referred to as the anchor epitope. We showed that as the HA bends towards the membrane, the angle of approach to at least one of the anchor epitopes drastically changes, adjusting to a favorable orientation for antibody binding. Our simulations complement previous structural experiments that reported several broadly neutralizing antibodies targeting the anchor epitope (*37, 96*), confirming that the epitope is occluded by the underneath membrane when the HA is untilted (Fig. S19). Trapping the HA ectodomain in a tilted state could bolster the immune response towards the anchor epitope. Finally, our simulation unveiled how the cryptic epitope located at the protomer interface between the HA heads (*38, 39, 91*) becomes transiently exposed when HA breathes. Most of the HA heads remain in the closed conformation during the dynamics, and rarely open. When breathing occurs, the monomers fleetingly open at different times and rapidly rotate back to a closed position. We could not create an MSM with all three monomers breathing symmetrically, meaning that all three monomers breathing symmetrically did not occur often enough to be appropriately sampled, and thus if it occurs at all, will be at a lower probability than a single monomer breathing. Nonetheless, we demonstrated that reversible breathing of at least one monomer is enough to accommodate a broadly protective antibody, such as FluA-20 (*39*), at the interface epitope.

From a functional point of view, the conformational plasticity of influenza glycoproteins’ ectodomains would mediate the virus’ ability to traverse the cell surface (*9, 97, 98*). This plasticity could also facilitate gaining access to SA moieties on the host cell surface, either during the viral entry or egress processes. Tilting and breathing may help HA engage the flexible sialylated receptors in a dynamic and crowded environment. Head tilting makes NA like a weedwhacker on a rotating axis, capable of trimming terminal SA moieties from the host cell with a 180° swing range. Improved substrate binding may also indirectly increase the catalytic efficiency of NA, which was shown to be enzymatically active only as a tetramer (*99*). In both cases, the plasticity of the ectodomains helps accommodate the hypermobility of the polysaccharide chains presenting the targeted SA residues. Consequently, the functional HA/NA balance also depends on the HA/NA dynamic interplay (*8, 74*), i.e., the interactive set of interactions, conformational changes, and collective motions by which the glycoproteins affect each other’s dynamics and orientation on the virion surface. Structural factors such as NA stalk length, HA:NA stoichiometry, or the spatial distribution of the glycoproteins within the viral membrane, also contribute to fine-tuning the HA/NA balance (*84*). Yet, influenza virus is not a static system. As portrayed in the results section, the HA/NA interplay is indeed dynamic because the glycoproteins fleetingly tilt, bend, “breathe”, and move in such a way that the pattern of their interactions continuously changes over time (Fig. 6 and Figs. S24, S25, Movie S9 and Movie S10). Glycoproteins that were not contacting any other glycoproteins at the beginning of the simulations, ended up interacting with one, two, or more glycoproteins, forming clusters (Fig. 7A) and larger macro-clusters (Fig. 7B). Our simulations provided unprecedented views on the HA/NA interplay, where the SA binding sites on HA/NA move in proximity to each other upon concomitant tilting or breathing motions (Fig. 7A and Fig. 7B). Hence, NA and HA can compete for the same substrate (Fig. 7C), as suggested by a recent single-molecule force spectroscopy study (*26*). Owing to its head flexibility, NA can reach multiple HA binding sites at the same time and contend for the same sialylated cell receptors (Fig. 7C). When influenza virus is in the proximity of the host cell sialoglycan receptors, HA and NA establish multivalent interactions, i.e., multiple, non-covalent weak interactions, with the terminal sialic acid residues, leading to a firm attachment (*30*). This process allows virus internalization but can also modulate progeny release (*30*). Although the number of multivalent HA/NA-cell receptor interactions needed for a firm attachment is still unknown (*29, 30*), knowledge of the large degree of HA/NA ectodomain flexibility provided by our simulations may improve future predictions in this area.

**Fig. 7.**
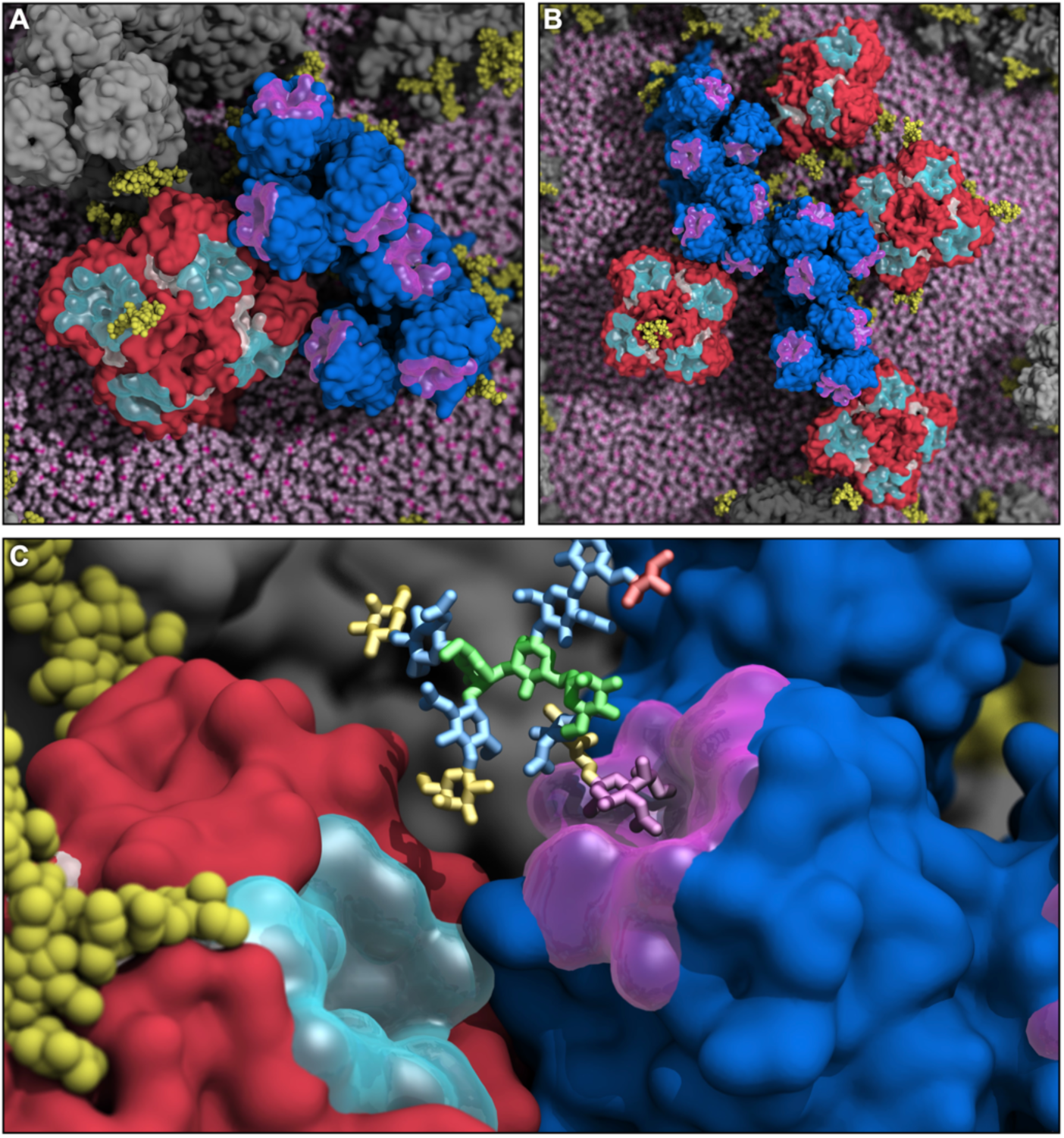
HA and NA interplay at the basis of the viral egress process. **A-B)** Molecular representation of snapshots retrieved from the simulation of H1N1-Shan2009 where HA and NA clump together, leading to the formation of clusters (**A**) and macro-clusters (**B**). The same representation scheme as panel A is adopted. The lipid envelope is depicted using pink vdW spheres. **C)** Close-up molecular representation of a snapshot from the simulation of H1N1-Shan2009 where HA binding site (semi-transparent magenta surface) and NA active site (semi-transparent cyan surface) are in proximity to each other, possibly competing for the same substrate. A complex glycan terminating with sialic acid, mimicking a putative host-cell receptor and represented with colored sticks, was docked into the HA binding site based on the pose of an HA substrate crystalized in PDB ID: 3UBE (*100*). The glycan residues are colored according to the Symbol Nomenclature for Glycans (SNFG) (*101*), where N-Acetylglucosamine is colored in blue, mannose in green, fucose in red, galactose in yellow, and sialic acid in purple. HA is shown with a blue surface, whereas NA with a red surface. N-glycans linked to HA and NA are highlighted with yellow vdW spheres.

Glycans are fundamental post-translational modifications of viral envelope proteins (*102, 103*), including HA and NA. As such, they exert a profound impact on enveloped virus biology and so on the overall fitness of the virus (*10, 104*). Among their many roles, glycans modulate evasion of the immune system by shielding antigenic epitopes from antibody recognition (*10, 105–109*), tune the interaction and affinity with the host cell (*110*), affect to a certain extent the binding of HA (*111*) and NA (*112*) to sialylated receptors, or can even contribute to protein folding (*113*) and HA cleavage (*114*). Here, we show that N-linked glycans seem to play a role in enhancing HA/NA interplay. Indeed, glycans form interactions with neighboring glycoproteins, acting as antennas. We note that the impact of glycans could be even more extensive than described in this work since the glycoproteins are not fully glycosylated in our virion models (see Material and Methods for further details). Despite the inter-glycoprotein interaction patterns being similar in H1N1-Shan2009 and H1N1-Mich2015 (Fig. 6A), substantial differences exist between the two strains in how the interactions are made (Fig. 6B, Fig. S26). Especially in the case of NA, the loss of glycan N386 in H1N1-Mich2015 within the head has reduced the ability of NA to form new connections, pinpointing possible impact on the structural/functional balance with HA during infection.

Overall, our massive whole-virion MD simulations provide previously unappreciated views on the influenza virus dynamics, highlighting the remarkable flexibility of the glycoproteins’ ectodomains. We characterized reversible conformational transitions, such as NA head tilt, HA ectodomain tilt, and HA head breathing, which also revealed potential vulnerabilities of influenza glycoproteins to the immune system that are conserved across two evolutionarily related H1N1 strains, specifically H1N1-Shan2009 and H1N1-Mich2015. We identified a novel epitope located on the underside face of the NA head, further attesting the NA as a key antigenic target. Finally, our work substantially advances the knowledge of the HA/NA dynamic interplay in the context of viral entry and egress processes, ushering in unprecedented views on glycoprotein cooperativity. Altogether, while expanding knowledge on the molecular determinants of infection, we envision that our work will contribute to a viable strategy for the development of a universal or more effective vaccine against influenza virus.

## Materials and methods summary

### Computational methods

Initial coordinates for the all-atom whole-virion models of H1N1-Shan2009 and H1N1-Mich2015 were derived from the last frame of all-atom MD simulations of the unglycosylated H1N1-Shan2009 whole-virion model previously performed by us and thoroughly described in Durrant et al. (*45*) This model includes 236 HA trimers, 30 NA tetramers, and 11 M2 proton channels embedded in a semi-spherical lipid bilayer (*45*). The M1 matrix proteins, coating the inner leaflet of the lipid bilayer, and the ribonucleoproteins contained inside of the virion were not included in the model. Instead, the virion interior was filled in with water molecules. The shape of the virion and the distribution of the glycoproteins in the viral membrane were adopted from a point model of the virion exterior provided by collaborators (*45, 115*), which was in turn derived from a cryoET map of the influenza virus pleiomorphy (*83*). H1N1-Mich2015 was generated from H1N1-Shan2009 upon modeling the point mutations within HA (15 per monomer) and NA (14 per monomer) by means of NAMD’s PSFGEN (*116*) and the CHARMM36 amino acid topologies (*117*). From this initial point, the main challenge was the glycosylation of the HA trimers and NA tetramers. In H1N1-Shan2009, HA and NA present six and eight N-linked sequons on each monomer, respectively. H1N1-Mich2015 introduced an additional glycosylation site within the HA head at position N179, whereas glycan N386 within the NA head was deleted. Building upon the coordinates derived from Durrant et al. (*118*), glycosylation of HA and NA was carried out with the doGlycans tool (*119*). The glycoprofile was based on available glycomics data (*11, 120–124*). Since glycans were added to an unglycosylated construct resulting from 69.74 ns of MD (*45*), many sequons ended up not being glycosylated either because sterically hindered by adjacent residues, occluded by neighboring glycoproteins, or not fully exposed to the solvent. The resulting, glycosylated H1N1-Shan2009 and H1N1-Mich2015 whole virion constructs, comprising 236 glycosylated HA homo-trimers, 30 glycosylated NA homo-tetramers, and 11 M2 homo-tetramers embedded in the POPC lipid bilayer, were parameterized using NAMD’s PSFGEN (*116*) and CHARMM36 all-atom additive force fields for protein (*117*), lipids (*125, 126*), glycans (*127*), and ions (*128*). As mentioned above, explicit TIP3 water molecules (*129*) located inside and outside of the lipid bilayer as well as Ca^2+^ ions bound to the NA heads, were retained from the original unglycosylated system (*45*). The number of Na^+^ and Cl^-^ ions was adjusted to neutralize the outstanding charge, with ionic strength set at 150 mM. The total number of atoms of the final systems is 160,919,594 for H1N1-Shan2009 and 160,981,954 for H1N1-Mich2015, with an orthorhombic periodic cell of ∼114 nm × ∼119 nm × ∼115 nm. All-atom MD simulations were performed on the ORNL TITAN supercomputer and the NSF Blue Waters supercomputer, respectively, using the CUDA memory-optimized version of NAMD 2.13 (*130*) and CHARMM36 force fields (*117, 125–127*). Upon minimization, heating, and equilibration, the systems were submitted to productive MD simulations in NPT conditions using the Langevin thermostat (*131*) and the Nosé-Hoover Langevin piston (*132, 133*) to achieve pressure (1.01325 bar) and temperature (298 K) control. One continuous replica was performed for each system using a timestep of 2 fs. Non-bonded interactions (van der Waals (vdW) and short-range electrostatic) were calculated at each timestep using a cutoff of 12 Å and a switching distance of 10 Å. All simulations were performed using periodic boundary conditions, employing the particle-mesh Ewald method (*134*) with a grid spacing of 2.1 Å to evaluate long-range electrostatic interactions every three timesteps. SHAKE algorithm (*135*) was adopted to keep the atomic bonds involving hydrogens fixed. Frames were saved every 30,000 steps (60 ps). For H1N1-Shan2009, we collected a total of ∼441.78 ns (7363 frames) continuous productive MD. For H1N1-Mich2015, we collected a total of 424.98 ns (7083 frames) continuous productive MD. All the analyses, including RMSD, RMSF, NA head tilt angle, HA ectodomain tilt angle, HA head breathing, AbASA, and glycoprotein interactions were all performed with Visual Molecular Dynamics (VMD) software (*136*) using in-house developed scripts. Figure panels and movies of the whole-virion simulations were rendered using VMD (*136*). MSMs were created with the PyEmma2 software program (*137*). We used one to two features, whose dimensionalities were then reduced and/or data reformatted through TICA (*138–140*). After discretization of the trajectories, we clustered them through k-means clustering (*141*). We then created Bayesian MSMs and validated them through Chapman-Kolmogorov tests (*142*). Next, we used Robust Perron Cluster Center Analysis (PCCA++) (*143*) to assign microstates to corresponding macrostates, and finally extracted 10 representative trajectory frames from the microstate with the highest probability assignment to its macrostate. Averaging the tilt angles from those 10 frames showed clear conformational transitions, allowing us to confidently use their associated MFPTs.

### Experimental methods

Monoclonal antibody (mAb) NDS.1 was isolated from an H3N2 influenza convalescent human donor by flow cytometry-based single B cell sorting with the N2 NA probe (A/Wisconsin/67/2005). Recombinant mAb NDS.1 was produced by transient transfection of expression vectors encoding the antibody’s heavy and light chains and purified by affinity chromatography using the protein A resin. Fab NDS.1 was generated by proteolytic digested using Lys-C endoproteinase. Binding kinetics of the Fab NDS.1 were measured by biolayer interferometry using the Octet HTX instrument. Negative-stain electron microscopy reconstruction model of N2 NA–Fab NDS.1 complex was generated with a final dataset of 2,304 particles.

## Acknowledgments

L.C. would like to thank Dr. Zied Gaieb for useful discussions on simulation analysis, and Dr. Jacob Durrant for the help he provided for system setup and simulation. C.S. would like to thank Dr. Emilia Pecora de Barros and Dr. Sarah Kochanek for useful discussions on MSM theory and use, and Moritz Hoffman for help with MSM feature selection. We thank Dr. James C. Phillips and Dr. John E. Stone for the incredible help and support with NAMD and VMD. We thank Dr. Meghan Altman and Dr. Pascal Gagneux for useful discussions on the glycan evolution pattern in the influenza virus.

## Author Contributions

L.C. constructed the influenza whole-virion models, performed MD simulations, designed and performed simulation analyses, prepared data for MSM analyses, and created figures and movies. C.S. designed and performed MSM analyses, created related plots, and helped design HA and NA glycoprofile for computational modeling. J.L. isolated and produced anti-NA antibody, performed BLI experiments, created respective figure panels, and wrote related results and methods. Y.T performed negative-stain EM experiments. M.K. designed and oversaw experiments. L.C., C.S., and R.E.A. wrote the original draft and all the authors contributed to reviewing and editing. I.A.W. mentored L.C. and provided critical feedback throughout. R.E.A. designed, oversaw, and secured resources for the research project.

## Data and materials availability

Jupyter-notebooks for the MSM analysis of NA head tilt, HA ectodomain tilt, and HA head breathing are available in the supplementary material (Data S1). The density file of the 3D map visualization of Fab NDS.1 in complex with A/Darwin/9/2021 at 24.1 Å resolution is made available as Data S2. Final snapshots from the whole-virion simulations will be made available on https://amarolab.ucsd.edu/. Full simulation data (30 TB), including trajectories and analysis scripts, will be made available upon request.

## Supplementary Materials

Materials and Methods

Figs. S1 to S27

Tables S1 to S3

Movies S1 to S10

Data S1 to S2

References (*144–173*)

## References and Notes

1. N. C. Wu, I. A. Wilson, Influenza Hemagglutinin Structures and Antibody Recognition. Cold Spring Harbor Perspectives in Medicine. 10, a038778 (2020).

2. S. Wilks, M. de Graaf, D. J. Smith, D. F. Burke, A review of influenza haemagglutinin receptor binding as it relates to pandemic properties. Vaccine. 30, 4369–4376 (2012).

3. J. J. Skehel, D. C. Wiley, Receptor Binding and Membrane Fusion in Virus Entry: The Influenza Hemagglutinin. Annual Review of Biochemistry. 69, 531–569 (2003).

4. G. M. Air, Influenza neuraminidase. Influenza and Other Respiratory Viruses. 6, 245 (2012).

5. R. M. Pielak, J. J. Chou, Influenza M2 proton channels. Biochimica et Biophysica Acta (BBA) - Biomembranes. 1808, 522–529 (2011).

6. L. H. Pinto, R. A. Lamb, The M2 Proton Channels of Influenza A and B Viruses *. Journal of Biological Chemistry. 281, 8997–9000 (2006).

7. R. Wagner, M. Matrosovich, H. D. Klenk, Functional balance between haemagglutinin and neuraminidase in influenza virus infections. Reviews in Medical Virology. 12, 159–166 (2002).

8. I. Kosik, J. W. Yewdell, Influenza Hemagglutinin and Neuraminidase: Yin–Yang Proteins Coevolving to Thwart Immunity. Viruses. 11, 346 (2019).

9. E. de Vries, W. Du, H. Guo, C. A. M. de Haan, Influenza A Virus Hemagglutinin–Neuraminidase–Receptor Balance: Preserving Virus Motility. Trends in Microbiology. 28, 57–67 (2020).

10. M. O. Altman, M. Angel, I. Košík, N. S. Trovão, S. J. Zost, J. S. Gibbs, L. Casalino, R. E. Amaro, S. E. Hensley, M. I. Nelson, J. W. Yewdell, Human influenza a virus hemagglutinin glycan evolution follows a temporal pattern to a glycan limit. mBio. 10, e00204–19 (2019).

11. S. Sun, Q. Wang, F. Zhao, W. Chen, Z. Li, Glycosylation Site Alteration in the Evolution of Influenza A (H1N1) Viruses. PLOS ONE. 6, e22844 (2011).

12. B. P. Blackburne, A. J. Hay, R. A. Goldstein, Changing Selective Pressure during Antigenic Changes in Human Influenza H3. PLOS Pathogens. 4, e1000058 (2008).

13. J. Gao, L. Couzens, D. F. Burke, H. Wan, P. Wilson, M. J. Memoli, X. Xu, R. Harvey, J. Wrammert, R. Ahmed, J. K. Taubenberger, D. J. Smith, R. A. M. Fouchier, M. C. Eichelberger, Antigenic drift of the influenza A(H1N1)pdm09 virus neuraminidase results in reduced effectiveness of A/California/7/2009 (H1N1pdm09)-specific antibodies. mBio. 10, 1–17 (2019).

14. F. Carrat, A. Flahault, Influenza vaccine: The challenge of antigenic drift. Vaccine. 25, 6852–6862 (2007).

15. M. F. Boni, Vaccination and antigenic drift in influenza. Vaccine. 26, C8–C14 (2008).

16. S. J. Zost, N. C. Wu, S. E. Hensley, I. A. Wilson, Immunodominance and Antigenic Variation of Influenza Virus Hemagglutinin: Implications for Design of Universal Vaccine Immunogens. The Journal of Infectious Diseases. 219, S38–S45 (2019).

17. B. L. Bullard, E. A. Weaver, Strategies Targeting Hemagglutinin as a Universal Influenza Vaccine. Vaccines (Basel). 9, 257 (2021).

18. S. Creytens, M. N. Pascha, M. Ballegeer, X. Saelens, C. A. M. de Haan, Influenza Neuraminidase Characteristics and Potential as a Vaccine Target. Frontiers in Immunology. 12, 786617 (2021).

19. R. Nachbagauer, J. Feser, A. Naficy, D. I. Bernstein, J. Guptill, E. B. Walter, F. Berlanda-Scorza, D. Stadlbauer, P. C. Wilson, T. Aydillo, M. A. Behzadi, D. Bhavsar, C. Bliss, C. Capuano, J. M. Carreño, V. Chromikova, C. Claeys, L. Coughlan, A. W. Freyn, C. Gast, A. Javier, K. Jiang, C. Mariottini, M. McMahon, M. McNeal, A. Solórzano, S. Strohmeier, W. Sun, M. van der Wielen, B. L. Innis, A. García-Sastre, P. Palese, F. Krammer, A chimeric hemagglutinin-based universal influenza virus vaccine approach induces broad and long-lasting immunity in a randomized, placebo-controlled phase I trial. Nature Medicine. 27, 106–114 (2020).

20. M. Lakadamyali, M. J. Rust, X. Zhuang, Endocytosis of influenza viruses. Microbes and Infection. 6, 929–936 (2004).

21. J. S. Rossman, R. A. Lamb, Influenza virus assembly and budding. Virology. 411, 229–236 (2011).

22. D. P. Nayak, R. A. Balogun, H. Yamada, Z. H. Zhou, S. Barman, Influenza virus morphogenesis and budding. Virus Research. 143, 147–161 (2009).

23. A. P. Schmitt, R. A. Lamb, Influenza Virus Assembly and Budding at the Viral Budozone. Advances in Virus Research. 64, 383–416 (2005).

24. C. Liu, † Maryna, C. Eichelberger, R. W. Compans, G. M. Air, Influenza type A virus neuraminidase does not play a role in viral entry, replication, assembly, or budding. Journal of Virology. 69, 1099–1106 (1995).

25. D. P. Nayak, E. K. W. Hui, S. Barman, Assembly and budding of influenza virus. Virus Research. 106, 147–165 (2004).

26. V. Reiter-Scherer, J. L. Cuellar-Camacho, S. Bhatia, R. Haag, A. Herrmann, D. Lauster, J. P. Rabe, Force Spectroscopy Shows Dynamic Binding of Influenza Hemagglutinin and Neuraminidase to Sialic Acid. Biophysical Journal. 116, 1037–1048 (2019).

27. R. Du, Q. Cui, L. Rong, Competitive Cooperation of Hemagglutinin and Neuraminidase during Influenza A Virus Entry. Viruses. 11, 458 (2019).

28. J. Yang, S. Liu, L. Du, S. Jiang, A new role of neuraminidase (NA) in the influenza virus life cycle: implication for developing NA inhibitors with novel mechanism of action. Reviews in Medical Virology. 26, 242–250 (2016).

29. N. J. Overeem, E. van der Vries, J. Huskens, N. J. Overeem, J. Huskens, E. van der Vries, A Dynamic, Supramolecular View on the Multivalent Interaction between Influenza Virus and Host Cell. Small. 17, 2007214 (2021).

30. M. Müller, D. Lauster, H. H. K. Wildenauer, A. Herrmann, S. Block, Mobility-Based Quantification of Multivalent Virus-Receptor Interactions: New Insights into Influenza A Virus Binding Mode. Nano Letters. 19, 1875–1882 (2019).

31. J. L. Cuellar-Camacho, S. Bhatia, V. Reiter-Scherer, D. Lauster, S. Liese, J. P. Rabe, A. Herrmann, R. Haag, Quantification of Multivalent Interactions between Sialic Acid and Influenza A Virus Spike Proteins by Single-Molecule Force Spectroscopy. J Am Chem Soc. 142, 12181–12192 (2020).

32. C. Sieben, C. Kappel, R. Zhu, A. Wozniak, C. Rankl, P. Hinterdorfer, H. Grubmüller, A. Herrmann, Influenza virus binds its host cell using multiple dynamic interactions. Proc Natl Acad Sci U S A. 109, 13626–13631 (2012).

33. A. Varki, Sialic acids as ligands in recognition phenomena. The FASEB Journal. 11, 248–255 (1997).

34. N. M. Varki, A. Varki, Diversity in cell surface sialic acid presentations: implications for biology and disease. Laboratory Investigation 2007 87:9. 87, 851–857 (2007).

35. D. C. Wiley, J. J. Skehel, The structure and function of the hemagglutinin membrane glycoprotein of influenza virus. Annual Review of Biochemistry. 56, 365–394 (2003).

36. Y. Suzuki, T. Ito, T. Suzuki, Jr. Robert E. Holland, T. M. Chambers, M. Kiso, H. Ishida, Y. Kawaoka, Sialic Acid Species as a Determinant of the Host Range of Influenza A Viruses. Journal of Virology. 74, 11825 (2000).

37. D. J. Benton, A. Nans, L. J. Calder, J. Turner, U. Neu, Y. P. Lin, E. Ketelaars, N. L. Kallewaard, D. Corti, A. Lanzavecchia, S. J. Gamblin, P. B. Rosenthal, J. J. Skehel, Influenza hemagglutinin membrane anchor. Proc Natl Acad Sci U S A. 115, 10112–10117 (2018).

38. A. Watanabe, K. R. McCarthy, M. Kuraoka, A. G. Schmidt, Y. Adachi, T. Onodera, K. Tonouchi, T. M. Caradonna, G. Bajic, S. Song, C. E. McGee, G. D. Sempowski, F. Feng, P. Urick, T. B. Kepler, Y. Takahashi, S. C. Harrison, G. Kelsoe, Antibodies to a Conserved Influenza Head Interface Epitope Protect by an IgG Subtype-Dependent Mechanism. Cell. 177, 1124–1135.e16 (2019).

39. S. Bangaru, S. Lang, M. Schotsaert, H. A. Vanderven, X. Zhu, N. Kose, R. Bombardi, J. A. Finn, S. J. Kent, P. Gilchuk, I. Gilchuk, H. L. Turner, A. García-Sastre, S. Li, A. B. Ward, I. A. Wilson, J. E. Crowe, A Site of Vulnerability on the Influenza Virus Hemagglutinin Head Domain Trimer Interface. Cell. 177, 1136–1152.e18 (2019).

40. G. Bajic, M. J. Maron, Y. Adachi, T. Onodera, K. R. McCarthy, C. E. McGee, G. D. Sempowski, Y. Takahashi, G. Kelsoe, M. Kuraoka, A. G. Schmidt, Influenza Antigen Engineering Focuses Immune Responses to a Subdominant but Broadly Protective Viral Epitope. Cell Host & Microbe. 25, 827–835.e6 (2019).

41. S. J. Zost, J. Dong, I. M. Gilchuk, P. Gilchuk, N. J. Thornburg, S. Bangaru, N. Kose, J. A. Finn, R. Bombardi, C. Soto, E. C. Chen, R. S. Nargi, R. E. Sutton, R. P. Irving, N. Suryadevara, J. B. Westover, R. H. Carnahan, H. L. Turner, S. Li, A. B. Ward, J. E. Crowe, Canonical features of human antibodies recognizing the influenza hemagglutinin trimer interface. The Journal of Clinical Investigation. 131, e146791 (2021).

42. K. R. McCarthy, J. Lee, A. Watanabe, M. Kuraoka, L. R. Robinson-Mccarthy, G. Georgiou, G. Kelsoe, S. C. Harrison, A prevalent focused human antibody response to the influenza virus hemagglutinin head interface. mBio. 12, e01144–21 (2021).

43. X. Zhu, J. Han, W. Sun, E. Puente-Massaguer, W. Yu, P. Palese, F. Krammer, A. B. Ward, I. A. Wilson, Influenza chimeric hemagglutinin structures in complex with broadly protective antibodies to the stem and trimer interface. Proc Natl Acad Sci U S A. 119, 1–8 (2022).

44. D. Ellis, J. Lederhofer, O. J. Acton, Y. Tsybovsky, S. Kephart, C. Yap, R. A. Gillespie, A. Creanga, A. Olshefsky, T. Stephens, D. Pettie, M. Murphy, C. Sydeman, M. Ahlrichs, S. Chan, A. J. Borst, Y.-J. Park, K. K. Lee, B. S. Graham, D. Veesler, N. P. King, M. Kanekiyo, Structure-based design of stabilized recombinant influenza neuraminidase tetramers. Nature Communications. 13, 1–16 (2022).

45. J. D. Durrant, S. E. Kochanek, L. Casalino, P. U. Ieong, A. C. Dommer, R. E. Amaro, Mesoscale All-Atom Influenza Virus Simulations Suggest New Substrate Binding Mechanism. ACS Central Science. 6, 189–196 (2020).

46. E. Tarasova, D. Nerukh, All-Atom Molecular Dynamics Simulations of Whole Viruses. Journal of Physical Chemistry Letters. 9, 5805–5809 (2018).

47. R. G. Huber, J. K. Marzinek, D. A. Holdbrook, P. J. Bond, Multiscale molecular dynamics simulation approaches to the structure and dynamics of viruses. Progress in Biophysics and Molecular Biology. 128, 121–132 (2017).

48. M. Chavent, A. L. Duncan, M. S. P. Sansom, Molecular dynamics simulations of membrane proteins and their interactions: from nanoscale to mesoscale. Current Opinion in Structural Biology. 40, 8–16 (2016).

49. S. J. Marrink, V. Corradi, P. C. T. Souza, H. I. Ingólfsson, D. P. Tieleman, M. S. P. Sansom, Computational Modeling of Realistic Cell Membranes. Chemical Reviews. 119, 6184–6226 (2019).

50. T. Reddy, D. Shorthouse, D. L. Parton, E. Jefferys, P. W. Fowler, M. Chavent, M. Baaden, M. S. P. Sansom, Nothing to Sneeze At: A Dynamic and Integrative Computational Model of an Influenza A Virion. Structure. 23, 584–597 (2015).

51. A. Yu, A. J. Pak, P. He, V. Monje-Galvan, L. Casalino, Z. Gaieb, A. C. Dommer, R. E. Amaro, G. A. Voth, A multiscale coarse-grained model of the SARS-CoV-2 virion. Biophysical Journal. 120, 1097–1104 (2021).

52. T. Reddy, M. S. P. Sansom, The Role of the Membrane in the Structure and Biophysical Robustness of the Dengue Virion Envelope. Structure. 24, 375–382 (2016).

53. L. Casalino, A. C. Dommer, Z. Gaieb, E. P. Barros, T. Sztain, S. H. Ahn, A. Trifan, A. Brace, A. T. Bogetti, A. Clyde, H. Ma, H. Lee, M. Turilli, S. Khalid, L. T. Chong, C. Simmerling, D. J. Hardy, J. D. C. Maia, J. C. Phillips, T. Kurth, A. C. Stern, L. Huang, J. D. McCalpin, M. Tatineni, T. Gibbs, J. E. Stone, S. Jha, A. Ramanathan, R. E. Amaro, AI-driven multiscale simulations illuminate mechanisms of SARS-CoV-2 spike dynamics. The International Journal of High Performance Computing Applications. 35, 432–451 (2021).

54. J. R. Perilla, K. Schulten, Physical properties of the HIV-1 capsid from all-atom molecular dynamics simulations. Nature Communications. 8, 1–10 (2017).

55. P. L. Freddolino, A. S. Arkhipov, S. B. Larson, A. McPherson, K. Schulten, Molecular Dynamics Simulations of the Complete Satellite Tobacco Mosaic Virus. Structure. 14, 437–449 (2006).

56. E. Tarasova, V. Farafonov, R. Khayat, N. Okimoto, T. S. Komatsu, M. Taiji, D. Nerukh, All-Atom Molecular Dynamics Simulations of Entire Virus Capsid Reveal the Role of Ion Distribution in Capsid’s Stability. Journal of Physical Chemistry Letters. 8, 779–784 (2017).

57. Y. Andoh, N. Yoshii, A. Yamada, K. Fujimoto, H. Kojima, K. Mizutani, A. Nakagawa, A. Nomoto, S. Okazaki, All-atom molecular dynamics calculation study of entire poliovirus empty capsids in solution. The Journal of Chemical Physics. 141, 165101 (2014).

58. E. Tarasova, I. Korotkin, V. Farafonov, S. Karabasov, D. Nerukh, Complete virus capsid at all-atom resolution: Simulations using molecular dynamics and hybrid molecular dynamics/hydrodynamics methods reveal semipermeable membrane function. Journal of Molecular Liquids. 245, 109–114 (2017).

59. Y. Miao, J. E. Johnson, P. J. Ortoleva, All-atom multiscale simulation of cowpea chlorotic mottle virus capsid swelling. Journal of Physical Chemistry B. 114, 11181–11195 (2010).

60. J. A. Hadden, J. R. Perilla, C. J. Schlicksup, B. Venkatakrishnan, A. Zlotnick, K. Schulten, All-atom molecular dynamics of the HBV capsid reveals insights into biological function and cryo-EM resolution limits. Elife. 7, e32478 (2018).

61. V. S. Farafonov, D. Nerukh, MS2 bacteriophage capsid studied using all-atom molecular dynamics. Interface Focus. 9 (2019), doi:10.1098/RSFS.2018.0081.

62. A. Dommer, L. Casalino, F. Kearns, M. Rosenfeld, N. Wauer, S.-H. Ahn, J. Russo, S. Oliveira, C. Morris, A. Bogetti, A. Trifan, A. Brace, T. Sztain, A. Clyde, H. Ma, C. Chennubhotla, H. Lee, M. Turilli, S. Khalid, T. Tamayo-Mendoza, M. Welborn, A. Christensen, D. G. A Smith, Z. Qiao, S. Krishna Sirumalla, F. Manby, A. Anandkumar, D. Hardy, J. Phillips, A. Stern, J. Romero, D. Clark, M. Dorrell, T. Maiden, L. Huang, J. McCalpin, C. Woods, A. Gray, M. Williams, B. Barker, H. Rajapaksha, R. Pitts, T. Gibbs, J. Stone, D. Zuckerman, A. Mulholland, T. Miller III, S. Jha, A. Ramanathan, L. Chong, R. Amaro, C. Woods, bioRxiv, in press, doi:10.1101/2021.11.12.468428.

63. B. E. Husic, V. S. Pande, Markov State Models: From an Art to a Science. J Am Chem Soc. 140, 2386–2396 (2018).

64. V. S. Pande, K. Beauchamp, G. R. Bowman, Everything you wanted to know about Markov State Models but were afraid to ask. Methods. 52, 99–105 (2010).

65. J. D. Chodera, F. Noé, Markov state models of biomolecular conformational dynamics. Current Opinion in Structural Biology. 25, 135–144 (2014).

66. J. L. McAuley, B. P. Gilbertson, S. Trifkovic, L. E. Brown, J. L. McKimm-Breschkin, Influenza Virus Neuraminidase Structure and Functions. Frontiers in Microbiology. 10, 39 (2019).

67. D. Stadlbauer, X. Zhu, M. McMahon, J. S. Turner, T. J. Wohlbold, A. J. Schmitz, S. Strohmeier, W. Yu, R. Nachbagauer, P. A. Mudd, I. A. Wilson, A. H. Ellebedy, F. Krammer, Broadly protective human antibodies that target the active site of influenza virus neuraminidase. Science (1979). 366, 499–504 (2019).

68. J. J. Guthmiller, J. Han, H. A. Utset, L. Li, L. Y. L. Lan, C. Henry, C. T. Stamper, M. McMahon, G. O’Dell, M. L. Fernández-Quintero, A. W. Freyn, F. Amanat, O. Stovicek, L. Gentles, S. T. Richey, A. T. de la Peña, V. Rosado, H. L. Dugan, N. Y. Zheng, M. E. Tepora, D. J. Bitar, S. Changrob, S. Strohmeier, M. Huang, A. García-Sastre, K. R. Liedl, J. D. Bloom, R. Nachbagauer, P. Palese, F. Krammer, L. Coughlan, A. B. Ward, P. C. Wilson, Broadly neutralizing antibodies target a haemagglutinin anchor epitope. Nature. 602, 314–320 (2021).

69. R. Xu, D. C. Ekiert, J. C. Krause, R. Hai, J. E. Crowe, I. A. Wilson, Structural basis of preexisting immunity to the 2009 H1N1 pandemic influenza virus. Science (1979). 328, 357–360 (2010).

70. C. J. Russell, J. A. Belser, J. A. Pulit-Penaloza, X. Sun, Hemagglutinin Stability and Its Impact on Influenza A Virus Infectivity, Pathogenicity, and Transmissibility in Avians, Mice, Swine, Seals, Ferrets, and Humans. Viruses. 13, 746 (2021).

71. R. Brian Dyer, M. W. Eller, Dynamics of hemagglutinin-mediated membrane fusion. Proc Natl Acad Sci U S A. 115, 8655–8657 (2018).

72. Y. Wu, G. F. Gao, “Breathing” Hemagglutinin Reveals Cryptic Epitopes for Universal Influenza Vaccine Design. Cell. 177, 1086–1088 (2019).

73. J. Lee, D. R. Boutz, V. Chromikova, M. G. Joyce, C. Vollmers, K. Leung, A. P. Horton, B. J. DeKosky, C. H. Lee, J. J. Lavinder, E. M. Murrin, C. Chrysostomou, K. H. Hoi, Y. Tsybovsky, P. v. Thomas, A. Druz, B. Zhang, Y. Zhang, L. Wang, W. P. Kong, D. Park, L. I. Popova, C. L. Dekker, M. M. Davis, C. E. Carter, T. M. Ross, A. D. Ellington, P. C. Wilson, E. M. Marcotte, J. R. Mascola, G. C. Ippolito, F. Krammer, S. R. Quake, P. D. Kwong, G. Georgiou, Molecular-level analysis of the serum antibody repertoire in young adults before and after seasonal influenza vaccination. Nature Medicine. 22, 1456–1464 (2016).

74. L. Byrd-Leotis, R. D. Cummings, D. A. Steinhauer, The Interplay between the Host Receptor and Influenza Virus Hemagglutinin and Neuraminidase. International Journal of Molecular Sciences. 18, 1541 (2017).

75. D. L. Parton, A. Tek, M. Baaden, M. S. P. Sansom, Formation of Raft-Like Assemblies within Clusters of Influenza Hemagglutinin Observed by MD Simulations. PLOS Computational Biology. 9, e1003034 (2013).

76. M. Takeda, G. P. Leser, C. J. Russell, R. A. Lamb, Influenza virus hemagglutinin concentrates in lipid raft microdomains for efficient viral fusion. Proc Natl Acad Sci U S A. 100, 14610–14617 (2003).

77. G. P. Leser, R. A. Lamb, Lateral organization of influenza virus proteins in the budozone region of the plasma membrane. Journal of Virology. 91, e02104–16 (2017).

78. R. D. Cohen, G. J. Pielak, Electrostatic Contributions to Protein Quinary Structure. J Am Chem Soc. 138, 13139–13142 (2016).

79. X. Mu, S. Choi, L. Lang, D. Mowray, N. v. Dokholyan, J. Danielsson, M. Oliveberg, Physicochemical code for quinary protein interactions in Escherichia coli. Proc Natl Acad Sci U S A. 114, E4556–E4563 (2017).

80. G. P. Leser, R. A. Lamb, Influenza virus assembly and budding in raft-derived microdomains: A quantitative analysis of the surface distribution of HA, NA and M2 proteins. Virology. 342, 215–227 (2005).

81. M. Veit, B. Thaa, Association of influenza virus proteins with membrane rafts. Advances in Virology. 2011, 370606 (2011).

82. M. M. Rickard, Y. Zhang, M. Gruebele, T. v. Pogorelov, In-Cell Protein-Protein Contacts: Transient Interactions in the Crowd. Journal of Physical Chemistry Letters. 10, 5667–5673 (2019).

83. A. Harris, G. Cardone, D. C. Winkler, J. B. Heymann, M. Brecher, J. M. White, A. C. Steven, Influenza virus pleiomorphy characterized by cryoelectron tomography. Proc Natl Acad Sci U S A. 103, 19123–19127 (2006).

84. R. Du, Q. Cui, L. Rong, Competitive Cooperation of Hemagglutinin and Neuraminidase during Influenza A Virus Entry. Viruses. 11, 458 (2019).

85. S. Diederich, Y. Berhane, C. Embury-Hyatt, T. Hisanaga, K. Handel, C. Cottam-Birt, C. Ranadheera, D. Kobasa, J. Pasick, Hemagglutinin-Neuraminidase Balance Influences the Virulence Phenotype of a Recombinant H5N3 Influenza A Virus Possessing a Polybasic HA 0 Cleavage Site. Journal of Virology. 89, 10724–10734 (2015).

86. H. L. Yen, C. H. Liang, C. Y. Wu, H. L. Forrest, A. Ferguson, K. T. Choy, J. Jones, D. D. Y. Wong, P. P. H. Cheung, C. H. Hsu, O. T. Li, K. M. Yuen, R. W. Y. Chan, L. L. M. Poon, M. C. W. Chan, J. M. Nicholls, S. Krauss, C. H. Wong, Y. Guan, R. G. Webster, R. J. Webby, M. Peiris, Hemagglutinin-neuraminidase balance confers respiratory-droplet transmissibility of the pandemic H1N1 influenza virus in ferrets. Proc Natl Acad Sci U S A. 108, 14264–14269 (2011).

87. B. Isin, P. Doruker, I. Bahar, Functional Motions of Influenza Virus Hemagglutinin: A Structure-Based Analytical Approach. Biophysical Journal. 82, 569–581 (2002).

88. K. Rantalainen, Z. T. Berndsen, A. Antanasijevic, T. Schiffner, X. Zhang, W. H. Lee, J. L. Torres, L. Zhang, A. Irimia, J. Copps, K. H. Zhou, Y. D. Kwon, W. H. Law, C. A. Schramm, R. Verardi, S. J. Krebs, P. D. Kwong, N. A. Doria-Rose, I. A. Wilson, M. B. Zwick, J. R. Yates, W. R. Schief, A. B. Ward, HIV-1 Envelope and MPER Antibody Structures in Lipid Assemblies. Cell Reports. 31, 107583 (2020).

89. B. Turoňová, M. Sikora, C. Schürmann, W. J. H. Hagen, S. Welsch, F. E. C. Blanc, S. von Bülow, M. Gecht, K. Bagola, C. Hörner, G. van Zandbergen, J. Landry, N. T. D. de Azevedo, S. Mosalaganti, A. Schwarz, R. Covino, M. D. Mühlebach, G. Hummer, J. K. Locker, M. Beck, In situ structural analysis of SARS-CoV-2 spike reveals flexibility mediated by three hinges. Science. 370, 203–208 (2020).

90. D. K. Das, R. Govindan, I. Nikić-Spiegel, F. Krammer, E. A. Lemke, J. B. Munro, Direct Visualization of the Conformational Dynamics of Single Influenza Hemagglutinin Trimers. Cell. 174, 926–937.e12 (2018).

91. Y. Wu, G. F. Gao, “Breathing” Hemagglutinin Reveals Cryptic Epitopes for Universal Influenza Vaccine Design. Cell. 177, 1086–1088 (2019).

92. K. A. Dowd, T. C. Pierson, The many faces of a dynamic virion: implications of viral breathing on flavivirus biology and immunogenicity. Annual Review of Virology. 5, 185–207 (2018).

93. G. Ozorowski, J. Pallesen, N. de Val, D. Lyumkis, C. A. Cottrell, J. L. Torres, J. Copps, R. L. Stanfield, A. Cupo, P. Pugach, J. P. Moore, I. A. Wilson, A. B. Ward, Open and closed structures reveal allostery and pliability in the HIV-1 envelope spike. Nature. 547, 360–363 (2017).

94. P. O. Byrne, J. S. McLellan, Principles and practical applications of structure-based vaccine design. Current Opinion in Immunology. 77, 102209 (2022).

95. H. Wan, J. Gao, H. Yang, S. Yang, R. Harvey, Y. Q. Chen, N. Y. Zheng, J. Chang, P. J. Carney, X. Li, E. Plant, L. Jiang, L. Couzens, C. Wang, S. Strohmeier, W. W. Wu, R. F. Shen, F. Krammer, J. F. Cipollo, P. C. Wilson, J. Stevens, X. F. Wan, M. C. Eichelberger, Z. Ye, The neuraminidase of A(H3N2) influenza viruses circulating since 2016 is antigenically distinct from the A/Hong Kong/4801/2014 vaccine strain. Nature Microbiology 2019 4:12. 4, 2216–2225 (2019).

96. J. J. Guthmiller, J. Han, H. A. Utset, L. Li, L. Y. L. Lan, C. Henry, C. T. Stamper, M. McMahon, G. O’Dell, M. L. Fernández-Quintero, A. W. Freyn, F. Amanat, O. Stovicek, L. Gentles, S. T. Richey, A. T. de la Peña, V. Rosado, H. L. Dugan, N. Y. Zheng, M. E. Tepora, D. J. Bitar, S. Changrob, S. Strohmeier, M. Huang, A. García-Sastre, K. R. Liedl, J. D. Bloom, R. Nachbagauer, P. Palese, F. Krammer, L. Coughlan, A. B. Ward, P. C. Wilson, Broadly neutralizing antibodies target a haemagglutinin anchor epitope. Nature. 602, 314–320 (2021).

97. F. Ziebert, I. M. Kulić, How Influenza’s Spike Motor Works. Physical Review Letters. 126, 218101 (2021).

98. T. Sakai, S. I. Nishimura, T. Naito, M. Saito, Influenza A virus hemagglutinin and neuraminidase act as novel motile machinery. Scientific Reports 2017 7:1. 7, 1–11 (2017).

99. G. M. Air, Influenza neuraminidase. Influenza and Other Respiratory Viruses. 6, 245 (2012).

100. R. Xu, R. McBride, C. M. Nycholat, J. C. Paulson, I. A. Wilson, Structural Characterization of the Hemagglutinin Receptor Specificity from the 2009 H1N1 Influenza Pandemic. Journal of Virology. 86, 982–990 (2012).

101. A. Varki, R. D. Cummings, M. Aebi, N. H. Packer, P. H. Seeberger, J. D. Esko, P. Stanley, G. Hart, A. Darvill, T. Kinoshita, J. J. Prestegard, R. L. Schnaar, H. H. Freeze, J. D. Marth, C. R. Bertozzi, M. E. Etzler, M. Frank, J. F. G. Vliegenthart, T. Lütteke, S. Perez, E. Bolton, P. Rudd, J. Paulson, M. Kanehisa, P. Toukach, K. F. Aoki-Kinoshita, A. Dell, H. Narimatsu, W. York, N. Taniguchi, S. Kornfeld, Symbol Nomenclature for Graphical Representations of Glycans. Glycobiology. 25, 1323–1324 (2015).

102. Y. Watanabe, Z. T. Berndsen, J. Raghwani, G. E. Seabright, J. D. Allen, O. G. Pybus, J. S. McLellan, I. A. Wilson, T. A. Bowden, A. B. Ward, M. Crispin, Vulnerabilities in coronavirus glycan shields despite extensive glycosylation. Nature Communications. 11, 1–10 (2020).

103. Y. Watanabe, T. A. Bowden, I. A. Wilson, M. Crispin, Exploitation of glycosylation in enveloped virus pathobiology. Biochimica et Biophysica Acta (BBA) - General Subjects. 1863, 1480–1497 (2019).

104. I. T. Schulze, Effects of Glycosylation on the Properties and Functions of Influenza Virus Hemagglutinin. The Journal of Infectious Diseases. 176, S24–S28 (1997).

105. M. D. Tate, E. R. Job, Y. M. Deng, V. Gunalan, S. Maurer-Stroh, P. C. Reading, Playing Hide and Seek: How Glycosylation of the Influenza Virus Hemagglutinin Can Modulate the Immune Response to Infection. Viruses. 6, 1294–1316 (2014).

106. E. R. Job, Y.-M. Deng, K. K. Barfod, M. D. Tate, N. Caldwell, S. Reddiex, S. Maurer-Stroh, A. G. Brooks, P. C. Reading, Addition of Glycosylation to Influenza A Virus Hemagglutinin Modulates Antibody-Mediated Recognition of H1N1 2009 Pandemic Viruses. The Journal of Immunology. 190, 2169–2177 (2013).

107. X. Sun, A. Jayaraman, P. Maniprasad, R. Raman, K. v. Houser, C. Pappas, H. Zeng, R. Sasisekharan, J. M. Katz, T. M. Tumpey, N-Linked Glycosylation of the Hemagglutinin Protein Influences Virulence and Antigenicity of the 1918 Pandemic and Seasonal H1N1 Influenza A Viruses. Journal of Virology. 87, 8756–8766 (2013).

108. R. A. Medina, S. Stertz, B. Manicassamy, P. Zimmermann, X. Sun, R. A. Albrecht, H. Uusi-Kerttula, O. Zagordi, R. B. Belshe, S. E. Frey, T. M. Tumpey, A. García-Sastre, Glycosylations in the globular head of the hemagglutinin protein modulate the virulence and antigenic properties of the H1N1 influenza viruses. Science Translational Medicine. 5, 187ra70 (2013).

109. L. Casalino, Z. Gaieb, J. A. Goldsmith, C. K. Hjorth, A. C. Dommer, A. M. Harbison, C. A. Fogarty, E. P. Barros, B. C. Taylor, J. S. Mclellan, E. Fadda, R. E. Amaro, Beyond shielding: The roles of glycans in the SARS-CoV-2 spike protein. ACS Central Science. 6, 1722–1734 (2020).

110. N. L. Miller, T. Clark, R. Raman, R. Sasisekharan, Glycans in Virus-Host Interactions: A Structural Perspective. Frontiers in Molecular Biosciences. 8, 666756 (2021).

111. C. C. Wang, J. R. Chen, Y. C. Tseng, C. H. Hsu, Y. F. Hung, S. W. Chen, C. M. Chen, K. H. Khoo, T. J. Cheng, Y. S. E. Cheng, J. T. Jan, C. Y. Wu, C. Ma, C. H. Wong, Glycans on influenza hemagglutinin affect receptor binding and immune response. Proc Natl Acad Sci U S A. 106, 18137–18142 (2009).

112. C. Seitz, L. Casalino, R. Konecny, G. Huber, R. E. Amaro, J. A. McCammon, Multiscale Simulations Examining Glycan Shield Effects on Drug Binding to Influenza Neuraminidase. Biophysical Journal. 119, 2275–2289 (2020).

113. R. Daniels, B. Kurowski, A. E. Johnson, D. N. Hebert, N-Linked Glycans Direct the Cotranslational Folding Pathway of Influenza Hemagglutinin. Molecular Cell. 11, 79–90 (2003).

114. K. L. Deshpande, V. A. Fried, M. Ando, R. G. Webster, Glycosylation affects cleavage of an H5N2 influenza virus hemagglutinin and regulates virulence. Proc Natl Acad Sci U S 84, 36–40 (1987).

115. R. E. Amaro, P. U. Ieong, G. Huber, A. Dommer, A. C. Steven, R. M. Bush, J. D. Durrant, L. W. Votapka, A Computational Assay that Explores the Hemagglutinin/Neuraminidase Functional Balance Reveals the Neuraminidase Secondary Site as a Novel Anti-Influenza Target. ACS Central Science. 4, 1570–1577 (2018).

116. J. C. Phillips, R. Braun, W. Wang, J. Gumbart, E. Tajkhorshid, E. Villa, C. Chipot, R. D. Skeel, L. Kalé, K. Schulten, Scalable molecular dynamics with NAMD. J Comput Chem. 26, 1781–1802 (2005).

117. J. Huang, A. D. Mackerell, CHARMM36 all-atom additive protein force field: validation based on comparison to NMR data. J Comput Chem. 34, 2135–2145 (2013).

118. J. D. Durrant, R. E. Amaro, LipidWrapper: An Algorithm for Generating Large-Scale Membrane Models of Arbitrary Geometry. PLOS Computational Biology. 10, e1003720 (2014).

119. R. Danne, C. Poojari, H. Martinez-Seara, S. Rissanen, F. Lolicato, T. Róg, I. Vattulainen, DoGlycans-Tools for Preparing Carbohydrate Structures for Atomistic Simulations of Glycoproteins, Glycolipids, and Carbohydrate Polymers for GROMACS. Journal of Chemical Information and Modeling. 57, 2401–2406 (2017).

120. E. Cruz, J. Cain, B. Crossett, V. Kayser, Site-specific glycosylation profile of influenza A (H1N1) hemagglutinin through tandem mass spectrometry. Human Vaccines & Immunotherapeutics. 14, 508–517 (2017).

121. K. Khatri, J. A. Klein, M. R. White, O. C. Grant, N. Leymarie, R. J. Woods, K. L. Hartshorn, J. Zaia, Integrated omics and computational glycobiology reveal structural basis for influenza a virus glycan microheterogeneity and host interactions. Molecular and Cellular Proteomics. 15, 1895–1912 (2016).

122. Y. J. Liu, S. L. Wu, K. R. Love, W. S. Hancock, Characterization of Site-Specific Glycosylation in Influenza A Virus Hemagglutinin Produced by Spodoptera frugiperda Insect Cell Line. Analytical Chemistry. 89, 11036–11043 (2017).

123. Y. M. She, A. Farnsworth, X. Li, T. D. Cyr, Topological N-glycosylation and site-specific N-glycan sulfation of influenza proteins in the highly expressed H1N1 candidate vaccines. Scientific Reports. 7, 10232 (2017).

124. T. A. Blake, T. L. Williams, J. L. Pirkle, J. R. Barr, Targeted N-Linked Glycosylation Analysis of H5N1 Influenza Hemagglutinin by Selective Sample Preparation and Liquid Chromatography/Tandem Mass Spectrometry. Analytical Chemistry. 81, 3109–3118 (2009).

125. J. B. Klauda, R. M. Venable, J. A. Freites, J. W. O’Connor, D. J. Tobias, C. Mondragon-Ramirez, I. Vorobyov, A. D. MacKerell, R. W. Pastor, Update of the CHARMM All-Atom Additive Force Field for Lipids: Validation on Six Lipid Types. Journal of Physical Chemistry B. 114, 7830–7843 (2010).

126. J. B. Klauda, V. Monje, T. Kim, W. Im, Improving the CHARMM force field for polyunsaturated fatty acid chains. Journal of Physical Chemistry B. 116, 9424–9431 (2012).

127. O. Guvench, E. Hatcher, R. M. Venable, R. W. Pastor, A. D. MacKerell, CHARMM additive all-atom force field for glycosidic linkages between hexopyranoses. Journal of Chemical Theory and Computation. 5, 2353–2370 (2009).

128. D. Beglov, B. Roux, Finite representation of an infinite bulk system: Solvent boundary potential for computer simulations. The Journal of Chemical Physics. 100, 9050 (1998).

129. W. L. Jorgensen, J. Chandrasekhar, J. D. Madura, R. W. Impey, M. L. Klein, Comparison of simple potential functions for simulating liquid water. The Journal of Chemical Physics. 79, 926 (1998).

130. J. C. Phillips, D. J. Hardy, J. D. C. Maia, J. E. Stone, J. v. Ribeiro, R. C. Bernardi, R. Buch, G. Fiorin, J. Hénin, W. Jiang, R. McGreevy, M. C. R. Melo, B. K. Radak, R. D. Skeel, A. Singharoy, Y. Wang, B. Roux, A. Aksimentiev, Z. Luthey-Schulten, L. v. Kalé, K. Schulten, C. Chipot, E. Tajkhorshid, Scalable molecular dynamics on CPU and GPU architectures with NAMD. The Journal of Chemical Physics. 153, 044130 (2020).

131. A. Brünger, C. L. Brooks, M. Karplus, Stochastic boundary conditions for molecular dynamics simulations of ST2 water. Chemical Physics Letters. 105, 495–500 (1984).

132. G. J. Martyna, D. J. Tobias, M. L. Klein, Constant pressure molecular dynamics algorithms. The Journal of Chemical Physics. 101, 4177 (1998).

133. S. E. Feller, Y. Zhang, R. W. Pastor, B. R. Brooks, Constant pressure molecular dynamics simulation: The Langevin piston method. The Journal of Chemical Physics. 103, 4613 (1998).

134. T. Darden, D. York, L. Pedersen, Particle mesh Ewald: An N⋅log(N) method for Ewald sums in large systems. The Journal of Chemical Physics. 98, 10089 (1998).

135. J. P. Ryckaert, G. Ciccotti, H. J. C. Berendsen, Numerical integration of the cartesian equations of motion of a system with constraints: molecular dynamics of n-alkanes. Journal of Computational Physics. 23, 327–341 (1977).

136. W. Humphrey, A. Dalke, K. Schulten, VMD: visual molecular dynamics. J Mol Graph. 14, 33–38 (1996).

137. M. K. Scherer, B. Trendelkamp-Schroer, F. Paul, G. Pérez-Hernández, M. Hoffmann, N. Plattner, C. Wehmeyer, J. H. Prinz, F. Noé, PyEMMA 2: A Software Package for Estimation, Validation, and Analysis of Markov Models. Journal of Chemical Theory and Computation. 11, 5525–5542 (2015).

138. G. Pérez-Hernández, F. Paul, T. Giorgino, G. de Fabritiis, F. Noé, Identification of slow molecular order parameters for Markov model construction. The Journal of Chemical Physics. 139, 015102 (2013).

139. L. Molgedey, H. G. Schuster, Separation of a mixture of independent signals using time delayed correlations. Physical Review Letters. 72, 3634 (1994).

140. C. R. Schwantes, V. S. Pande, Improvements in Markov State Model construction reveal many non-native interactions in the folding of NTL9. Journal of Chemical Theory and Computation. 9, 2000–2009 (2013).

141. H.-H. Bock, Clustering Methods: A History of k-Means Algorithms, 161–172 (2007).

142. J. H. Prinz, H. Wu, M. Sarich, B. Keller, M. Senne, M. Held, J. D. Chodera, C. Schtte, F. Noé, Markov models of molecular kinetics: Generation and validation. The Journal of Chemical Physics. 134, 174105 (2011).

143. S. Röblitz, M. Weber, Fuzzy spectral clustering by PCCA+: Application to Markov state models and data classification. Advances in Data Analysis and Classification. 7, 147–179 (2013).

144. S. Jo, J. B. Lim, J. B. Klauda, W. Im, CHARMM-GUI Membrane Builder for Mixed Bilayers and Its Application to Yeast Membranes. Biophysical Journal. 97, 50 (2009).

145. S. Jo, T. Kim, W. Im, Automated Builder and Database of Protein/Membrane Complexes for Molecular Dynamics Simulations. PLOS ONE. 2, e880 (2007).

146. J. v. Vermaas, C. G. Mayne, E. Shinn, E. Tajkhorshid, Assembly and Analysis of Cell-Scale Membrane Envelopes. Journal of Chemical Information and Modeling. 62, 602–617 (2022).

147. P. T. Ivanova, D. S. Myers, S. B. Milne, J. L. McClaren, P. G. Thomas, H. A. Brown, Lipid composition of viral envelope of three strains of influenza virus – not all viruses are created equal. ACS Infect Dis. 1, 399–452 (2015).

148. J. D. Durrant, J. A. McCammon, AutoClickChem: Click Chemistry in Silico. PLOS Computational Biology. 8, e1002397 (2012).

149. P. Ropp, A. Friedman, J. D. Durrant, Scoria: A Python module for manipulating 3D molecular data. Journal of Cheminformatics. 9, 1–7 (2017).

150. M. Sharma, M. Yi, H. Dong, H. Qin, E. Peterson, D. D. Busath, H. X. Zhou, T. A. Cross, Insight into the mechanism of the influenza A proton channel from a structure in a lipid bilayer. Science (1979). 330, 509–512 (2010).

151. E. Kirkpatrick, X. Qiu, P. C. Wilson, J. Bahl, F. Krammer, The influenza virus hemagglutinin head evolves faster than the stalk domain. Scientific Reports 2018 8:1. 8, 1–14 (2018).

152. A. Pralow, M. Hoffmann, T. Nguyen-Khuong, M. Pioch, R. Hennig, Y. Genzel, E. Rapp, U. Reichl, Comprehensive N-glycosylation analysis of the influenza A virus proteins HA and NA from adherent and suspension MDCK cells. The FEBS Journal. 288, 4869–4891 (2021).

153. M. J. Abraham, T. Murtola, R. Schulz, S. Páll, J. C. Smith, B. Hess, E. Lindah, GROMACS: High performance molecular simulations through multi-level parallelism from laptops to supercomputers. SoftwareX. 1–2, 19–25 (2015).

154. J. Li, S. Liu, Y. Gao, S. Tian, Y. Yang, N. Ma, Comparison of N-linked glycosylation on hemagglutinins derived from chicken embryos and MDCK cells: a case of the production of a trivalent seasonal influenza vaccine. Applied Microbiology and Biotechnology. 105, 3559–3572 (2021).

155. Y. J. Liu, S. L. Wu, K. R. Love, W. S. Hancock, Characterization of Site-Specific Glycosylation in Influenza A Virus Hemagglutinin Produced by Spodoptera frugiperda Insect Cell Line. Analytical Chemistry. 89, 11036–11043 (2017).

156. J. R. Schnell, J. J. Chou, Structure and mechanism of the M2 proton channel of influenza A virus. Nature 2008 451:7178. 451, 591–595 (2008).

157. Preparing large MD simulations with VMD, (available at https://www.ks.uiuc.edu/Research/vmd/minitutorials/largesystems/).

158. T. Williams, C. Kelley, C. Bersch, H.-B. Bröker, J. Campbell, R. Cunningham, D. Denholm, G. Elber, R. Fearick, C. Grammes, L. Hart, L. Hecking, P. Juhász, T. Koenig, D. Kotz, E. Kubaitis, R. Lang, T. Lecomte, A. Lehmann, J. Lodewyck, A. Mai, B. Märkisch, E. A. Merritt, P. Mikulík, D. Sebald, C. Steger, S. Takeno, T. Tkacik, J. van der Woude, J. R. van Zandt, A. Woo, J. Zellner, “gnuplot 5.4 An Interactive Plotting Program” (1986), (available at http://sourceforge.net/projects/gnuplot).

159. A. Shrake, J. A. Rupley, Environment and exposure to solvent of protein atoms. Lysozyme and insulin. Journal of Molecular Biology. 79, 351–371 (1973).

160. Hochbaum D, Shmoys D, A best possible heuristic for the k-center problem. Mathematics of Operations Research. 10, 180–184 (1985).

161. W. C. Swope, J. W. Pitera, F. Suits, Describing protein folding kinetics by molecular dynamics simulations. 1. Theory. Journal of Physical Chemistry B. 108, 6571–6581 (2004).

162. B. Trendelkamp-Schroer, H. Wu, F. Paul, F. Noé, Estimation and uncertainty of reversible Markov models. The Journal of Chemical Physics. 143, 174101 (2015).

163. R. Metzler, Generalized Chapman-Kolmogorov equation: A unifying approach to the description of anomalous transport in external fields. Physical Review E. 62, 6233 (2000).

164. B. S. Chapman, On the Brownian displacements and thermal diffusion of grains suspended in a non-uniform fluid. Proceedings of the Royal Society of London. Series A, Containing Papers of a Mathematical and Physical Character. 119, 34–54 (1928).

165. A. Kolmogoroff, Über die analytischen Methoden in der Wahrscheinlichkeitsrechnung. Mathematische Annalen. 104, 415–458 (1931).

166. R. T. McGibbon, K. A. Beauchamp, M. P. Harrigan, C. Klein, J. M. Swails, C. X. Hernández, C. R. Schwantes, L. P. Wang, T. J. Lane, V. S. Pande, MDTraj: A Modern Open Library for the Analysis of Molecular Dynamics Trajectories. Biophysical Journal. 109, 1528–1532 (2015).

167. J. E. Stone, K. L. Vandivort, K. Schulten, GPU-accelerated molecular visualization on petascale supercomputing platforms. Proceedings of the 8th International Workshop on Ultrascale Visualization, 6 (2013).

168. N. A. Doria-Rose, J. N. Bhiman, R. S. Roark, C. A. Schramm, J. Gorman, G.-Y. Chuang, M. Pancera, E. M. Cale, M. J. Ernandes, M. K. Louder, M. Asokan, R. T. Bailer, A. Druz, I. R. Fraschilla, N. J. Garrett, M. Jarosinski, R. M. Lynch, K. McKee, S. O’Dell, A. Pegu, S. D. Schmidt, R. P. Staupe, M. S. Sutton, K. Wang, C. K. Wibmer, B. F. Haynes, S. Abdool-Karim, L. Shapiro, P. D. Kwong, P. L. Moore, L. Morris, J. R. Mascola, New Member of the V1V2-Directed CAP256-VRC26 Lineage That Shows Increased Breadth and Exceptional Potency. J Virol. 90, 76–91 (2015).

169. T. Tiller, E. Meffre, S. Yurasov, M. Tsuiji, M. C. Nussenzweig, H. Wardemann, Efficient generation of monoclonal antibodies from single human B cells by single cell RT-PCR and expression vector cloning. J Immunol Methods. 329, 112–124 (2008).

170. D. N. Mastronarde, Automated electron microscope tomography using robust prediction of specimen movements. J Struct Biol. 152, 36–51 (2005).

171. S. H. W. Scheres, RELION: implementation of a Bayesian approach to cryo-EM structure determination. J Struct Biol. 180, 519–530 (2012).

172. E. F. Pettersen, T. D. Goddard, C. C. Huang, G. S. Couch, D. M. Greenblatt, E. C. Meng, T. E. Ferrin, UCSF Chimera--a visualization system for exploratory research and analysis. J Comput Chem. 25, 1605–1612 (2004).

173. S. F. Altschul, W. Gish, W. Miller, E. W. Myers, D. J. Lipman, Basic local alignment search tool. J Mol Biol. 215, 403–410 (1990).

