## Supporting Information for "Breathing and tilting: mesoscale simulations illuminate influenza glycoprotein vulnerabilities"

**This PDF file includes:**

Materials and Methods
Figs. S1 to S27
Tables S1 to S3
Captions for Movies S1 to S10
Captions for Data S1 to S2
References

**Other Supplementary Materials for this manuscript include the following:**

Movies S1 to S10
Data S1 to S2

### Materials and Methods

Mesoscale all-atom modeling of the influenza virus

In the work presented here, two glycosylated, whole-virion models of the H1N1 influenza A virus were built, simulated, and analyzed. The first corresponds to the strain A/swine/Shandong/N1/2009(H1N1) (hereafter referred to as H1N1-Shan2009), whereas the second one accounts for the A/45/Michigan/2015(H1N1) strain (hereafter referred to as H1N1-Mich2015). Both models are based on the unglycosylated H1N1-Shan2009 whole-virion model previously built by us and presented in Durrant et al. (*45*) The main difference lies in the addition of N-linked glycans to HA and NA glycoproteins, which substantially improved the biological relevance of the simulated constructs. Although we refer to Durrant et al. (*45*) for a detailed description of the original modeling of the individual protein, assembly procedure of the whole-virion model, and molecular dynamics (MD) simulations of both the individual proteins (HA, NA, M2) and the whole-virion model, a summary is also provided in the following paragraph.

The unglycosylated, H1N1-Shan2009 whole-virion model constructed and simulated by Durrant et al. (*45*) comprises 236 HA trimers, 30 NA tetramers, and 11 M2 ion channels embedded in a 3-palmitoyl-2-oleoyl-dglycero-1-phosphatidylcholine (POPC) quasi-spherical lipid bilayer (*45*). HA, NA and M2 were individually modeled and separately equilibrated (HA and NA) through MD simulations before being incorporated into the viral membrane (*45*). We note that in Durrant et al.(*45*) glycans were not modeled. LipidWrapper (*118*) was used to create the spherical lipid bilayer starting from a large planar POPC-bilayer model generated with the CHARMM-GUI server (*144*, *145*). As specified in ref. (*45*), slight differences in the leaflet densities are magnified by several times in the mesoscale systems, possibly creating large instabilities during simulation (*146*). Although a bilayer made of only POPC represents an oversimplification of the actual membrane composition (*147*), it has the advantage of minimizing such instabilities during simulations. This was the main reason behind the decision to build a POPC-only lipid bilayer. The M1 matrix proteins, coating the inner leaflet of the lipid bilayer, and the ribonucleoproteins contained inside of the virion were not modeled. As described in Amaro et al. (*115*), the shape of the virion and the distribution of the glycoproteins in the viral membrane were adopted from a point model of the virion exterior provided by collaborators, which was in turn derived from a cryo-electron tomography (cryoET) map of the influenza virus pleiomorphy (*83*). Positioning of the glycoproteins and insertion of M2 channels in the lipid bilayer according to the point model was done with PyMolecule (*148*, *149*). Full procedure, including handling of clashes, is described in refs. (*45*, *115*). The whole-virion model was solvated inside and outside of the lipid bilayer with explicit TIP3P water molecules (*129*), and ionized at 150 mM concentration with $Na^+/Cl^-$, tallying 160,653,271 atoms. All-atom MD simulations were performed on the Blue Waters supercomputer using NAMD2.10 (*130*) and CHARMM36 all-atom additive force fields for protein (*117*), lipids (*125*, *126*), and ions (*128*).

In the next paragraphs, we detail the system setup and all-atom MD simulations of the glycosylated H1N1-Shan2009 and H1N1-Mich2015 whole-virion models presented in this work.

System setup of the glycosylated H1N1-Shan2009 whole-virion model

UniProtKB accession codes for the sequences of HA and NA modeled in H1N1-Shan2009 are F2YI86 and F2YI87, respectively. HA was modeled in its uncleaved form (HA0), with residues ranging from 1 to 566. NA residues range from 1 to 469. We note that for HA and NA we adopted the absolute numbering system that starts with the first residue of the sequence in the corresponding UniProtKB entry. Sequences are reported in Fig. S2 and Fig. S3. The M2 construct (*45*) was based on the 2L0J structure (*150*), tallying 41 residues (from 22 to 62) since N-terminal and C-terminal tails were not modeled.

Initial coordinates for the setup of the glycosylated whole-virion model of H1N1-Shan2009 were derived from the last frame of the all-atom MD simulation of the unglycosylated whole-virion model previously performed by us and briefly described in the previous paragraph (*45*). Specifically, we used the last frame obtained from the third simulation branch, which is depicted in green in the schematic reported in the supplementary information file of ref. (*45*). At that point of the simulation, corresponding to 69.74 ns, lipid-bilayer instabilities had been already resolved and the system had been steadily simulated without major issues. Our goal was to improve the simulated system by adding N-linked glycans to HA and NA while preserving the relaxed, MD-derived morphology of the virion, thus facilitating a smooth continuation of the simulation endeavor. For this reason, we retained the MD-derived coordinates of all the atoms of the system, including water molecules and ions, and then performed glycosylation of HA and NA accordingly (Fig. S4). Specific oligo-mannose, hybrid, and complex N-linked glycans were identified to be linked to HA and NA as described in the following paragraphs.

Each H1N1-Shan2009 HA monomer exhibits six N-linked sequons (Fig. S5), of which one (N104) is located within the head and five (N28, N40, N293, N304, N498) within the stalk (*11*), for a total of 18 potential N-linked glycans per trimer. It must be noted that N27 is also a potential glycosylation site; however, as reported in previous studies (*11*, *124*), it is not glycosylated as N-X-T sequon at residue 28 is preferentially glycosylated over the N-X-S sequence at residue 27. N293 is located on the edge between the head and the stalk, but it is considered part of the stalk (*151*, *152*). Several glycomics studies were used as a reference to build the glycosylation profile of HA (*11*, *120–122*), which is shown in Fig. S5. Instead, each H1N1-Shan2009 NA monomer presents eight N-linked sequons (Fig. S5), of which four are located within the head (N88, N146, N235, N386) and four within the stalk (N50, N58, N63, N68) (*11*), for a total of 32 potential N-linked glycans per tetramer. Several glycomics studies were used as a reference to build the glycosylation profile of NA (*11*, *123*, *124*), which is displayed in Fig. S5.

Upon defining the respective glycosylation profiles, N-glycans were iteratively added to all the 236 HA trimers and 30 NA tetramers using the doGlycans tool (*119*). VMD (*136*), NAMD (*116*), GROMACS (*153*) and the tools therein contained were also used in the glycosylation workflow for pulling each glycoprotein from the virion, assessing sequon accessibility, renaming residues, local minimization, and parameterization. HA and NA N-linked sequons are overall all highly occupied (*120*, *154*, *155*). However, since glycans were added to a construct resulting from

69.74 ns of MD (*45*), many sequons occurred to be sterically hindered by adjacent residues or not fully exposed to the solvent as a result of the previous dynamics. Moreover, since the viral membrane was originally densely studded with unglycosylated HAs and NAs (*83*, *115*), adding glycans to sequons located at the interface between adjacent glycoproteins was often not possible. For these reasons, all the sequons were subjected to a preliminary screening to evaluate their accessibility. The accessible surface area (ASA) of the asparagine side chain's ND2 atom was calculated to determine whether a sequon was accessible in our initial model. Both scenarios, where HD21 or HD22 were removed from ND2, were considered (Fig. S4A). ASA was calculated with the *measure sasa* command implemented in VMD (*136*), combined with in-house scripts. A sphere of radius 10 Å, approximating the size of the basic $(GlcNAc)_2(Man)_3$ N-core present in all glycans (Fig. S4B), was used as a probe for the ASA screening. If the ASA of the putative asparagine's ND2 atom was larger than 10 $Å^2$, then a glycan would be tentatively linked (Fig. S4C). Clashes with surrounding amino acids or other glycans were subsequently examined and solved, first by performing a local minimization with NAMD2.13 (*116*) and then by eliminating the clashing glycan if the clashes could not be solved with minimization. Once the glycosylated glycoproteins were positioned back in the viral membrane, possible clashes arising between the glycans and the neighboring (glycosylated) glycoproteins were examined, resulting in the depletion of the clashing glycans. This further reduced the total number of glycans that were added. A total of 1521 glycans (29.2% of all the available sequons), among which 1277 were linked to HA (30.0%) and 244 to NA (25.4%) (Fig. S5). We note that we could only populate 29 out of the 708 available N104 sequons within the HA head. This low number is due to the overall limited accessibility, in our system, of residue N104, which is occluded by neighboring residues. A similar scenario occurred during modeling of HA's N104 glycosylation site is also reported in ref. (*11*).

Next, we adjusted the orientation of the 11 M2 ion channels within the lipid bilayer, positioning the N-ter end extravirion and the C-ter end intravirion (*5*, *156*), since they had been oriented flipped in the original system (*45*). No alterations in the composition and the coordinates of the POPC lipid bilayer were done with respect to the original system (*45*). The resulting glycosylated H1N1-Shan2009 virion construct, comprising 236 glycosylated HA homo-trimers, 30 glycosylated NA homo-tetramers, and 11 M2 homo-tetramers embedded in the POPC lipid bilayer, was parameterized using NAMD's PSFGEN (*116*) and CHARMM36 all-atom additive force fields for protein (*117*), lipids (*125*, *126*), glycans (*127*), and ions (*128*). As mentioned above, explicit TIP3 water molecules (*129*) located inside and outside of the lipid bilayer, $Na^+$ and $Cl^-$ ions, as well as $Ca^{2+}$ ions bound to the NA heads, were retained from the original unglycosylated system (*45*). The addition of glycans altered the overall electric charge of the solute, thus requiring tweaking of the total number of $Na^+$ and $Cl^-$ ions to ensure electrical neutrality. These modifications did not impact the overall ionic strength, which remained at ~150 mM. The following minimization steps solved eventual clashes between newly added glycans and pre-existing water molecules and ions. The total number of atoms for the final system is 160,919,594, with an orthorhombic periodic cell of 1142 Å × 1194 Å × 1155 Å. After generating suitable files for mesoscale simulations with NAMD (*157*), the system was ready to undergo all-atom MD simulations.

System setup of the glycosylated H1N1-Mich2015 whole-virion model

UniProtKB accession codes for the sequences of HA and NA modeled in H1N1-Mich2015 are A0A144YDV8 and A0A0X9QTS2, respectively. HA was modeled in its uncleaved form (HA0), with residues ranging from 1 to 566. NA residues range from 1 to 469. We note that for HA and NA, we adopted the absolute numbering system that starts with the first residue of the sequence in the corresponding UniProtKB entry. Sequences are reported in Fig. S2 and Fig. S3. The M2 construct (*45*) was based on the 2L0J structure (*150*), tallying 41 residues (from 22 to 62) since N-terminal and C-terminal tails were not modeled.

Building on the same initial coordinates obtained from the MD simulation of the unglycosylated virion (*45*), the system setup for H1N1-Mich2015 was conducted in the same way as described for H1N1-Shan2009. The main challenge was introducing the H1N1-Mich2015 point mutations occurring in HA and NA. All the mutations within HA (15 per monomer) and NA (14 per monomer) were modeled onto the initial construct using NAMD's PSFGEN (*116*) *mutate* command and the CHARMM36 amino acid topologies to build side chain rotamers (*117*). The mutations are listed and displayed in Fig. S6. No mutations were introduced to M2 ion channels, which were treated and positioned in the lipid bilayer in the same way as in H1N1-Shan2009. The resulting, mutated models of HA and NA were then subjected to the same glycosylation procedure adopted for H1N1-Shan2009 and described above.

S179N mutation introduced a second glycan within the HA head (Fig. S7), in addition to the already existent N104. Therefore, each H1N1-Mich2015 HA monomer exhibits seven N-linked sequons (Fig. S8), of which two (N104, N179) are located within the head and five (N28, N40, N293, N304, N498) within the stalk (*11*), for a total of 21 potential N-linked glycans per trimer. The same glycoprofile as in H1N1-Shan2009 was adopted (*11*, *120–122*). The new glycan linked to N179 was modeled as a complex tri-antennary (Fig. S8). N386K mutation ablated the N386 sequon within the NA head (Fig. S7). Therefore, each H1N1-Mich2015 NA monomer presents seven N-linked sequons (Fig. S8), of which three are located within the head (N88, N146, N235) and four within the stalk (N50, N58, N63, N68) (*11*), for a total of 28 potential N-linked glycans per tetramer. The same glycoprofile as in H1N1-Shan2009 was adopted except for the deleted N386 glycan (Fig. S8) (*11*, *123*, *124*).

A total of 1791 glycans (30.9 % of all the available sequons) were added to H1N1-Mich2015, among which 1626 were linked to HA (32.8%) and 165 to NA (19.6%) (Fig. S8). We note that also for H1N1-Mich2015 we could populate a low number of N104 sequons within the HA head (33 of 708). Instead, we could populate 351 sequons out of 708 available (49.6%) for the new N179 glycan within the HA head (Fig. S8).

Similar to H1N1-Shan2009, the resulting glycosylated H1N1-Mich2015 virion construct was parameterized using NAMD's PSFGEN (*116*) and CHARMM36 all-atom additive force fields for protein (*117*), lipids (*125*, *126*), glycans (*127*), and ions (*128*). As for H1N1-Shan2009, explicit

TIP3P water molecules (*129*) located inside and outside of the lipid bilayer, $Na^+$ and $Cl^-$ ions, as well as $Ca^{2+}$ ions bound to the NA heads, were retained from the original unglycosylated system (*45*). The amino acid mutations and the addition of glycans altered the overall electric charge of the solute, thus requiring further tweaking of the total number of $Na^+$ and $Cl^-$ ions to establish electrical neutrality. These modifications did not impact the overall ionic strength, which remained at ~150 mM. The following minimization steps solved eventual clashes involving mutated residues or between newly added glycans and pre-existing water molecules and ions. The total number of atoms for the final system is 160,981,954, with an orthorhombic periodic cell of 1138 Å × 1189 Å × 1150 Å. After generating suitable files for mesoscale simulations with NAMD (*157*), the system was ready to undergo all-atom MD simulations.

<u>All-atom molecular dynamics simulations</u>

All-atom MD simulations of H1N1-Shan2009 and H1N1-Mich2015 were performed on the TITAN supercomputer at Oak Ridge National Laboratory (ORNL) and the NSF Blue Waters supercomputer at the University of Illinois at Urbana-Champaign, respectively, using the CUDA memory-optimized version of NAMD 2.13 (*130*). The same simulation protocol was adopted for both H1N1-Shan2009 and H1N1-Mich2015. CHARMM36 all-atom additive force fields were used to describe proteins (*117*), lipids (*125*, *126*), glycans (*127*), and ions (*128*). Simulations were performed in explicit water using the TIP3P model (*129*).

Preliminary tests were initially run to assess the stability of the systems, familiarize with the computing facilities, and debug possible errors. Then, the systems were initially minimized in two consecutive cycles of 15,000 steps, each using the NAMD's conjugate gradient energy approach. During the first cycle, the position of protein backbone atoms and of POPC P atoms within 15 Å of the M2 ion channels was kept harmonically restrained at 100 kcal/mol/Å$^2$. We note that the orientation of M2 ion channels was modified with respect to our previous work (*45*), thus requiring more caution. We also remark that the initial coordinates for water molecules, ions, lipids, and proteins (except for M2) were retained from a previous MD simulation as described above (*45*), thus being already equilibrated. During the second minimization cycle, the harmonic restraints on the protein backbone atoms were released, whereas the harmonic restraints on the POPC P atoms within 15 Å of M2 ion channels were decreased to 10 kcal/mol/Å$^2$. After minimization, the systems were gradually heated up with no restraints from 0 K to 298 K in 220 ps using isothermal–isobaric conditions (NPT ensemble). The temperature was controlled using the Langevin thermostat (*131*) with a damping coefficient of 5/ps, whereas the Nosé-Hoover Langevin piston (*132*, *133*) was used to maintain the pressure at 1.01325 bar. A time-step of 2 fs was used. Next, the systems were equilibrated for 0.5 ns under the NPT ensemble. Temperature (298 K) and pressure (1.01325 bar) control were achieved again using the Langevin thermostat (*131*) with a damping coefficient of 5/ps, and the Nosé-Hoover Langevin piston (*132*, *133*). During this phase, harmonic restraints at 100 kcal/mol/Å$^2$ were applied again to POPC P atoms within 15 Å of M2 ion channels and gradually released to 10 kcal/mol/Å$^2$.

Finally, the systems were submitted to productive MD simulations in NPT conditions using the same barostat and thermostat as above to achieve pressure (1.01325 bar) and temperature (298 K) control. Langevin damping coefficient was set to 1/ps during productive MD. One continuous replica was performed for each system using a time-step of 2 fs. To preserve the overall geometry of the virion models in the absence of the interior component, we applied a harmonic position restraint of 20 kcal/mol/Å$^2$ to every 20$^{th}$ inner-leaflet P atom. A similar protocol was adopted in our previous virion simulation (*45*), where instead every 10$^{th}$ POPC head group was kept fixed. Non-bonded interactions (van der Waals and short-range electrostatic) were calculated at each timestep using a cutoff of 12 Å and a switching distance of 10 Å. All simulations were performed using periodic boundary conditions, employing the particle-mesh Ewald method (*134*) with a grid spacing of 2.1 Å to evaluate long-range electrostatic interactions every three timesteps. SHAKE algorithm (*135*) was adopted to keep the atomic bonds involving hydrogens fixed with the NAMD option *rigidbonds all*. Frames were saved every 30,000 steps (60 ps).

For H1N1-Shan2009, we collected a total of ~441.78 ns (7363 frames) continuous productive MD (Movie S1). Simulations were performed with the CUDA memory-optimized version of NAMD 2.13 (*130*) running on 4,096 physical nodes (28,672 processors) on the TITAN supercomputer, with an average benchmark of ~13.83 ns/day. A total of ~17 TB of data were generated, including stripped trajectories.

For H1N1-Mich2015, we collected a total of 424.98 ns (7083 frames) continuous productive MD (Movie S2). Simulations were performed with the CUDA memory-optimized version of NAMD 2.13 (*130*) running on 4,096 physical nodes (114,688 processors) on the Blue Waters supercomputer, with an average benchmark of ~13.78 ns/day. A total of ~16 TB of data was generated including stripped trajectories.

Root Mean Squared Deviation (RMSD) analysis of HA and NA

For both strains, each of the 236 HA trimeric and 30 NA tetrameric trajectories was separately aligned using as a reference the respective initial coordinates of all the backbone atoms (C, N, O, Cα). Then, for each of the HA trimers/NA tetramers, we calculated the RMSD of the backbone atoms using the *measure rmsd* command implemented in VMD (*136*). Subsequently, we computed the mean and standard deviation of the RMSD values obtained at each frame across all the 236 HA trimers / 30 NA tetramers, generating, respectively, the averaged RMSD profiles for HA and NA (Fig. S9).

Root Mean Squared Fluctuation (RMSF) analysis of HA and NA residues

We performed the calculation of Cα-atom RMSF to probe the average mobility of HA and NA residues. Before the calculation of Cα-atom RMSF, rotation and translation motions of each individual HA trimer and NA tetramer were removed through targeted, specific alignments. In detail, two sets of aligned trajectories, based on different residue masks, were generated for each of the 236 HA trimers and 30 NA tetramers of both strains. For each of the HA trimers, the first set of alignment was performed using as a reference the respective initial coordinates of all the backbone atoms (C, N, O, Cα) of the ectodomain region (residues 18 to 503) of the respective glycoprotein, whereas the second set was based on the region comprising the transmembrane domain (TMD) (residues 529 to 549). The former served for the calculation of RMSF of residues 1 to 503, the latter for the calculation of RMSF of residues 504 to 566. For each of the NA tetramers, the first set of alignment was performed using as a reference the respective initial coordinates of all the backbone atoms (C, N, O, Cα) of the head region (residues 96 to 461) of the respective glycoprotein, whereas the second set was based on the region comprising the stalk and the TMD (residues 7 to 81). The former served for the calculation of RMSF of residues 96 to 469, the latter for the calculation of RMSF of residues 1 to 95. This procedure was performed due to peculiar structures of HA and NA that present domains undergoing tilt motions during the dynamics, i.e., HA ectodomain tilting and NA head tilting. If we had performed a single alignment based on the backbone atoms of the whole structure, the extensive motions of the HA ectodomain and NA head, which are larger in size and account for most of the residues, would have biased the RMSF values of the complementary region encompassing the hinge linker (both for HA and NA), the stalk (for NA), the TMD, and the cytosolic tail.

For each trajectory of both sets and for each monomer, we calculated the RMSF of the Cα atoms of the residues belonging to the respective region using the *measure rmsf* command implemented in VMD (*136*). All the frames were used for the calculation. We then combined the RMSF values obtained for the two regions into a single RMSF profile for each monomer. Finally, we computed the mean and standard deviation of the RMSF values obtained for each Cα atom across all the 708 HA monomers / 120 NA monomers, generating the respectively averaged RMSF profiles for HA and NA (Fig. S10).

Positional fluctuations of HA's and NA's TMD and ectodomain center of masses

For both H1N1-Shan2009 and H1N1-Mich2015, we calculated the RMSF of HA's and NA's TMD center of mass (COM) and ectodomain COM using the first frame as a reference. The analysis was performed using in-house tcl scripts within the VMD framework (*136*). We note that the aim of this analysis was estimating the average positional fluctuations of the TMD COM and ectodomain COM of each glycoprotein within the virion, therefore only the global rotation and translation motions of the whole virion were removed and not those of each individual glycoprotein within the virion. Global rotation and translation motions of the whole virion coordinates during dynamics were removed by aligning the trajectory onto the initial coordinates of all the protein backbone atoms (C, N, O, Cα). The COM of HA TMD was defined using the coordinates of backbone atoms of residues 529 to 549. The COM of HA ectodomain was defined using the coordinates of backbone atoms of residues 1 to 528. The COM of NA TMD was defined using the coordinates of backbone atoms of residues 7 to 29. The COM of NA ectodomain was defined using the coordinates of backbone atoms of residues 30 to 469.

At each frame of the whole virion trajectory, we calculated, for each glycoprotein, the squared positional fluctuation of both COMs with respect to their initial position (reference). Next, always for each glycoprotein, we averaged the squared fluctuations of the COMs and calculated the square root of the averages, obtaining the respective RMSF value.

NA head-tilt angle

The NA head-tilt angle was calculated for all 30 NA tetramers at each frame of both virion trajectories using in-house tcl scripts within the VMD framework (*136*). The NA head-tilt angle is defined by the principal axis of the tetrameric upper stalk region (residues 50 to 78) and the vector joining residue 78 at the top of the stalk with the COM of the tetrameric head (residues 91 to 469). The distribution of NA head-tilt angle values was computed through kernel density estimation using Gaussian kernels with gnuplot plotting program (*158*).

HA ectodomain-tilt angle

The HA ectodomain-tilt angle was calculated for all 236 HA trimers at each frame of both virion trajectories using in-house tcl scripts within the VMD framework (*136*). The HA ectodomain-tilt angle is defined by the principal axis of the trimeric TMD (residues 529 to 549) and the vector joining residue 529 at the top of the TMD with the COM of the trimeric ectodomain's long α-helices (LαHs) (residues 419 to 471). The distribution of HA ectodomain-tilt angle values was computed through kernel density estimation using Gaussian kernels with gnuplot plotting program (*158*).

HA head breathing

The extent of HA head breathing was estimated by calculating the distance between the COM of each monomeric HA head (residues 66 to 286) and the COM of the apical residues of the three LαHs. The apical residue of each LαH corresponds to residue 419 and the COM was calculated based on the three residues 419. This distance was calculated for all 708 HA monomers at each frame of both virion trajectories using in-house tcl scripts within the VMD framework (*136*). The distribution of HA head–LαHs distance values was computed through kernel density estimation using Gaussian kernels with gnuplot plotting program (*158*).

NA head breathing

The extent of NA head breathing was estimated by calculating the inter-protomer distances between the four NA head COMs (residues 96 to 461). Considering the four NA heads within each tetramer, a total of six distances have been monitored for each NA tetramer (Fig. S27), at each frame of both virion trajectories, using in-house tcl scripts within the VMD framework (*136*). The distribution of the values of the inter-protomer distances was computed through kernel density estimation using Gaussian kernels with gnuplot plotting program (*158*).

FluA-20 epitope accessible surface area (ASA)

Accessible surface area (ASA) of the FluA-20 (*39*) epitope, located on the HA head domain trimer interface, was calculated using the *measure sasa* VMD command (*136*), an implementation based on the Shrake and Rupley algorithm (*159*), along with in-house tcl scripts to handle data. The following HA residues were considered for the epitope: residues 109, 230, 232-237, 243. An 18.6 Å probe radius, which approximately encloses the FluA-20 Fab's variable domains (Fig. 5 in the main text), was used. ASA evaluations were conducted on the HA trimer showing the highest degree of head domain breathing, at every frame of the trajectory.

Evaluation of glycoprotein–glycoprotein interactions

For both virion simulations, interactions between glycoproteins were evaluated using the *measure contacts* implemented in VMD (*136*) along with in-house tcl scripts to handle data.

For each glycoprotein, we first pre-computed a list of neighbor glycoproteins using a cutoff of 75 Å. Then, always for each glycoprotein, we assessed the number of connections formed with any of the glycoproteins included in its respective neighbor list every $10^{th}$ frame of the virion trajectory. At each evaluation, a "connection" was considered formed and counted as one when any protein or glycan heavy atoms of the screened glycoprotein were detected to be within 5 Å of any protein or glycan heavy atoms of the screened neighbor glycoprotein. Only heavy atoms belonging to the ectodomains were considered for the analysis. Importantly, if glycoprotein "A" was, for example, found to form a connection with glycoprotein "B", that connection was counted one time for glycoprotein "A" and one time also for glycoprotein "B". Relative fractions of glycoproteins forming respectively 0, 1, 2, 3, 4, or 5 connections at each evaluation were subsequently computed with respect to the total number of glycoproteins (266). Results are shown in Fig. 7 for all the glycoproteins, i.e., HAs + NAs, in the main text and in Fig. S24 for HA and NA.

Within the same workflow, we then evaluated the formation of macro-clusters of glycoproteins. At each assessment, we calculated the number of glycoproteins that were part of a macro-cluster using the same cutoff of 5 Å between protein/glycan heavy atoms. If glycoprotein "A" was, for example, found to form a connection with glycoprotein "B," and, glycoprotein "B" was in turn found to establish at the same time a connection with glycoprotein "C," the three glycoproteins A, B, and C were considered as part of a macro-cluster of 3, regardless of the presence of a direct connection between A and C. Results are shown in Fig. S25.

Next, we evaluated the nature of the connections formed by each glycoprotein. The connections were broken down into three categories depending on the residues used by each glycoprotein to form such connections: i) "glycan-only" if only glycan residues were used to form the connection; ii) "protein-only" if only protein residues were used to form the connection; "protein and glycan" if both protein and glycan residues were used at the same time to interact with the screened neighbor glycoprotein. Relative frequencies of "glycan-only", "protein-only", and "protein and glycan" types of connection were subsequently computed with respect to the total number of counted connections. Results are shown in Fig. 7 in the main text.

Finally, if glycans were involved in a connection between two glycoproteins, for each glycoprotein we counted the number of stalk/head glycans involved in the connection, respectively. Relative frequencies of the interacting stalk/head glycans were subsequently calculated for HA and NA with respect to the respective total number of stalk/head glycans in both strains. Results are shown in Fig. S26.

Markov State Models (MSMs)

The first step was the preparation of individual NA tetrameric (30) and HA trimeric (236) trajectories, which were generated from the whole-virion MD simulations of H1N1-Shan2009 and H1N1-Mich2015, respectively. Glycans were not included in the generated trajectories because a common topology, i.e., equal for all the NA tetramers and HA trimers, respectively, was necessary to load all the trajectories in the MSM workflow. However, the impact of glycans on the glycoproteins remains accounted for as the trajectories were generated from the simulations of the glycosylated whole-virions.

*NA head tilt.* The MD simulation trajectories were imported into the PyEMMA2 software (*137*) for MSM analysis. The NA head tilt MSM was created as follows. We used two features; the first was defined as the shortest distance between the COM of the head and the COM of the center of the stalk, while the second feature was defined as the shortest distance between the COM of the head and the COM of the bottom of the stalk. The exact residues defining these COM selections are given in the accompanying Jupyter notebook (Data S1). With these two features, we ran a time-lagged independent component analysis (TICA) as is commonly used on MSMs with multiple features (*138–140*) using a lag time of 100 steps. Our step size is 0.06, so 100 steps are equivalent to 6 ns. We ran k-means clustering (*160*) of the two features, selecting 300 clusters. From our implied timescale plots (*161*, *162*) we selected an MSM lag time of 250 steps (Fig. S12). We then created Bayesian Markov state models, selecting two macrostates for each MSM. Using these parameters, we created MSMs and validated our two-state systems through Chapman-Kolmogorov (CK) tests (*142*, *163–165*) at a confidence interval of 95% (Fig. S13). The estimated and predicted lines overlapped out to 750 steps (3x the MSM lag time number), which meant these MSMs are statistically valid (*142*). We then used Robust Perron Cluster Center Analysis (PCCA++) to cluster the first two eigenvectors of the MSM transition matrices to cluster the states (*143*). The first macrostate in the H1N1-Shan2009 strain MSM (untilted) had a stationary distribution of 51.4% of the total population, while the second macrostate (tilted) had a distribution of 48.6%. The first macrostate in the H1N1-Mich2015 strain MSM (untilted) had a stationary distribution of 44.3% of the total population, while the second macrostate (tilted) had a distribution of 55.7%. Finally, we extracted NA structures from the two macrostates per strain by creating a 99.99999% probability cutoff for the H1N1-Shan2009 strain macrostates and a 99.999999% probability cutoff for the H1N1-Mich2015 strain macrostates. This means that the NA structures (microstates) that we selected have a >99.99999% probability of residing in the macrostate state that we think they are in through the PCCA++ fuzzy spectral clustering method. Confirmation of which microstates corresponded to which conformations was performed through the MDTraj software program (*166*).

*HA head tilt.* The MD simulation trajectories were imported into the PyEmma2 software (*137*) for MSM analysis. The HA head tilt MSM was created as follows. We used two features, defined as the distance between the COM of the head and the center of the HA stalk, and the distance between the HA monomer head tips and the bottom of the stalk. The exact residues defining these COM selections are given in the accompanying Jupyter notebook. With these two features, we ran TICA (*138–140*) using a lag time of 25 steps. Our step size is 0.06, so 25 steps are equivalent to 1.5 ns. We ran k-means clustering (*160*) of the three features, selecting 300 clusters. From our implied timescale plots (*161*, *162*) we selected an MSM lag time of 50 steps (Fig. S17). We then created Bayesian Markov state models, selecting two macrostates for each MSM. Using these parameters, we created MSMs and validated our two-state systems through CK tests (*142*, *163–165*) at a confidence interval of 95% (Fig. S18). The estimated and predicted lines overlapped out to 150 steps (>3x the MSM lag time number), which meant these MSMs are statistically valid (*142*). We then used PCCA++ to cluster the first two eigenvectors of the MSM transition matrices to cluster the states (*143*). The first macrostate in the H1N1-Shan2009 strain MSM (tilted) had a stationary distribution of 6.3% of the total population, while the second macrostate (tilted) had a distribution of 93.7%. The first macrostate in the H1N1-Mich2015 strain MSM (tilted) had a stationary distribution of 6.2% of the total population, while the second macrostate (untilted) had a distribution of 93.8%. Finally, we extracted HA structures from the two macrostates per strain by creating a 99.999999% probability cutoff all macrostates. This means that the HA structures (microstates) that we selected have a 99.999999% probability of residing in the macrostate state that we think they are in through the PCCA++ fuzzy spectral clustering method. Confirmation of which microstates corresponded to which conformations was performed through the MDTraj software program (*166*).

*HA head breathing*. The MD simulation trajectories were imported into the PyEMMA2 software (*137*) for MSM analysis. The HA head breathing MSM was created as follows. We used one feature, defined as the smallest mean distance of the monomers to each other. The exact residues defining this feature are given in the accompanying Jupyter notebook. With this one feature, we ran TICA (*138–140*) using a lag time of 100 steps to properly format the data, even though the one dimension cannot be reduced, obviously. Our step size is 0.06, so 100 steps are equivalent to 6 ns. We ran k-means clustering (*160*) of the features, selecting 300 clusters. Running TICA and k-means clustering should not be necessary for an MSM with one feature, but we ran these anyways for continuity. From our implied timescale plots (*161*, *162*) we selected an MSM lag time of 500 steps (Fig. S20). We then created Bayesian Markov state models, selecting two macrostates for each MSM. Using these parameters, we created MSMs and validated our two-state systems through CK tests (*142*, *163–165*) at a confidence interval of 95% (Fig. S21). The estimated and predicted lines overlapped out to 1250 steps (>2.5x the MSM lag time number), which meant these MSMs are statistically valid (*142*). We then used PCCA++ to cluster the first two eigenvectors of the MSM transition matrices to cluster the states (*143*). The first macrostate in the H1N1-Shan2009 strain MSM (open) had a stationary distribution of 24.6% of the total population, while the second macrostate (closed) had a distribution of 75.4%. The first macrostate in the H1N1-Mich2015 strain MSM (open) had a stationary distribution of 26.7% of the total population, while the second macrostate (closed) had a distribution of 73.3%. Finally, we extracted HA structures from the two macrostates per strain by creating a 99.99999% probability cutoff for the H1N1-Shan2009 strain macrostates and a 99.9999% probability cutoff for the H1N1-Mich2015 strain macrostates. This means that the HA structures (microstates) that we selected have a >99.9999% probability of residing in the macrostate state that we think they are in through the PCCA++ fuzzy spectral clustering method. Confirmation of which microstates corresponded to which conformations was performed through the MDTraj software program (*166*).

Figures and movies of the whole-virion simulations

Figure panels and movies of the whole-virion simulations were rendered from the trajectories using VMD GPU-accelerated ray tracing (*136*, *167*).

Single-cell sorting, immunoglobulin amplification, sequencing and mAb production

Cryopreserved PBMC samples were thawed and stained with monoclonal antibodies against human CD3, CD14, CD56, CD19, CD21, CD27, CD38, IgM and IgG (BioLegends, BD BioSciences, Beckam Coulter). The NA probe N2 (A/Wisconsin/67/2005) was conjugated with either AX488 or AX647 fluorochrome according to the manufacturer's protocol (Microscale Protein Labeling Kit, Thermo Fisher Scientific) prior to use in flow cytometry. Non-B cells and dead cells were excluded with CD3, CD14, CD56, and aqua (live-dead) staining in the same fluorescent channel. Memory B cells were further gated on CD38low, CD27high, IgM- and IgG+ and NA-binding memory B cells were single-cell sorted into a 96-well plate (Bio-Rad) using an FACS Aria Instrument (BD Immunocytometry Systems). Plates were subjected to reverse transcription PCR and cDNA were used for PCR to amplify immunoglobulin (Ig) heavy and light chain genes as described previously (*168*, *169*). PCR products were sequenced by Sangar sequencing. Ig heavy and light chain sequences were synthesized and cloned into IgG1 heavy and kappa backbone expression vector (GenScript). Antibody NDS.1 was expressed recombinantly by transient transfection of Ig expression plasmids in Expi293 cells with ExpiFectamine (Thermo Fisher Scientific). Supernatant was harvested 5 days post transfection and antibody was purified by using Protein A Sepharose (Thermo Fisher Scientific).

Biolayer interferometry (BLI)

BLI experiments were performed by using the Octet HTX instrument (Sartorius). HIS1K biosensors (Sartorius) were hydrated in PBS prior to use. Recombinant N2 NA tetramers derived from A/Wisconsin/67/2005 and A/Darwin/9/2021 were immobilized on HIS1K biosensors through their hexahistidine tags. After brief equilibration in assay buffer (25 mM Tris pH 8.0, 150 mM NaCl, 1% BSA), the biosensors were dipped into a two-fold dilution series of NDS.1 Fab for 5 min. Starting concentration of NDS.1 was 800 nM. Biosensors were then dipped in the assay buffer to allow Fab to dissociate from NA for 10 min. All assay steps were performed at 30°C with agitation set at 1,000 rpm. Baseline correction was carried out by subtracting the measurements recorded for a sensor loaded with the NA in the same buffer with no Fab. Data analysis and curve fitting were done with the Octet analysis software (version 11). Experimental data were fitted with the binding equations describing a 1:1 (Langmuir model) interaction.

Negative-stain electron microscopy

Samples were diluted to 0.02 mg/ml with buffer containing 10 mM HEPES, pH 7.0, and 150 mM NaCl, adsorbed to glow-discharged carbon-coated copper grids for 10 s, washed with three drops of the same buffer and negatively stained with 0.75% uranyl formate. Micrographs were recorded at a nominal magnification of 100,000 (corresponding to a pixel size of 2.2 Å) on an FEI Tecnai T20 electron microscope equipped with a 2k x 2k Eagle CCD camera and operated at 200 kV. Data collection was performed with SerialEM (*170*). Particles were selected from micrographs using in-house written software (YT, unpublished) and extracted into 144 x 144-pixel boxes. After 2D classification, initial models were obtained and refined using Relion 3.1 (*171*) with C4 symmetry enforced. The final datasets contained 2,304 and 4,368 particles for NDS.1 Fab and NDS.1/1G01 Fabs, respectively. The corresponding resolutions determined using the Fourier shell correlation threshold of 0.5 was 24.1 and 24 Å. UCSF Chimera (*172*) was used for docking and map visualization.

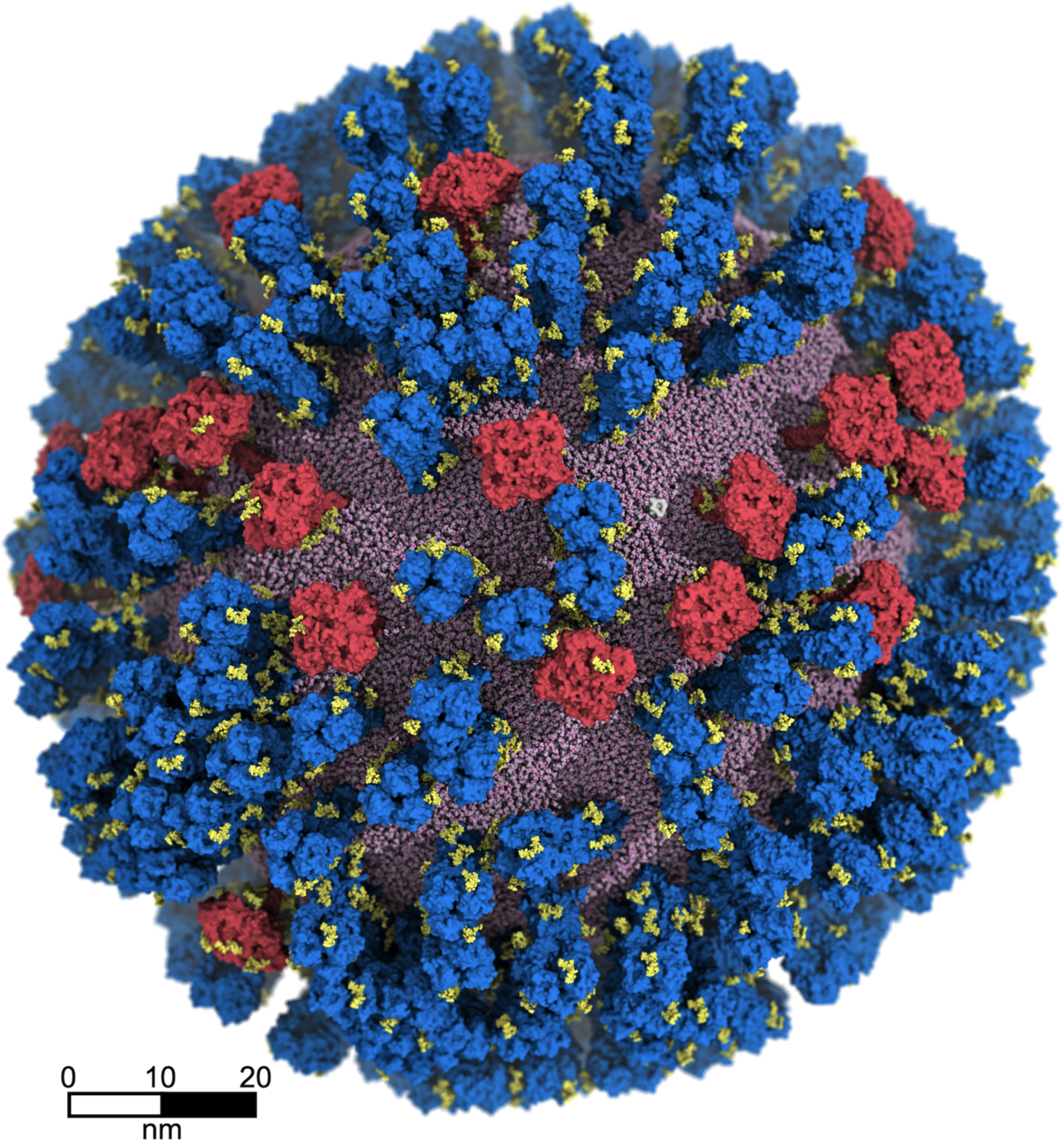


**Fig. S1.** All-atom glycosylated model of H1N1-Mich2015 influenza A virion. Neuraminidase (NA) and hemagglutinin (HA) glycoproteins, and the M2 ion channels, are depicted with red, blue, and white surfaces, respectively. N-linked glycans are shown in yellow. The spherical lipid envelope is represented with pink van der Waals (vdW) spheres.

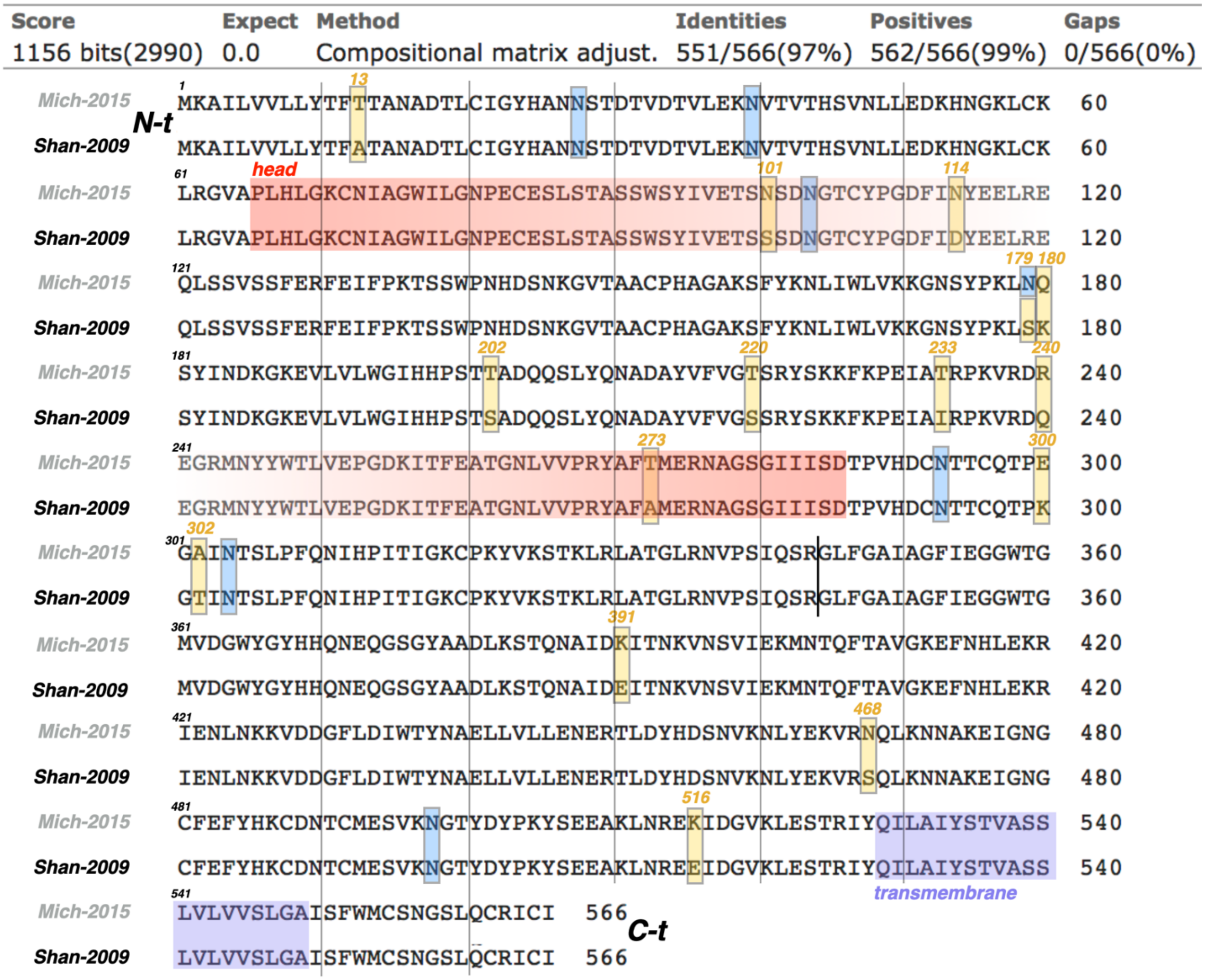


**Fig. S2.** Aligned sequences of HA in H1N1-Shan2009 (UniProtKB accession code: F2YI86) and H1N1-Mich2015 (UniProtKB accession code: A0A144YDV8) as modeled in this work. N-linked glycosylation sites are highlighted with blue squares, whereas mutation sites with yellow squares. Alignment was performed with BLAST (*173*).

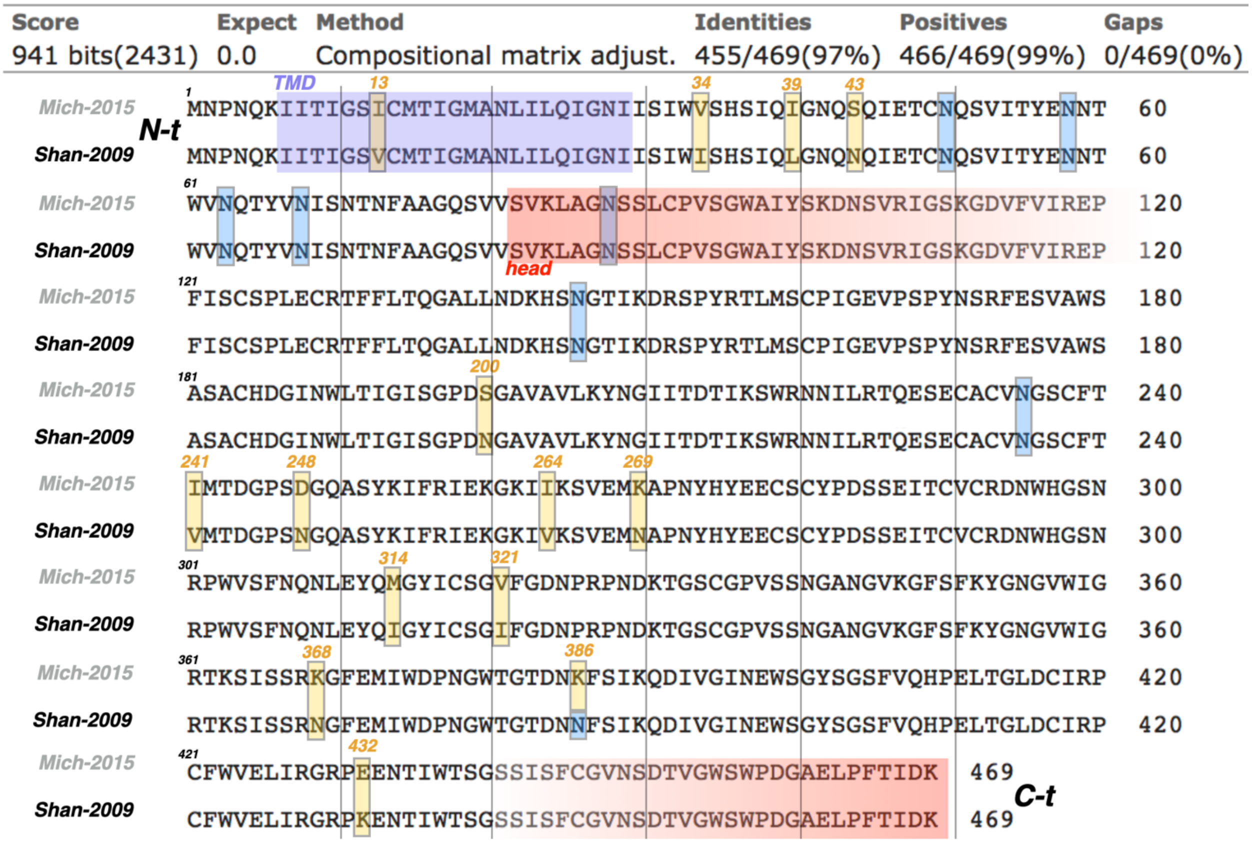

**Fig. S3.** Aligned sequences of NA in H1N1-Shan2009 (UniProtKB accession code: F2YI86) and H1N1-Mich2015 (UniProtKB accession code: A0A0X9QTS2) as modeled in this work. N-linked glycosylation sites are highlighted with blue squares, whereas mutation sites with yellow squares. Alignment was performed with BLAST (*173*).

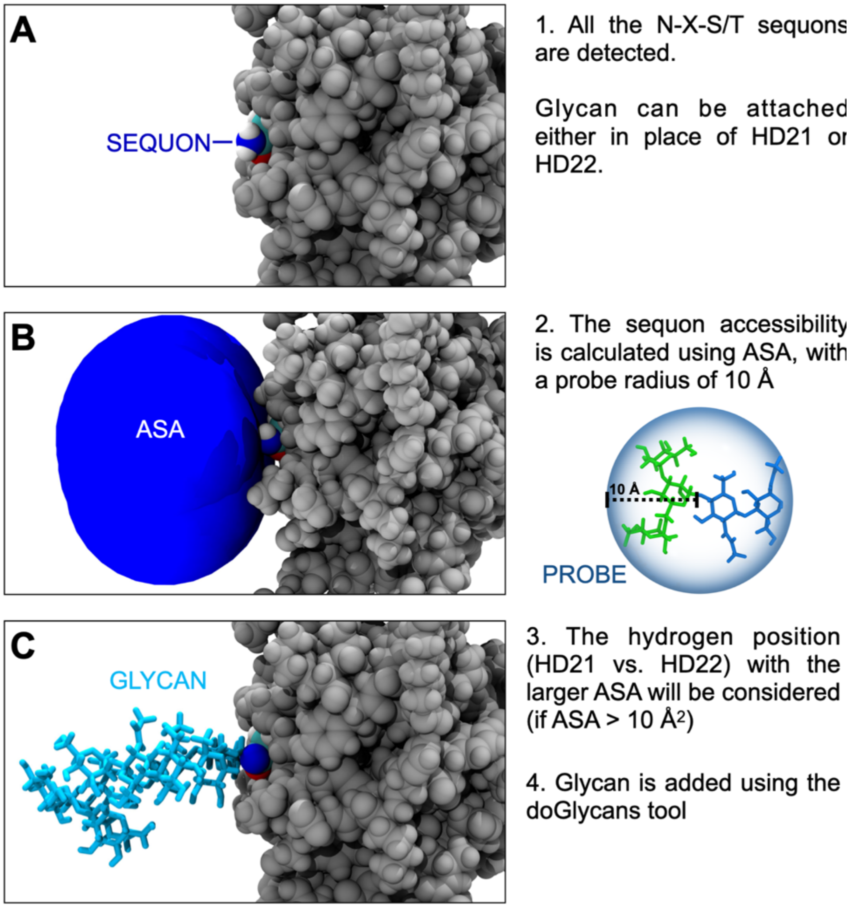


**Fig. S4.** Schematic representation of the glycosylation procedure. **A**) N-X-S/T N-linked sequons are selected. **B**) Although all sequons are overall highly occupied as reported by previous studies (*120*, *154*, *155*), in this work we evaluated the accessibility of each selected sequon by calculating the accessible surface area (ASA) of the asparagine side chain's ND2 atom using a probe radius of 10 Å. This was done since the initial coordinates of the whole-virion construct are derived from previous MD simulation of the unglycosylated whole-virion construct (*45*). **C**) Addition of glycan.

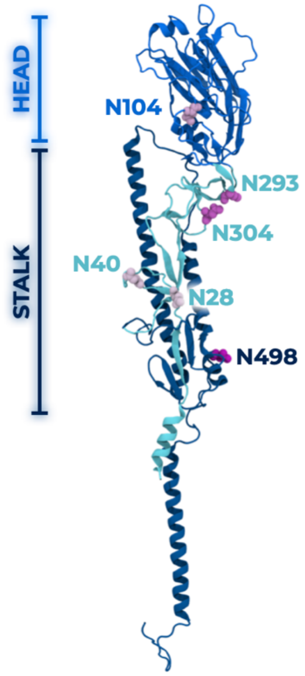


## HA

| Sequon | Region | Structure | # | % |
|---|---|---|---|---|
| N28 | Stalk | 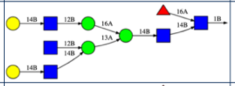 | 428/708 | 60.5% |
| N40 | Stalk | 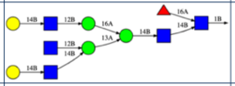 | 299/708 | 42.2% |
| N104 | Head | 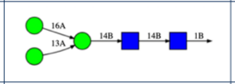 | 29/708 | 4.1% |
| N293 | Stalk | 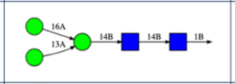 | 36/708 | 5.1% |
| N304 | Stalk | 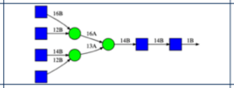 | 95/708 | 13.4% |
| N498 | Stalk | 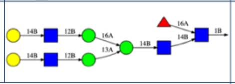 | 390/708 | 55.1% |
| TOTAL | | | 1277/4248 | 30% |

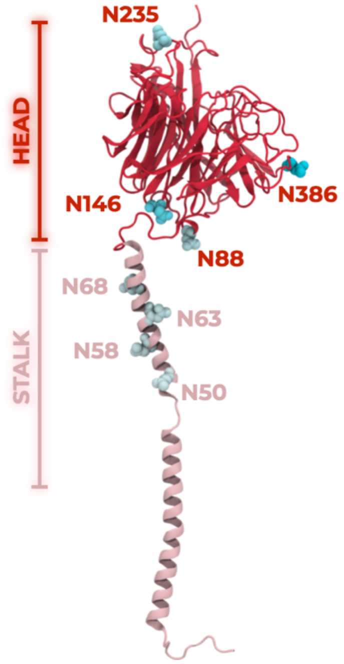


## NA

| Sequon | Region | Structure | # | % |
|---|---|---|---|---|
| N50 | Stalk | 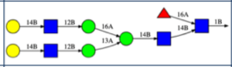 | 30/120 | 25.0% |
| N58 | Stalk | 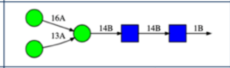 | 0/120 | 0% |
| N63 | Stalk | 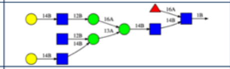 | 38/120 | 31.7% |
| N68 | Stalk | 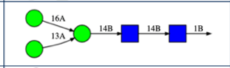 | 10/120 | 8.3% |
| N88 | Head | 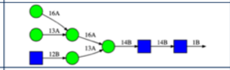 | 37/120 | 30.8% |
| N146 | Head | 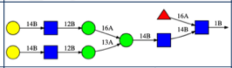 | 16/120 | 13.3% |
| N235 | Head | 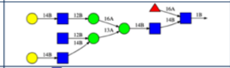 | 32/120 | 26.7% |
| N386 | Head | 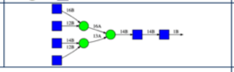 | 81/120 | 67.5% |
| TOTAL | | | 244/960 | 25.4% |

**Fig. S5.** Glycosylation of the 708 monomeric HAs (top) and 120 monomeric NAs (bottom) included in the H1N1-Shan2009 whole-virion model. The position of N-linked glycosylation sites on HA (top) and NA (bottom) monomeric structure is shown on the left panels. The structure, total number, and relative frequency (%) of glycans added at each N-linked site are summarized on the right panels.

**HA**

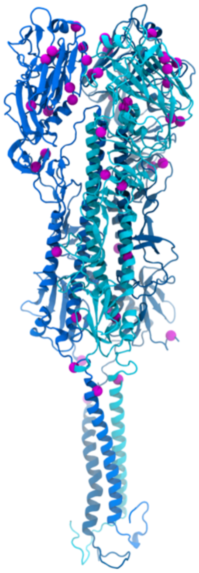

| # | RES NUMBER | H1N1-Mich2015 MUTATION |
|---|---|---|
| 1 | 13 | A13T |
| 2 | 101 | S101N |
| 3 | 114 | D114N |
| 4 | 179 | S179N |
| 5 | 180 | K180Q |
| 6 | 202 | S202T |
| 7 | 220 | S220T |
| 8 | 233 | I233T |
| 9 | 240 | Q240R |
| 10 | 273 | A273T |
| 11 | 300 | K300E |
| 12 | 302 | T302A |
| 13 | 391 | E391K |
| 14 | 468 | S468N |
| 15 | 516 | E516K |

**NA**

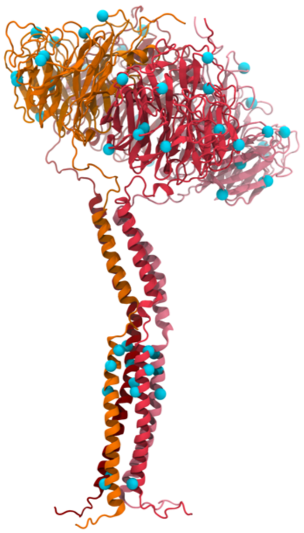

| # | RES NUMBER | H1N1-Mich2015 MUTATION |
|---|---|---|
| 1 | 13 | V13I |
| 2 | 34 | I34V |
| 3 | 39 | L39I |
| 4 | 43 | N43S |
| 5 | 200 | N200S |
| 6 | 241 | V241I |
| 7 | 248 | N248D |
| 8 | 264 | V264I |
| 9 | 269 | N269K |
| 10 | 314 | I314M |
| 11 | 321 | I321V |
| 12 | 368 | N368K |
| 13 | 386 | N386K |
| 14 | 432 | K432E |

**Fig. S6.** List of amino acid point mutations in H1N1-Mich2015 with respect to H1N-Shan2009 for HA (left panel) and NA (right panel). The addition of N-linked sequon within HA head at position N179 is highlighted in blue. Deletion of N-linked sequon within NA head at position N386 highlighted in red. The location of the point mutations within HA (left) and NA (right) structures is highlighted with magenta and cyan beads, respectively.

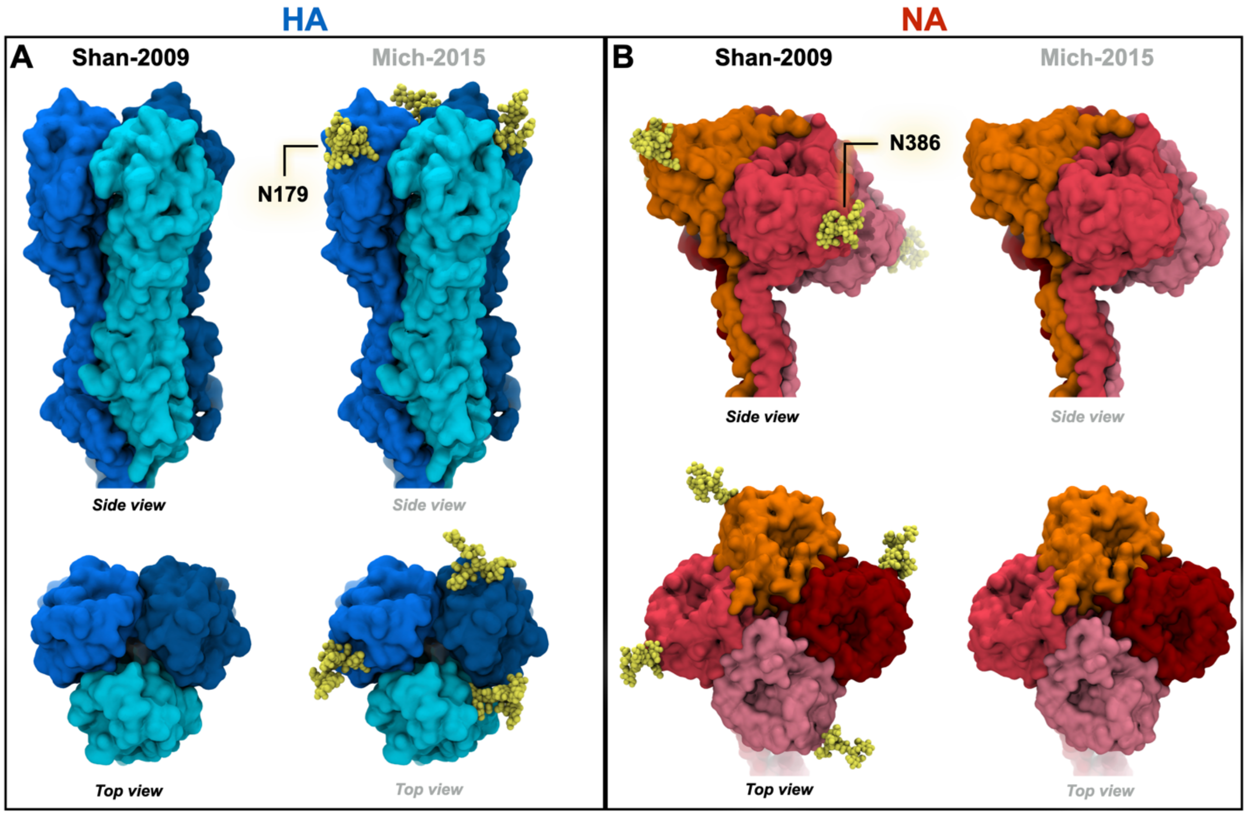


**Fig. S7.** Glycan variation in H1N1-Mich2015 with respect to H1N1-Shan2009 for HA (**A**) and NA (**B**). A new N-linked glycan is introduced at position N179 within the HA head (**A**). N-linked glycan at position N386 within the NA head is instead deleted (**B**). The glycan is illustrated with yellow van der Waals spheres, whereas HA and NA are represented with cyan and red surfaces, respectively.

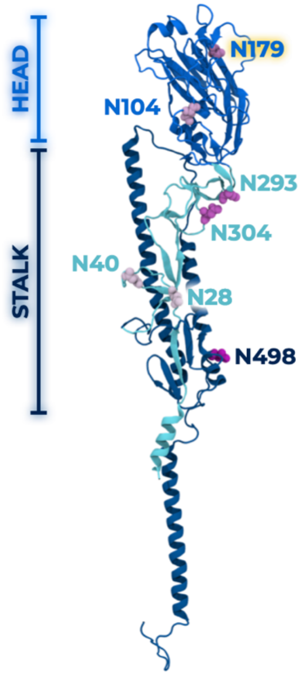


## HA

| Sequon | Region | Structure | # | % |
|---|---|---|---|---|
| N28 | Stalk | 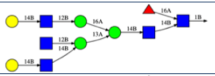 | 420/708 | 59.3 |
| N40 | Stalk | 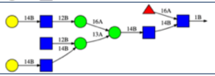 | 288/708 | 40.7% |
| N104 | Head | 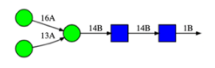 | 33/708 | 4.7% |
| N179 | Head | 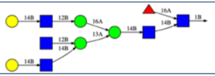 | 351/708 | 49.6% |
| N293 | Stalk | 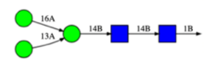 | 39/708 | 5.5% |
| N304 | Stalk | 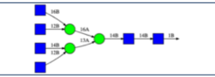 | 102/708 | 14.4% |
| N498 | Stalk | 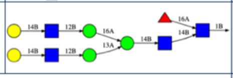 | 393/708 | 55.5% |
| TOTAL | | | 1626/4956 | 32.8% |

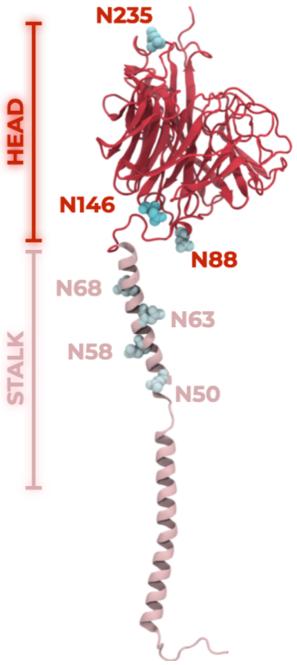


## NA

| Sequon | Region | Structure | # | % |
|---|---|---|---|---|
| N50 | Stalk | 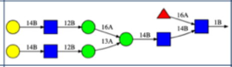 | 31/120 | 25.8% |
| N58 | Stalk | 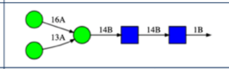 | 0/120 | 0% |
| N63 | Stalk | 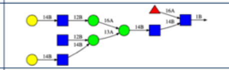 | 40/120 | 33.3% |
| N68 | Stalk | 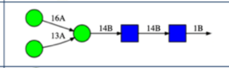 | 11/120 | 9.2% |
| N88 | Head | 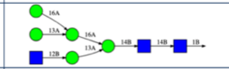 | 36/120 | 30.0% |
| N146 | Head | 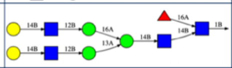 | 18/120 | 15.0% |
| N235 | Head | 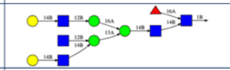 | 29/120 | 24.2% |
| ~~N386~~ | – | – | – | – |
| TOTAL | | | 165/840 | 19.6% |

**Fig. S8.** Glycosylation of the 708 monomeric HAs (top) and 120 monomeric NAs (bottom) included in the H1N1-Mich2015 whole-virion model. The position of N-linked glycosylation sites on the HA (top) and NA (bottom) monomeric structure is shown on the left panels. The structure, total number, and relative frequency (%) of glycans added at each N-linked site are summarized on the right panels.

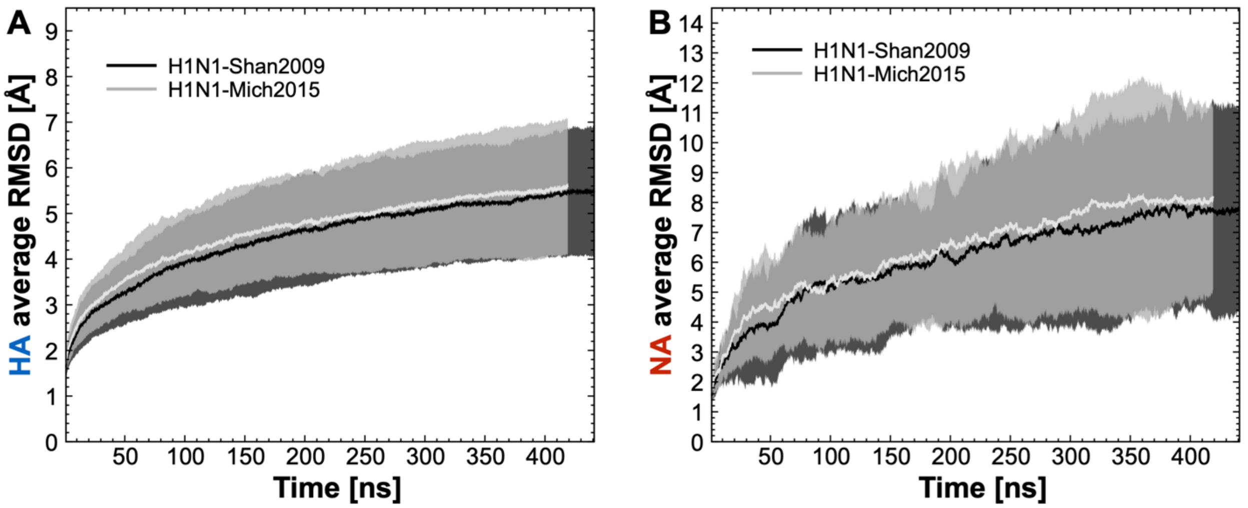


**Fig. S9.** Average RMSD profiles for HA (**A**) and NA (**B**) as calculated over the course of H1N1-Shan2009 (black line) and H1N1-Mich2015 (gray line) whole-virion MD simulations. RMSD values have been averaged across the 236 HA trimers and 30 NA tetramers, respectively. Standard deviation values are plotted with a semi-transparent filled area.

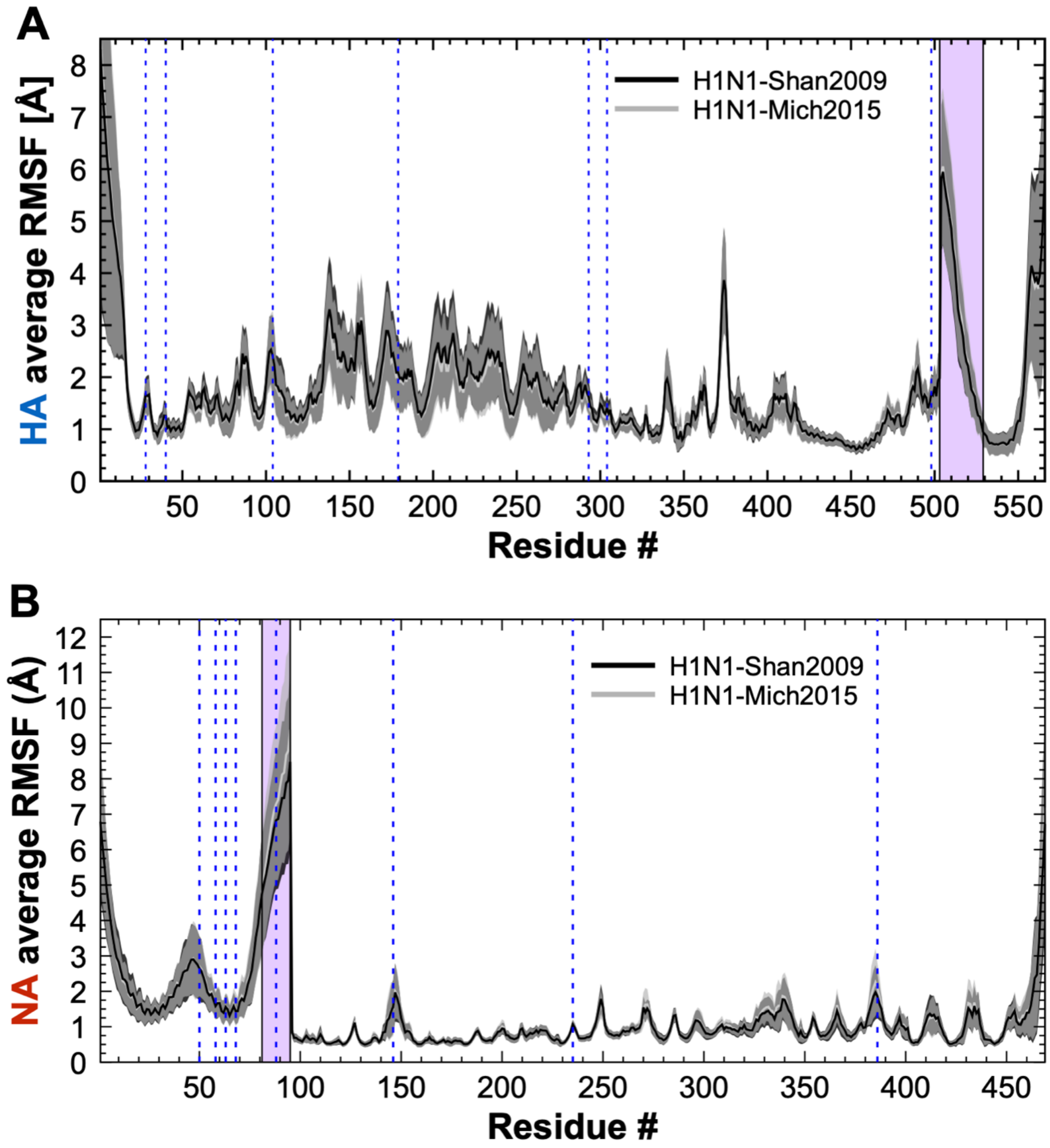


**Fig. S10.** Average RMSF profiles for HA (**A**) and NA (**B**) as calculated over the course of H1N1-Shan2009 (black line) and H1N1-Mich2015 (gray line) whole-virion MD simulations. RMSF values have been averaged across the 708 HA and 120 NA monomers, respectively. Standard deviation values are plotted with a semi-transparent filled area. Highlighted with a transparent purple rectangle is the flexible hinge region for HA and NA facilitating HA ectodomain tilting and NA head tilting, respectively. The left side of the purple rectangle in the HA RMSF profile demarcates the two alignment HA regions used for RMSF calculation. The left side of the purple rectangle in the HA RMSF profile demarcates the two alignment regions used for RMSF calculation.

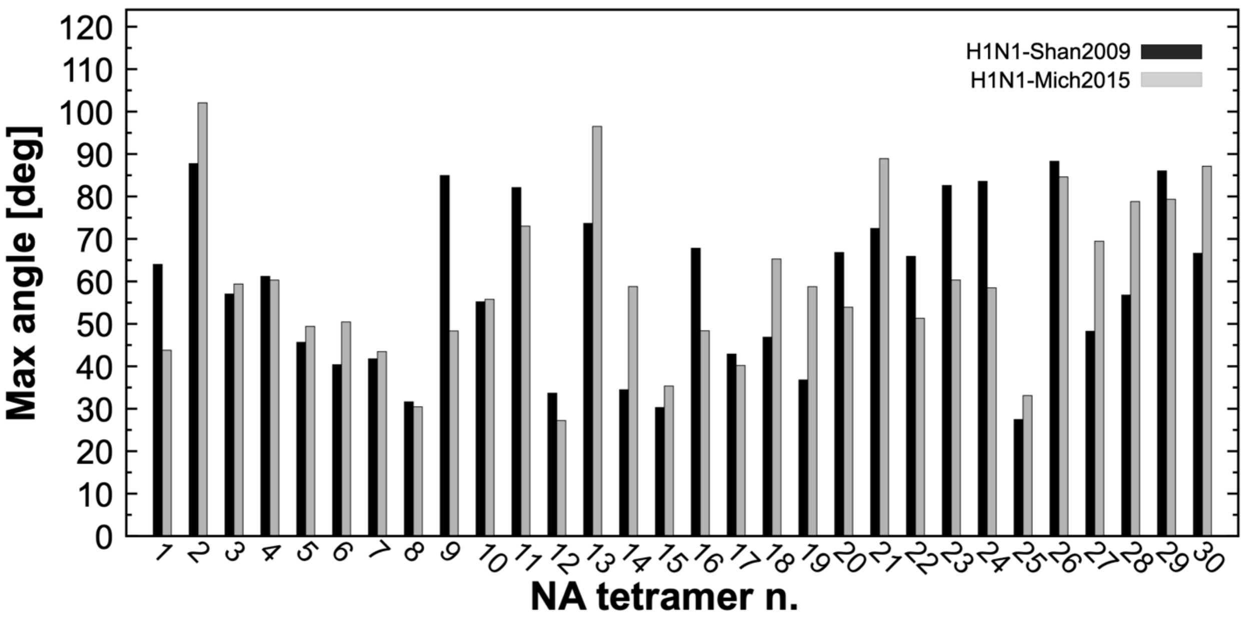


**Fig. S11.** Maximum NA head-tilt angle observed in H1N1-Shan2009 and H1N1-Mich2015 NA tetramers. For each NA tetramer (numbered from #1 to #30), the maximum NA head-tilt angle values (degree) calculated during the simulations of H1N1-Shan2009 and H1N1-Mich205 are shown with black and gray bars, respectively.

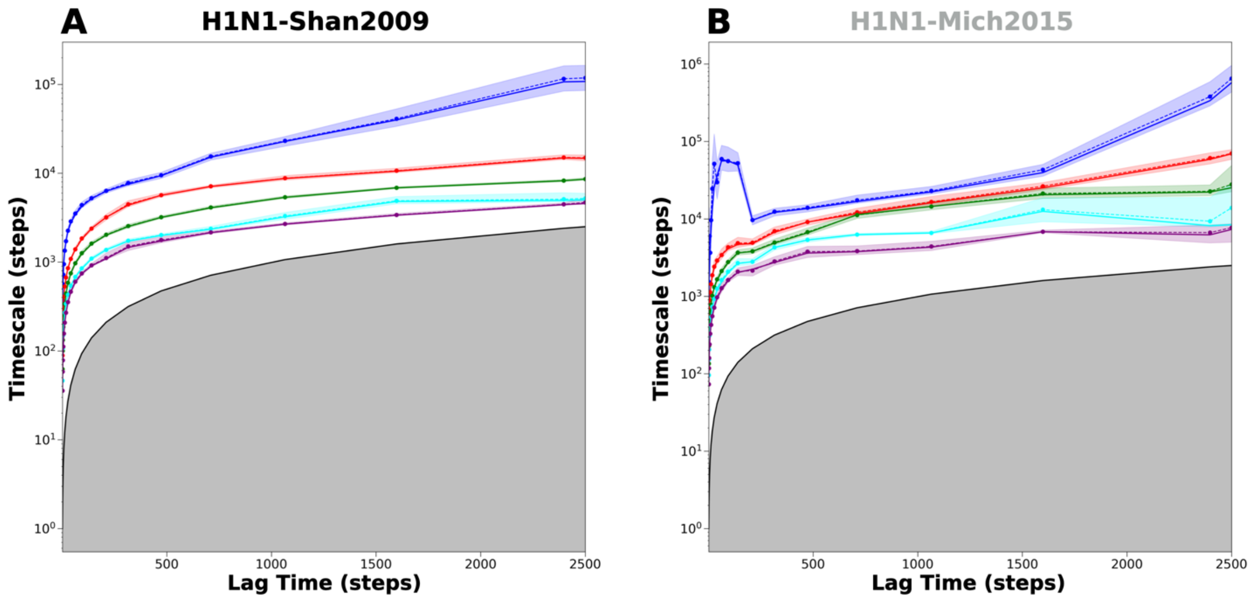


**Fig. S12.** The implied timescale plots for the NA head-tilt MSM. The shading on each of the implied timescales represents statistical uncertainties calculated through Bayesian sampling. From these figures, the lag time was selected to be 250 steps and two macrostates were selected. The gray region represents forbidden space, which is faster than the lag time; transitions in this region cannot be resolved.

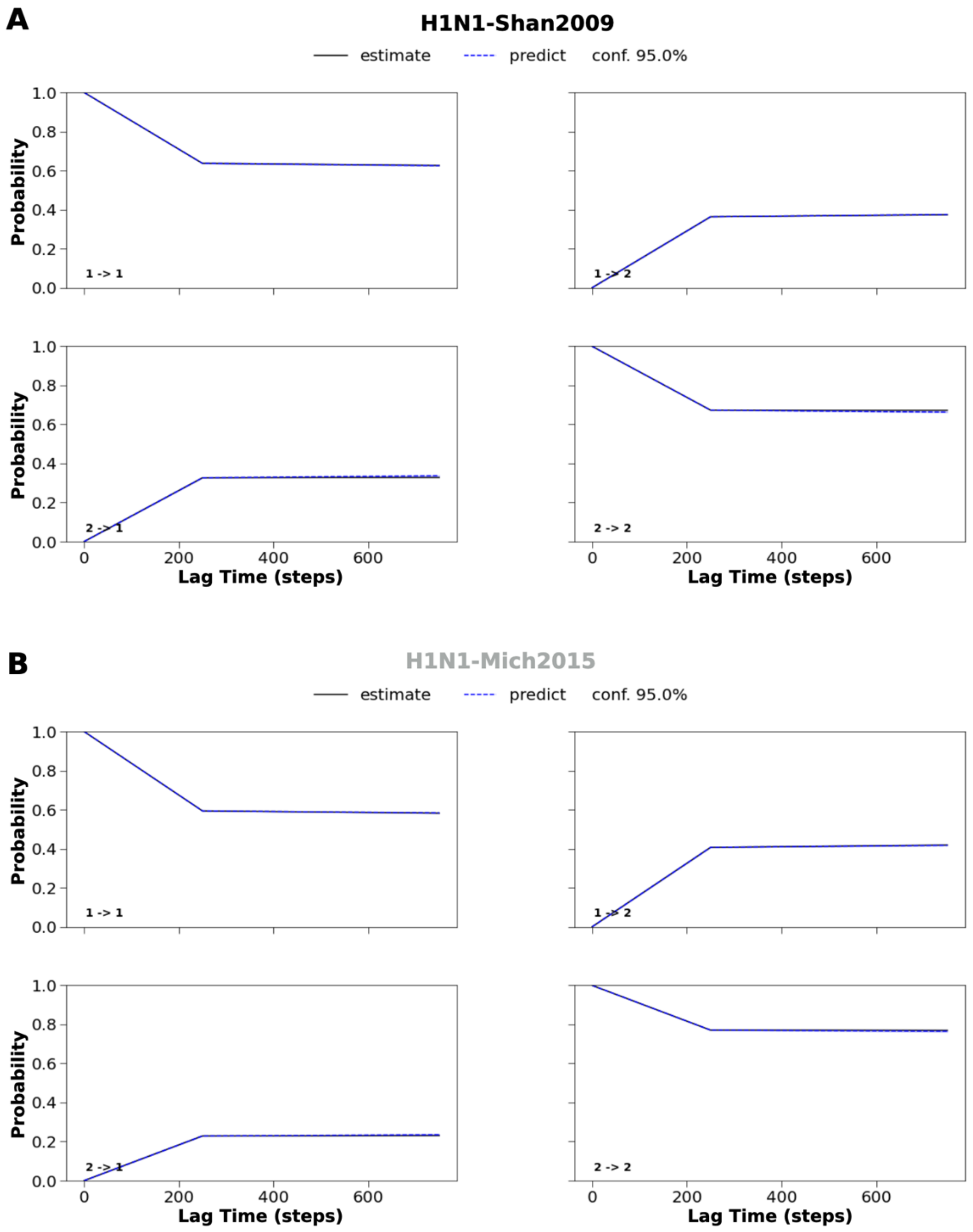


**Fig. S13.** The CK test plots for the NA head-tilt MSM of (**A**) H1N1-Shan2009 and (**B**) H1N1-Mich2015. The lag time was selected to be 250 steps, with two macrostates selected. The solid line (the "estimate") represents the transition probabilities of the model created at a lag time of 250 steps, with these probabilities then estimated at multiples of the lag time. The dotted line (the "prediction") represents predicted probabilities of the same model independently created at each of these multiples of the lag time. Dark blue shading represents a 95% confidence interval for the predicted probabilities. Since the estimates of the MSM model that we used overlap with the predicted probabilities of the model at 3x the lag time, this model satisfies Markovianity, passes the CK test and can be used to extract transition kinetics. The numbers in the lower left corners represent macrostate transitions; thus "1 -> 1" represents a transition from macrostate 1 to macrostate 1.

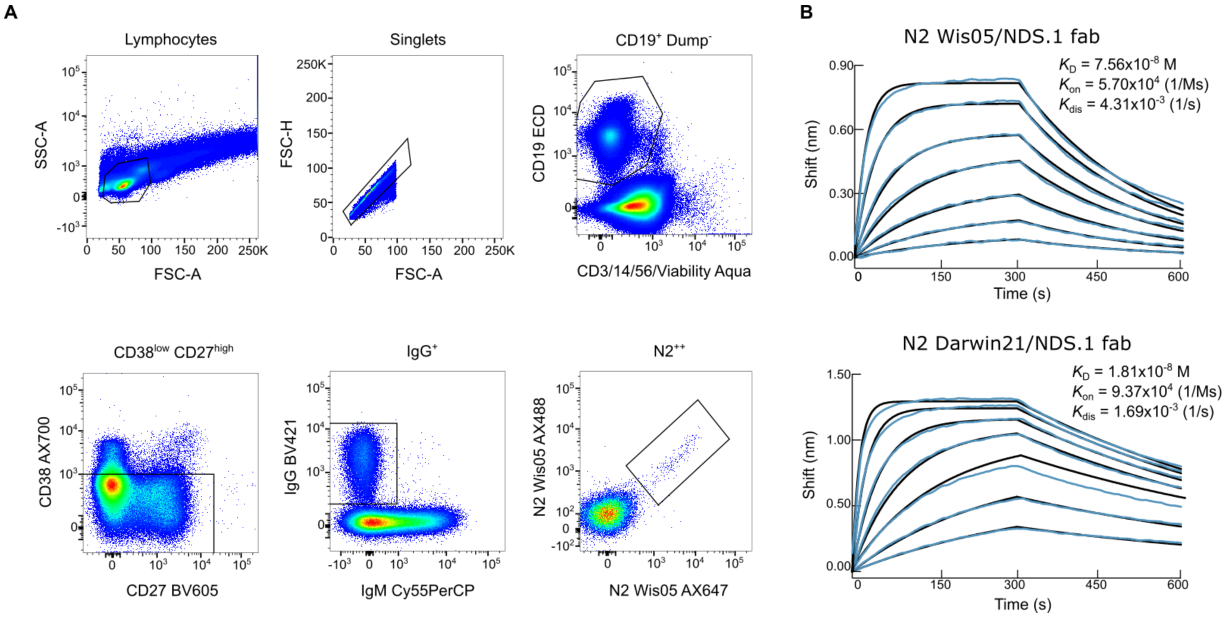


**Fig. S14.** Isolation and characterization of NA-directed mAb NDS.1. (**A**) Gating strategy of NA-specific memory B-cells by flow cytometry. (**B**) Binding kinetics of NDS.1 Fab to N2-Wis05 and N2-Darwin21 by biolayer interferometry. Experimental data (blue traces) were fitted (black lines) with the binding equations describing a 1:1 interaction.

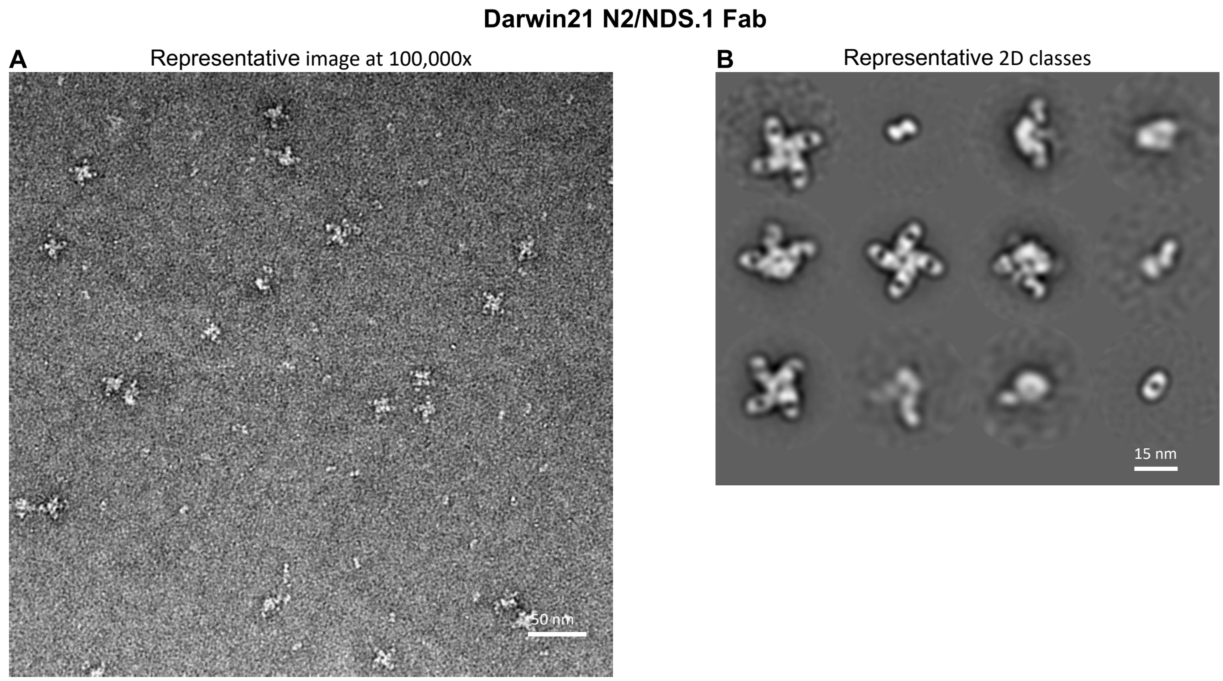


**Fig. S15.** Negative-stain EM analysis of N2 A/Darwin/9/2021 NA tetramer in complex with Fab NDS.1 (**A)** Representative raw NS-EM image and (**B**) 2D class averages of N2 A/Darwin/9/2021 NA in complex with Fab NDS.1 (scale bar: 50 nm and 15 nm).

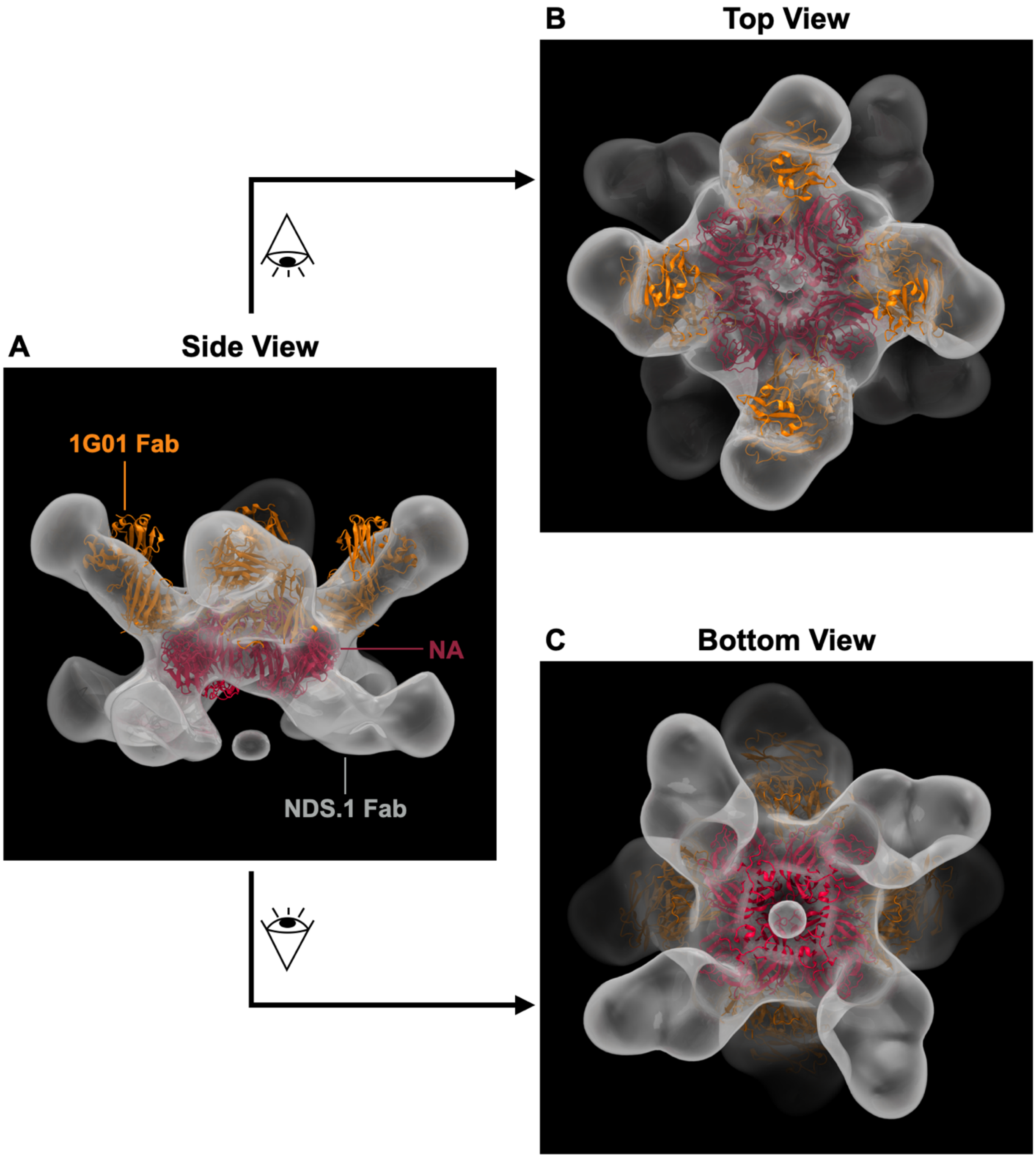


**Fig. S16.** NS-EM reconstruction model of N2 A/Indiana/10/2011 NA tetramer in complex with NDS.1 Fab and 1G01 Fab. Side view (**A**), top view (**B**) and bottom view (**C**) of the ternary complex. Crystal structure of NA-1G01 (PDB: 6Q23 (*67*)) is fitted into the NS-EM density map. The final model was generated using a dataset containing 4,368 particles.

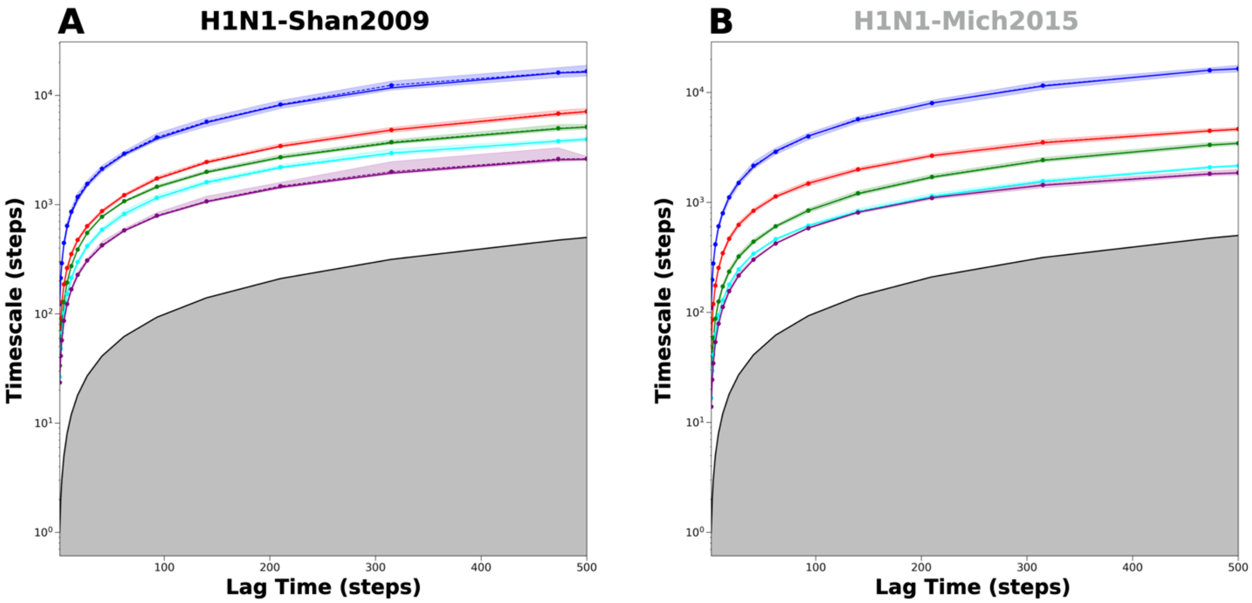


**Fig. S17.** The implied timescale plots for the HA ectodomain-tilt MSM of (**A**) H1N1-Shan2009 and (**B**) H1N1-Mich2015. The shading on each of the implied timescales represents statistical uncertainties calculated through Bayesian sampling. From these figures, the lag time was selected to be 50 steps and two macrostates were selected. The gray region represents forbidden space, which is faster than the lag time; transitions in this region cannot be resolved.

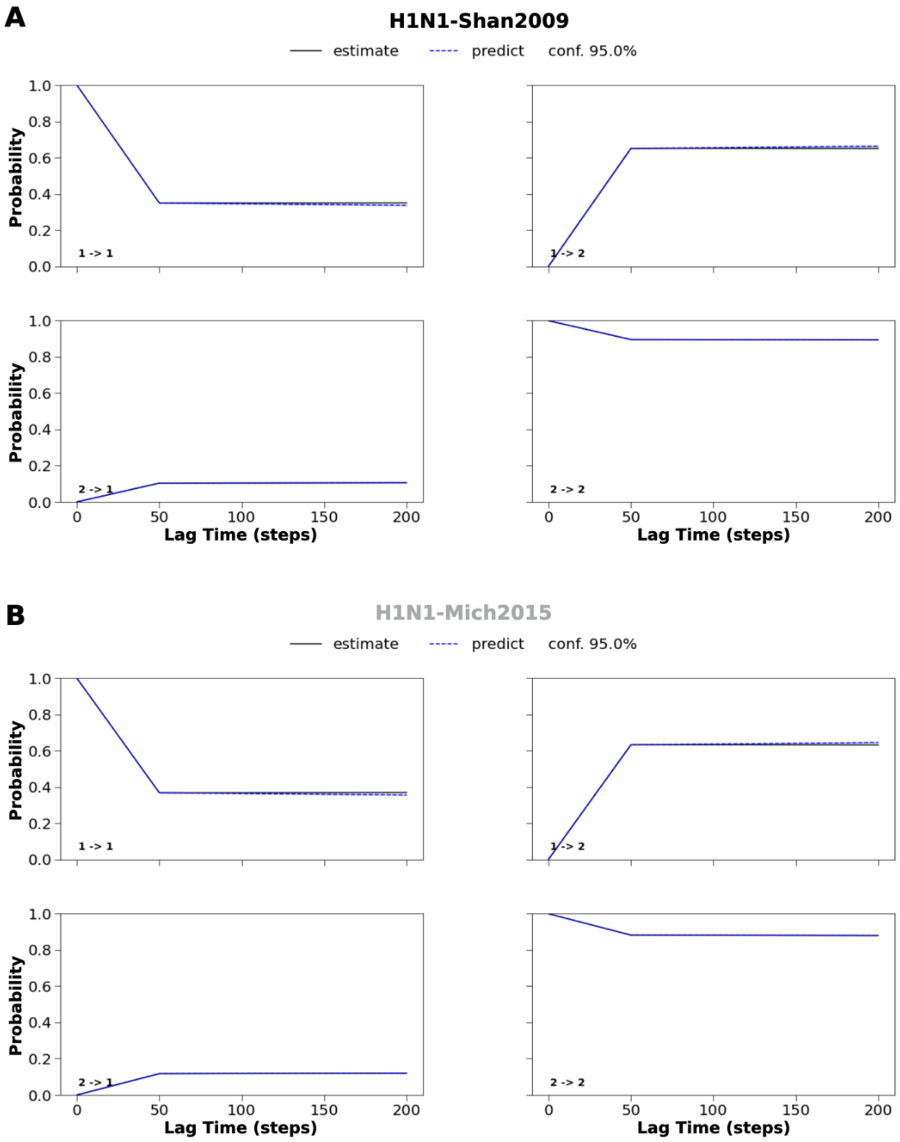


**Fig. S18.** The CK test plots for the HA ectodomain-tilt MSM of (**A**) H1N1-Shan2009 and (**B**) H1N1-Mich2015. The lag time was selected to be 50 steps, with two macrostates selected. The solid line (the "estimate") represents the transition probabilities of the model created at a lag time of 50 steps, with these probabilities then estimated at multiples of the lag time. The dotted line (the "prediction") represents predicted probabilities of the same model independently created at each of these multiples of the lag time. Dark blue shading represents a 95% confidence interval for the predicted probabilities. Since the estimates of the MSM model we used overlap with the predicted probabilities of the model at 3x the lag time, this model satisfies Markovianity, passes the CK test and can be used to extract transition kinetics. The numbers in the lower left corners represent macrostate transitions; thus "1 -> 1" represents a transition from macrostate 1 to macrostate 1.

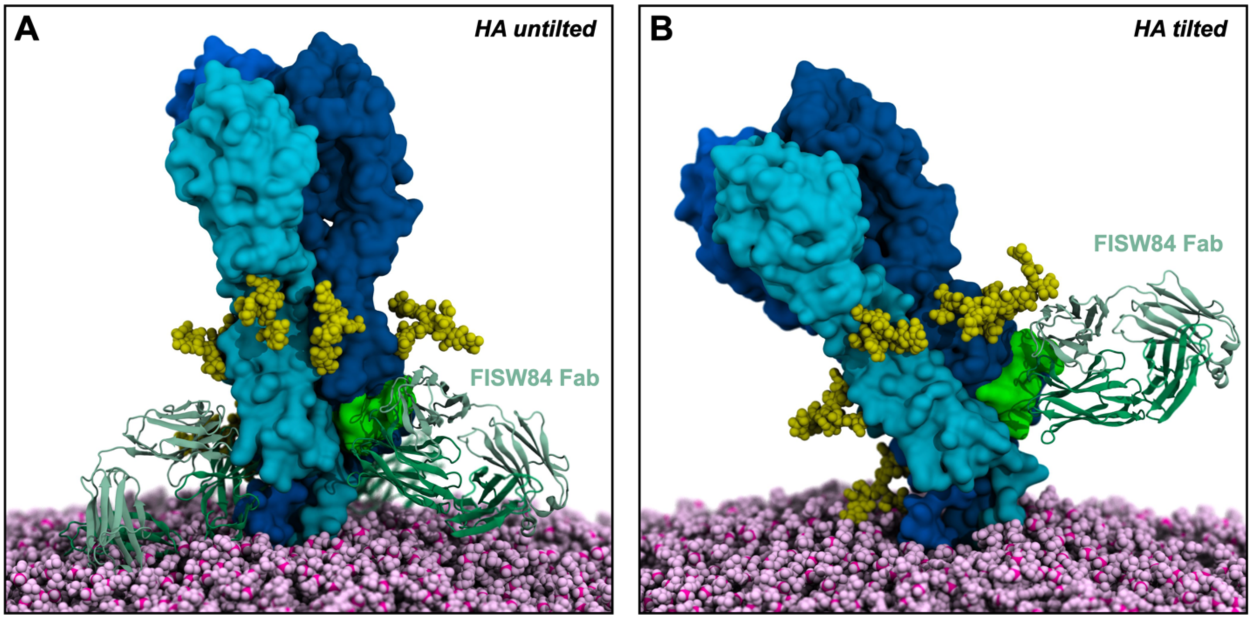


**Fig. S19.** HA anchor epitope accessibility. **A**) When HA is untilted, the FISW84 Fab (*37*) must approach the anchor epitope with an upward angle. FISW84 is depicted with green cartoons, whereas the three HA monomers are depicted with surfaces colored with different shades of blue. The anchor epitope is highlighted with a green surface. N-glycans are shown with yellow vdW spheres, whereas the POPC lipids are displayed with pink vdW spheres. **B**) When HA flexes onto the membrane, the angle of approach to at least one epitope gradually shifts from upward to lateral or slightly downward, facilitating FISW84 Fab binding.

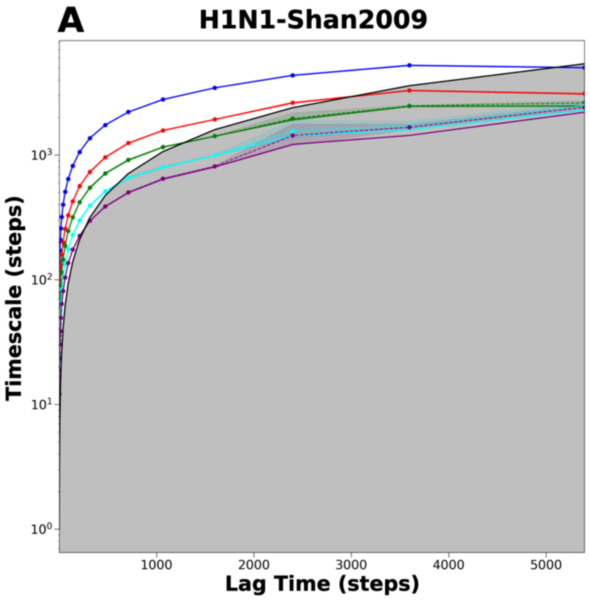


**Fig. S20.** The implied timescale plots for the HA head breathing MSM of (**A**) H1N1-Shan2009 and (**B**) H1N1-Mich2015. The shading on each of the implied timescales represents statistical uncertainties calculated through Bayesian sampling. From these figures, the lag time was selected to be 500 steps and two macrostates were selected. The gray region represents forbidden space, which is faster than the lag time; transitions in this region cannot be resolved.

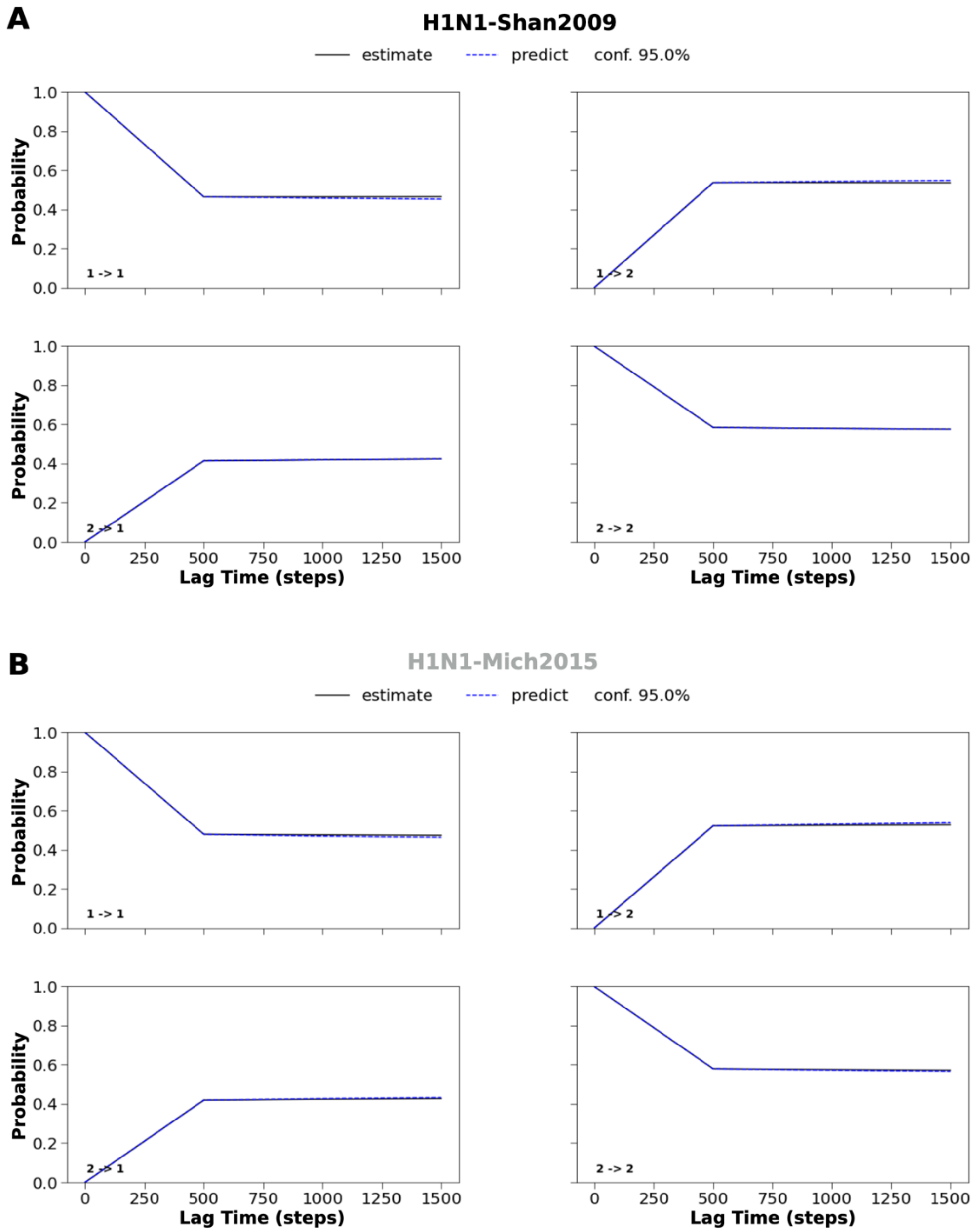


**Fig. S21.** The CK test plots for the HA head breathing MSM of (**A**) H1N1-Shan2009 and (**B**) H1N1-Mich2015. The lag time was selected to be 500 steps, with two macrostates selected. The solid line (the "estimate") represents the transition probabilities of the model created at a lag time of 500 steps, with these probabilities then estimated at multiples of the lag time. The dotted line (the "prediction") represents predicted probabilities of the same model independently created at each of these multiples of the lag time. Dark blue shading represents a 95% confidence interval for the predicted probabilities. Since the estimates of the MSM model we used overlap with the predicted probabilities of the model at 3x the lag time, this model satisfies Markovianity, passes the CK test and can be used to extract transition kinetics. The numbers in the lower left corners represent macrostate transitions; thus "1 -> 1" represents a transition from macrostate 1 to macrostate 1.

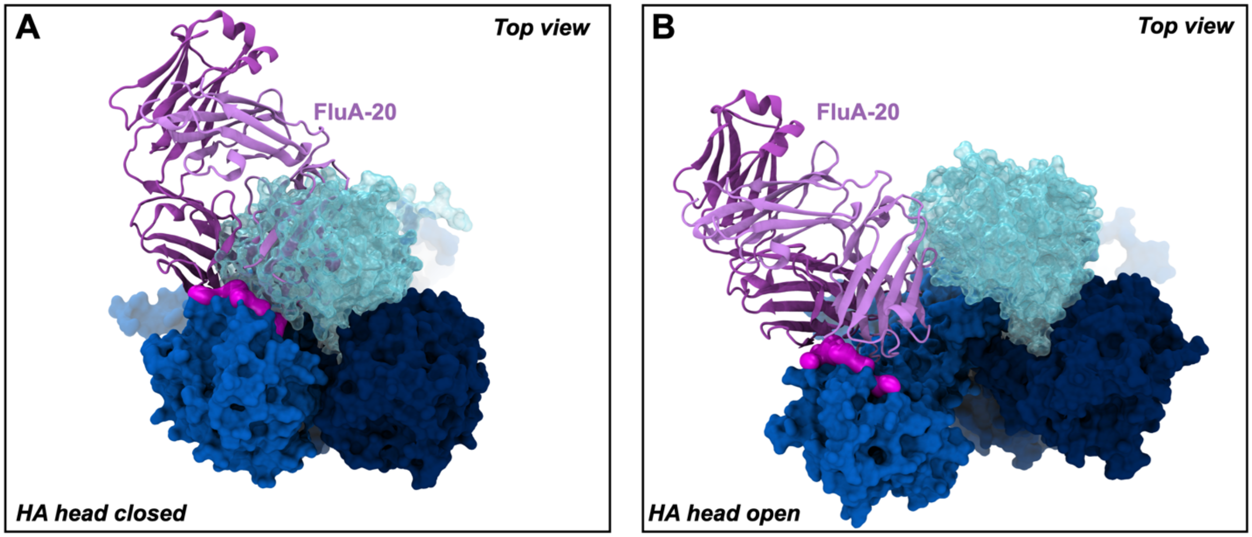


**Fig. S22.** HA head trimer interface epitope accessibility. **A**) When HA heads are closed, the FluA-20 antibody (*39*) cannot access the epitope buried in the head trimer interface. FluA-20 is depicted with purple cartoons, whereas the three HA monomers are depicted with surfaces colored with different shades of blue. The FluA-20 epitope is highlighted with a magenta surface on one HA head. **B**) When one HA breathes, the FluA-20 epitope becomes accessible allowing antibody binding.

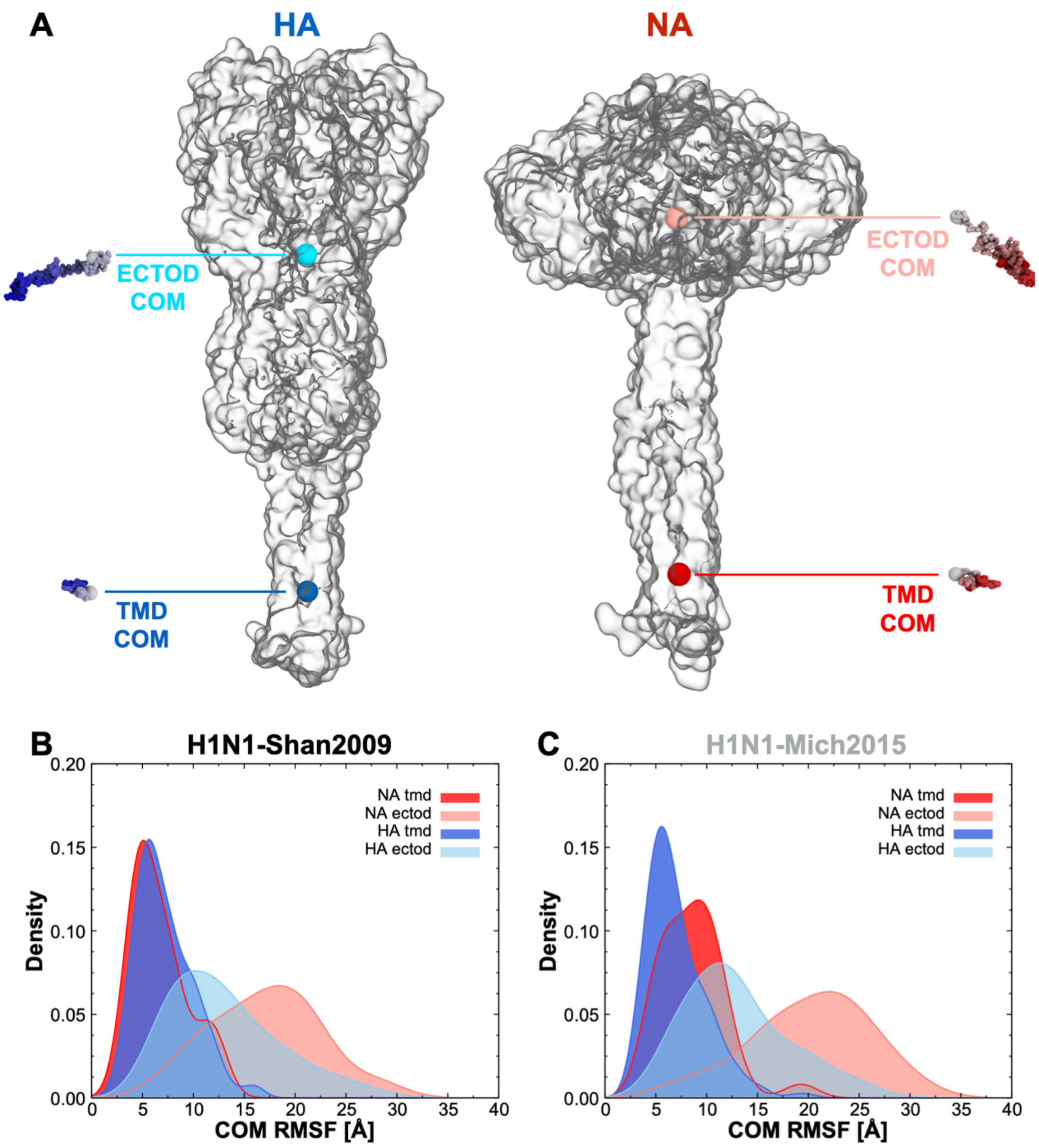


**Fig. S23.** RMSF of HA and NA transmembrane domain (TMD) COM and ectodomain COM. The initial frame was used as a reference for RMSF calculation. **A**) Molecular representation of an HA trimer (left) and a NA tetramer (right) using a transparent surface. TMD COM and ectodomain COM of HA and NA are indicated with cyan and red spheres, respectively. As an example, the position of TMD COM and ectodomain COM is displayed with a sphere every 10$^{th}$ frame of the MD simulation for a representative HA trimer and a representative NA tetramer selected from H1N1-Shan2009. The spheres are colored according to simulation time using a white-to-blue gradient for HA and a white-to-red gradient for NA. **B**) Kernel density distribution profiles of RMSF values calculated for the HA (236 trimers) and NA (30 tetramers) TMD COMs and ectodomain COMs in H1N1-Shan2009. **C**) Kernel density distribution profiles of RMSF values calculated for the HA (236 trimers) and NA (30 tetramers) TMD COMs and ectodomain COMs in H1N1-Mich2015.

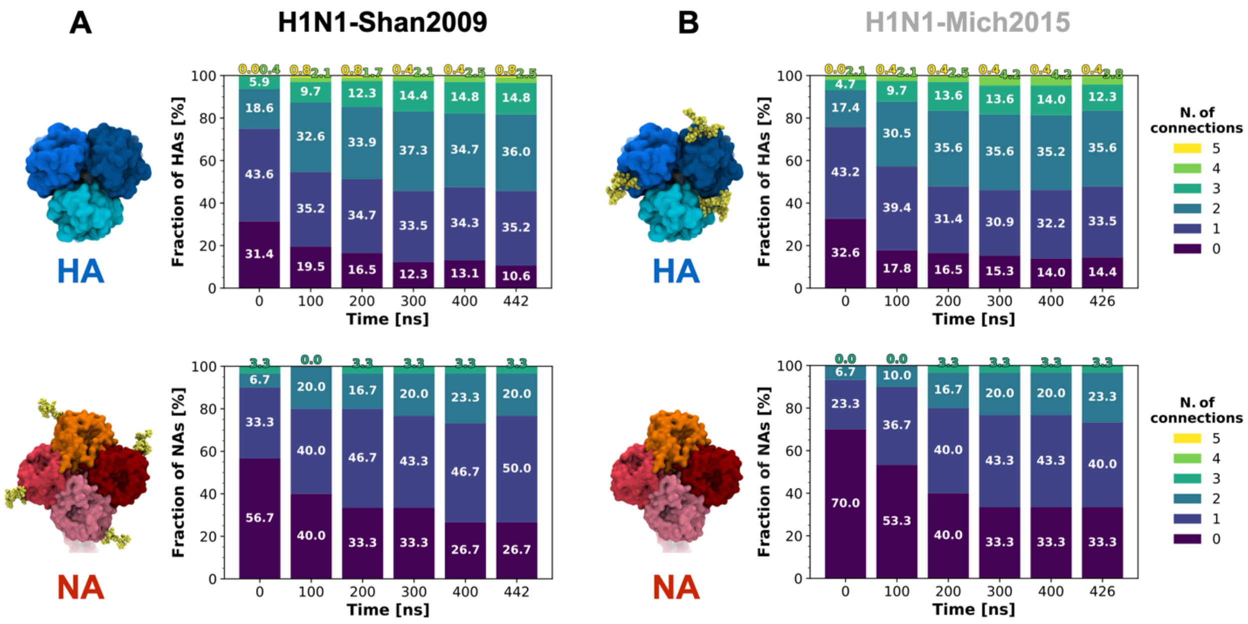


**Fig. S24.** Analysis of glycoprotein inter-connections for HA and NA. Stacked bar plots representing the fraction (%) of glycoproteins (HA top panels, NA bottom panels) establishing a certain number of connections (0 to 5, accordingly colored using the viridis palette) with surrounding glycoproteins along the simulation of (**A**) H1N1-Shan2009 and (**B**) H1N1-Mich2015. Percentages rounded to the first decimal digit are reported. A molecular representation of the HA head and NA head is also provided for both strains on the left side of each bar plot, highlighting with yellow vdW spheres the glycan that is added/deleted in H1N1-Mich2015: +1 glycan (N179) in H1N1-Mich2015 HA head and -1 glycan (N386) in H1N1-Mich2015 NA head.

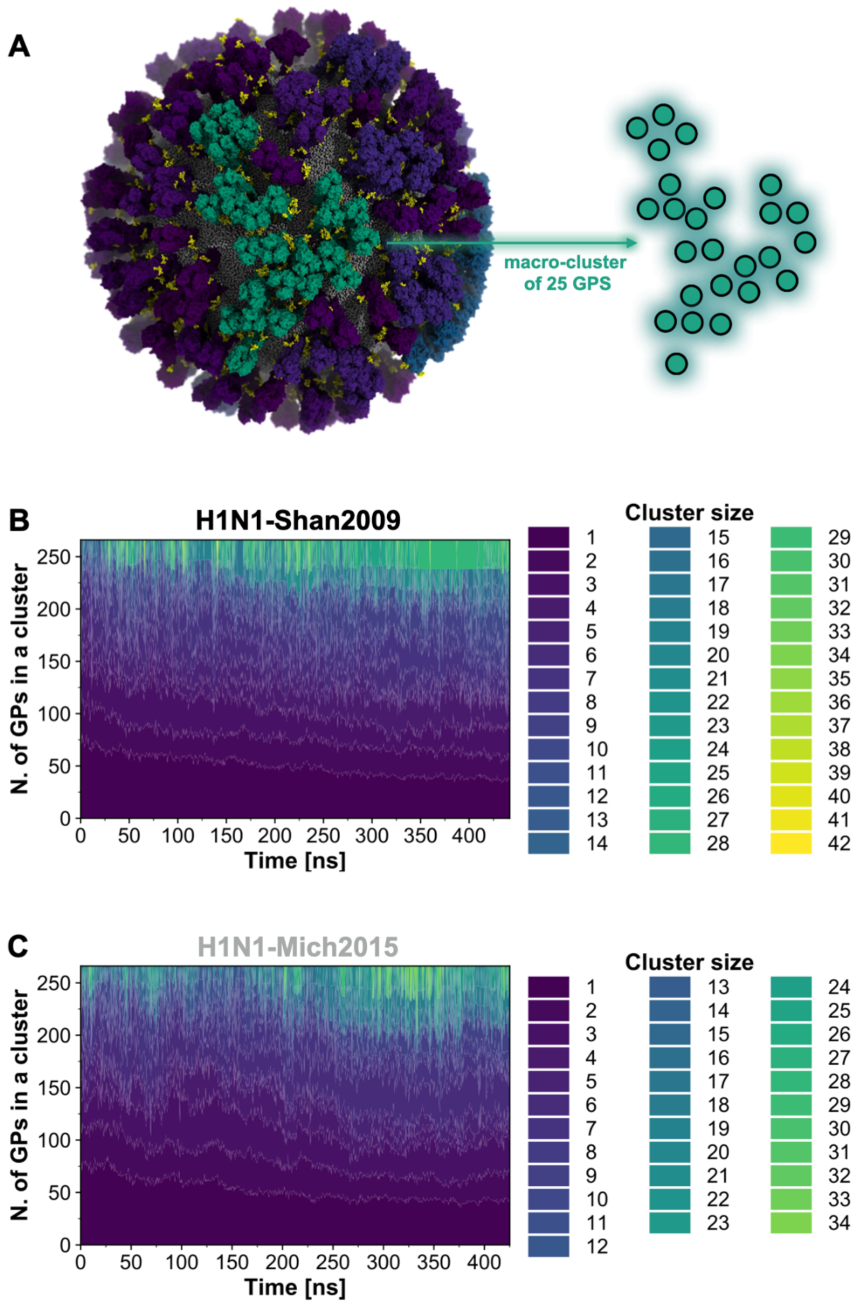


**Fig. S25.** Analysis of glycoprotein macro-cluster formation. **A**) Molecular representation of the H1N1-Shan2009 virion at a frame of the simulation where a macro-cluster of 25 glycoproteins (GPs) is formed. The glycoproteins are depicted with a surface colored using a viridis palette according to the size of the macro-cluster that they belong to. The pattern of interactions of the macro-cluster of 25 GPs is highlighted using colored circles in the schematic on the right. **B-C**) Stacked plots representing the number of GPs involved in a macro-cluster of a certain size along the simulation of (**B**) H1N1-Shan2009 and (**C**) H1N1-Mich2015. Macro-clusters are colored using the *viridis* palette according to their size. The color gradient used for H1N1-Mich2015 (cluster size range: 0 to 34) has been normalized to the color gradient of H1N1-Shan2009 (cluster size range: 0 to 42).

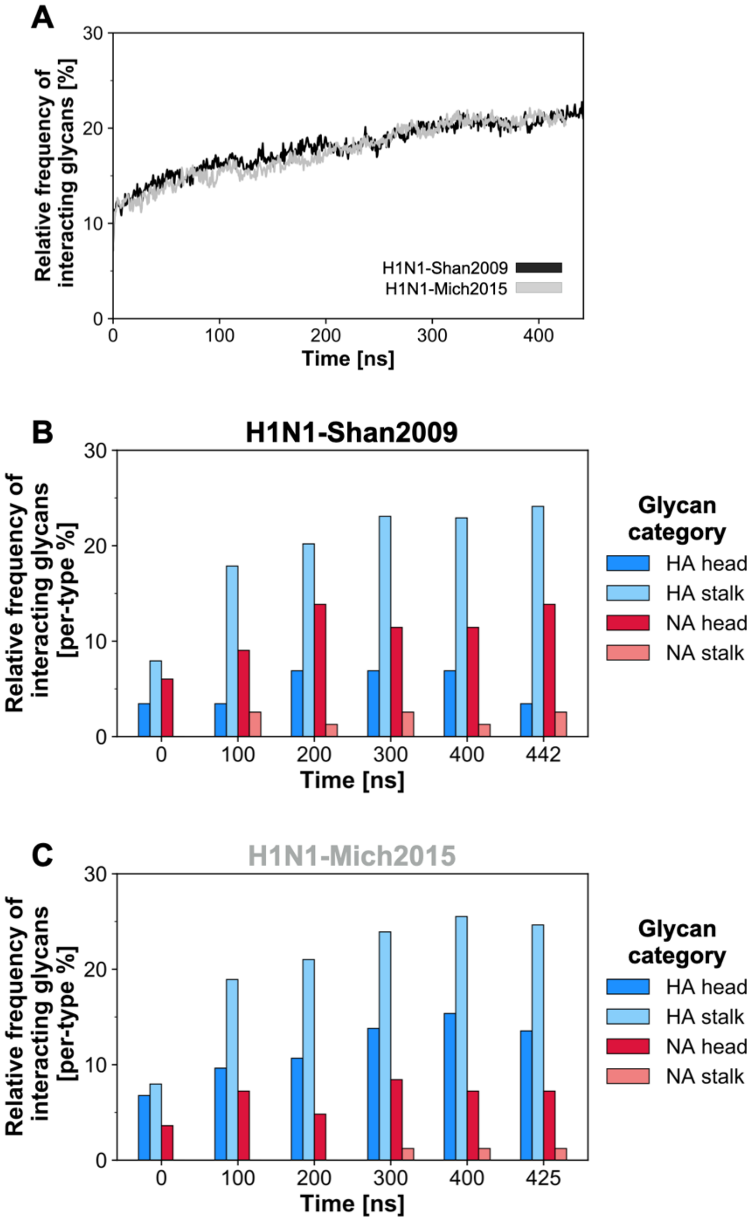


**Fig. S26.** Analysis of glycoprotein connections involving glycans. **A**) Time evolution (ns) of the relative frequency (%) of interacting glycans, i.e., glycans participating in glycoprotein interconnections. The relative frequency of interacting glycan was calculated with respect to the total number of glycans. **B**) Grouped bar plot depicting the time evolution (ns) of the relative frequency (%) of interacting glycans in H1N1-Shan2009, where glycans have been divided into "head" and "stalk" categories according to their position, both for HA (shades of blue) and NA (shades of red). The relative frequency for each category of interacting glycans was calculated with respect to the total number of glycans of the same category. **C**) Grouped bar plot depicting the time evolution (ns) of the relative frequency (%) of interacting glycans in H1N1-Mich2015, where glycans have been divided into "head" and "stalk" categories according to their position, both for HA (shades of blue) and NA (shades of red). The relative frequency for each category of interacting glycans was calculated with respect to the total number of glycans of the same category.

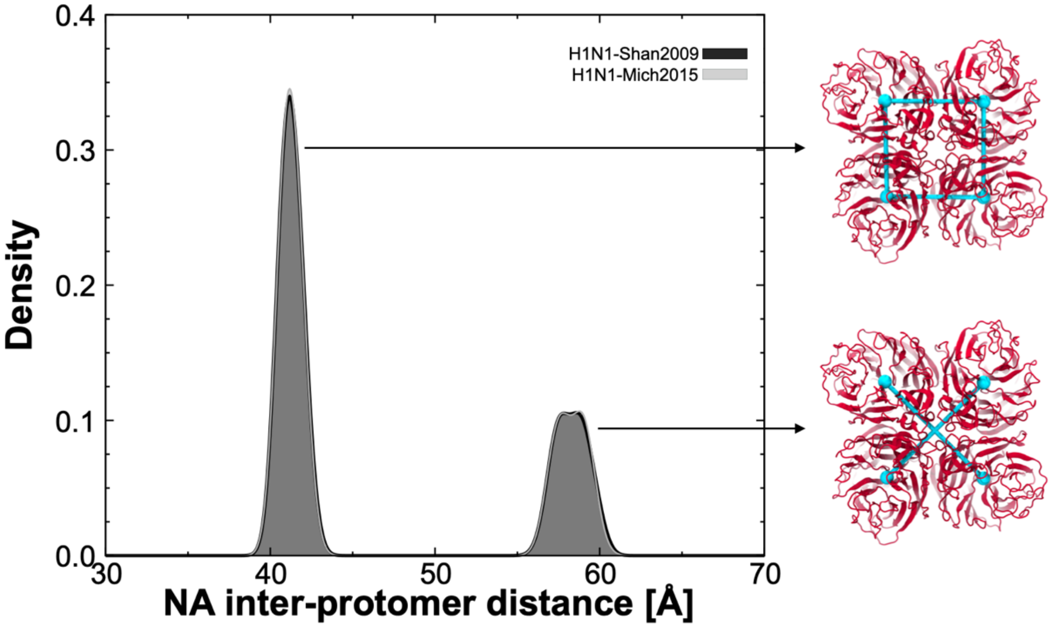


**Fig. S27.** Analysis of NA head breathing. Distribution of inter-protomer distances between the four NA head COMs in H1N1-Shan2009 (black) and H1N1-Mich2015 (gray) virions. The distribution is shown as a kernel density. On the right, a top view molecular representation of the NA head using red cartoons is provided, with the monitored distances between NA head COMs highlighted with cyan cylinders.

**Table S1.**

Tilt angles (degree) of ten tilted and untilted structures (microstates) extracted from the MSM of the NA head tilt. Note: the average is the average of the ten selected microstates (ten selected conformational states and their respective angles), not all microstates assigned to these respective macrostates.

| **NA HEAD-TILT ANGLES [°]** | | | | |
|---|---|---|---|---|
| | **H1N1-Shan2009** | | **H1N1-Mich2015** | |
| **Structure #** | **Untilted** | **Tilted** | **Untilted** | **Tilted** |
| **#1** | 25.1° | 79.6° | 6.7° | 91.7° |
| **#2** | 18.5° | 81.6° | 18.1° | 91.7° |
| **#3** | 28.7° | 78.4° | 19.7° | 89.6° |
| **#4** | 24.2° | 75.0° | 20.6° | 91.0° |
| **#5** | 24.7° | 80.5° | 18.8° | 93.4° |
| **#6** | 25.5° | 75.2° | 18.3° | 90.3° |
| **#7** | 28.5° | 81.7° | 20.7° | 93.5° |
| **#8** | 28.4° | 75.8° | 14.3° | 91.3° |
| **#9** | 22.0° | 80.5° | 15.8° | 92.7° |
| **#10** | 24.9° | 75.0° | 18.3° | 94.2° |
| **average** | **25.0°** | **78.3°** | **17.1°** | **91.9°** |

**Table S2.**

Tilt angles (degree) of ten tilted and untilted structures (microstates) extracted from the MSM of the HA ectodomain tilt. Note: the average is the average of the ten selected microstates (ten selected conformational states and their respective angles), not all microstates assigned to these respective macrostates.

| **HA ECTODOMAIN-TILT ANGLES [°]** | | | | |
|---|---|---|---|---|
| | **H1N1-Shan2009** | | **H1N1-Mich2015** | |
| **Structure #** | **Untilted** | **Tilted** | **Untilted** | **Tilted** |
| **#1** | 14.5° | 57.9° | 10.3° | 63.1° |
| **#2** | 7.9° | 60.8° | 18.7° | 64.4° |
| **#3** | 9.5° | 56.2° | 16.6° | 65.8° |
| **#4** | 10.0° | 56.7° | 15.6° | 65.2° |
| **#5** | 9.7° | 60.1° | 17.6° | 63.7° |
| **#6** | 2.5° | 57.2° | 8.1° | 62.6° |
| **#7** | 9.7° | 57.9° | 4.5° | 60.9° |
| **#8** | 9.8° | 58.6° | 14.7° | 62.3° |
| **#9** | 10.8° | 57.4° | 17.8° | 66.3° |
| **#10** | 3.1° | 59.0° | 20.1° | 64.2° |
| **average** | **8.8°** | **58.2°** | **14.4°** | **63.9°** |

**Table S3.**

The table reports the distance between the COM of the HA monomer head (residues 66 to 286) and the COM of LαHs' apical residue (419) calculated for open and closed structures (microstates) extracted from the MSM of the HA head breathing. The distances indicated here are the largest seen in each microstate, i.e., the largest of the three distances existing within the conformation of the HA trimer represented by the microstate. This applies also to the "closed" microstates, where the reported distance is still the largest distance seen in those microstates. Note: the average is the average of the ten selected microstates (ten selected conformational states and their respective angles), not all microstates assigned to these respective macrostates.

| **HA HEAD COM–LαHs' apical residue COM DISTANCE [Å]** | | | | |
|---|---|---|---|---|
| | **H1N1-Shan2009** | | **H1N1-Mich2015** | |
| **Structure #** | **Closed** | **Open** | **Closed** | **Open** |
| **#1** | 27.8 Å | 38.5 Å | 27.2 Å | 35.2 Å |
| **#2** | 28.2 Å | 37.1 Å | 27.1 Å | 36.1 Å |
| **#3** | 27.8 Å | 39.0 Å | 27.2 Å | 37.6 Å |
| **#4** | 27.2 Å | 38.1 Å | 27.4 Å | 35.8 Å |
| **#5** | 27.4 Å | 38.5 Å | 27.9 Å | 35.9 Å |
| **#6** | 27.4 Å | 38.5 Å | 27.0 Å | 37.3 Å |
| **#7** | 27.2 Å | 39.0 Å | 27.2 Å | 36.9 Å |
| **#8** | 27.6 Å | 38.5 Å | 27.2 Å | 36.4 Å |
| **#9** | 27.0 Å | 37.1 Å | 27.0 Å | 36.0 Å |
| **#10** | 27.4 Å | 38.5 Å | 27.0 Å | 35.5 Å |
| **average** | **27.5 Å** | **38.3 Å** | **27.2 Å** | **36.3 Å** |

**Movie S1.**

All-atom MD simulation of the glycosylated model of H1N1-Shan2009 influenza A virion. ~442 ns of MD are shown. Neuraminidase (NA) and hemagglutinin (HA) glycoproteins, and the M2 ion channels, are depicted with red, blue, and white surfaces, respectively. N-linked glycans are shown in yellow. The spherical lipid envelope is represented with pink van der Waals (vdW) spheres.

**Movie S2.**

All-atom MD simulation of the glycosylated model of H1N1-Mich2015 influenza A virion. ~425 ns of MD are shown. Neuraminidase (NA) and hemagglutinin (HA) glycoproteins, and the M2 ion channels, are depicted with red, blue, and white surfaces, respectively. N-linked glycans are shown in yellow. The spherical lipid envelope is represented with pink van der Waals (vdW) spheres.

**Movie S3.**

NA head-tilt motion. The head-tilt motion of a representative NA tetramer extracted from the H1N1-Shan2009 whole-virion simulation is displayed. NA tetramer is depicted with a transparent surface overlaid on top of a cartoon representation colored with a viridis palette according to the extent of head tilt. N-linked glycans and sequon asparagine are shown with yellow van der Waals (vdW) spheres.

**Movie S4.**

NA head-tilt motion. The head-tilt motion of a NA tetramer is displayed together with the dynamics of the surrounding crowded environment of the H1N1-Shan2009 whole-virion. ~442 ns of MD are shown. Neuraminidase (NA) and hemagglutinin (HA) glycoproteins, and the M2 ion channels, are depicted with red, blue, and white surfaces, respectively. N-linked glycans are shown in yellow. The spherical lipid envelope is represented with pink van der Waals (vdW) spheres. The NA exhibiting head-tilt motion is highlighted in red during the dynamics, whereas the other HAs and NAs are colored with gray and dark gray, respectively.

**Movie S5.**

HA ectodomain-tilt motion. The ectodomain-tilt motion of a representative HA trimer extracted from the H1N1-Shan2009 whole-virion simulation is displayed. HA trimer is depicted with a transparent surface overlaid on top of a cartoon representation colored with a viridis palette according to the extent of ectodomain tilt. N-linked glycans and sequon asparagine are shown with yellow van der Waals (vdW) spheres.

**Movie S6.**

HA ectodomain-tilt motion. The ectodomain-tilt motion of an HA trimer is displayed together with the dynamics of the surrounding crowded environment of the H1N1-Shan2009 whole-virion. ~442 ns of MD are shown. Neuraminidase (NA) and hemagglutinin (HA) glycoproteins, and the M2 ion channels, are depicted with red, blue, and white surfaces, respectively. N-linked glycans are shown in yellow. The spherical lipid envelope is represented with pink van der Waals (vdW) spheres. The

HA exhibiting ectodomain-tilt motion is highlighted in blue during the dynamics, whereas the other HAs and NAs are colored with gray and dark gray, respectively.

**Movie S7.**

HA head breathing motion. The head breathing motion of a representative HA trimer extracted from the H1N1-Shan2009 whole-virion simulation is displayed from a side (left) and a top (right) viewpoint simultaneously. Each HA monomer within the trimer is depicted with a transparent surface overlaid on top of a cartoon representation colored with a viridis palette according to the extent of the respective head breathing. N-linked glycans and sequon asparagine are shown with yellow van der Waals (vdW) spheres.

**Movie S8.**

HA head breathing motion. The head breathing motion of an HA trimer is displayed together with the dynamics of the surrounding crowded environment of the H1N1-Shan2009 whole-virion. ~442 ns of MD are shown. Neuraminidase (NA) and hemagglutinin (HA) glycoproteins, and the M2 ion channels, are depicted with red, blue, and white surfaces, respectively. N-linked glycans are shown in yellow. The spherical lipid envelope is represented with pink van der Waals (vdW) spheres. The HA exhibiting head breathing motion is highlighted in blue during the dynamics, whereas the other HAs and NAs are colored with gray and dark gray, respectively.

**Movie S9.**

The dynamic interplay between glycoproteins. The glycoprotein interplay taking place during the simulations of the glycosylated whole-virion models of H1N1-Shan2009 (left) and H1N1-Mich2015 (right) is displayed. ~442 ns and ~425 ns of MD are shown, respectively. The glycoproteins are depicted with a surface colored according to the number of connections made by the respective glycoprotein at that frame (0 to 5, colored in viridis palette). Interacting glycans are highlighted with yellow vdW spheres, whereas non-interacting glycans with white VdW spheres.

**Movie S10.**

Glycoprotein macro-cluster formation. The formation of macro-cluster of glycoproteins taking place during the simulations of the glycosylated whole-virion models of H1N1-Shan2009 (left) and H1N1-Mich2015 (right) is displayed. ~442 ns and ~425 ns of MD are shown, respectively. The glycoproteins are depicted with a surface colored according to the size of the macro-cluster they are involved in at that frame (colored in *viridis* palette). All glycans are highlighted with white vdW spheres.

**Data S1.**

Jupyter-notebooks for the MSM analysis of NA head tilt, HA ectodomain tilt, and HA head breathing.

**Data S2.**

3D map visualization of Fab NDS.1 in complex with A/Darwin/9/2021 at 24.1 Å resolution. Density file (.mrc).
